# ClustoCell reveals cell states and their markers from single-cell transcriptomes

**DOI:** 10.64898/2026.08.20.746095

**Authors:** Adrian Salavaty, Momeneh Foroutan, Natalia Pretel Pretel, Jacob Egelberg, Ian A Parish, Nicholas D. Huntington, Himisha Beltran, Shahneen Sandhu, Ramyar Molania

## Abstract

Accurate identification of cell types and states is essential for reliable single-cell RNA-sequencing analyses, yet current methods remain sensitive to continuous biological states, data preprocessing choices, and reference selection. Here we present ClustoCell, a reference-free method that resolves cell identity using within-cell transcriptional architecture. By stratifying gene expression of each cell into high and medium tiers, ClustoCell constructs cell-cell similarity graphs that prioritize intrinsic expression structure over global variance. Across 450 datasets spanning over 24 million cells, ClustoCell recovered expert annotations with high concordance (92%). Benchmarked against state-of-the-art methods, ClustoCell identifies more stable and coherent cell types and states, avoids excessive partitioning of closely related cells, and improves the identification of cell type-specific markers. From transcriptional structure alone, ClustoCell resolves rare and transitional cell states, distinguishes malignant from non-malignant cells, and refines expert cell annotations. Applied to immunotherapy datasets, ClustoCell uncovered coordinated pre-treatment immune circuits linking T cell states to PD-1 responsiveness in a tumour-type-specific manner. ClustoCell provides an interpretable and scalable foundation for single-cell analysis and translational profiling.

## Introduction

Single-cell RNA sequencing (scRNA-seq) has transformed our ability to resolve cellular heterogeneity, enabling gene expression profiling at single-cell resolution across tissues, developmental trajectories, and disease states^1–5^. The scale of this technology has expanded dramatically, with tens of millions of cells now profiled across thousands of studies^6, 7^, capturing broad biological diversity and substantial cross-study variation. Accurate biological interpretation of scRNA-seq data highly depends on precise identification and annotation of cell types and states, yet this remains a persistently challenging and inconsistent step across single-cell analysis pipelines^8–11^. This challenge reflects the continuous nature of cell states, the lack of universal marker gene standards, and methodological differences across studies, which together limit reproducibility at the scale now required by modern single-cell atlases^10–16^.

Current cell type annotation strategies broadly fall into two categories, including marker-based and reference-based annotation, each with limitations^8, 16–19^. Marker-based annotation typically involves unsupervised clustering of the data followed by assignment of cluster identities using canonical marker genes. While this approach is interpretable and independent from external resources, it is sensitive to preprocessing steps, including normalization, feature selection, dimensionality reduction, clustering resolution, and differential expression-based marker detection^13, 20, 21^. These preprocessing choices can obscure biologically meaningful signals or amplify technical artifacts^12, 22^. Feature selection for dimensionality reduction commonly relies on highly variable genes (HVGs). However, high variance does not necessarily imply cell-type specificity^23, 24^ and may instead arise from technical variation. Such variance-driven artifacts can compromise cluster identification, and consequently cluster-level differential expression (DE) analysis may fail to capture true defining markers. In this situation, DE genes are often shared among closely related cell types^25^ or may reflect batch effects and other confounders^17,26^.

Reference-based methods, such as Azimuth^27^ and CellTypist^28^, address some of these challenges by mapping new datasets to curated references. However, their performance depends on reference quality, including label accuracy, coverage of relevant cell types and states, and compatibility with the query data in terms of platform and preprocessing^20, 29^. Further, different references frequently produce divergent annotations for the same dataset, introducing a source of variability that is difficult to diagnose or reconcile and impedes clinical translation^30^.

To overcome these limitations, we developed ClustoCell, a data-driven method that constructs cell–cell similarity graphs using cell markers, defined as genes showing high or moderate expression within a given cell, irrespective of their variability across the dataset^31, 32^. ClustoCell leverages highly expressed markers to capture core cell-type identity, and moderately expressed markers to resolve transitional states and underrepresented subtypes.

By defining cell similarity using cell-intrinsic expression strata rather than globally variable genes across cells, ClustoCell bypasses the need for initial dimensionality reduction and clustering, as well as dependence on external references. We benchmarked ClustoCell against widely used clustering and marker identification methods across diverse tissue, cancer, and immunotherapy datasets, demonstrating improvements in annotation accuracy, reproducibility, and biological interpretability.

## Results

### ClustoCell method

ClustoCell is a data-driven method that constructs cell–cell similarity graphs from cell-intrinsic expression strata to robustly identify biologically meaningful cell clusters and their defining gene markers from scRNA-seq data (Fig. 1). The pipeline proceeds through three major steps: gene filtering and transformation, cell–cell similarity graph construction, and cluster identification with marker ranking.

**Fig. 1:**
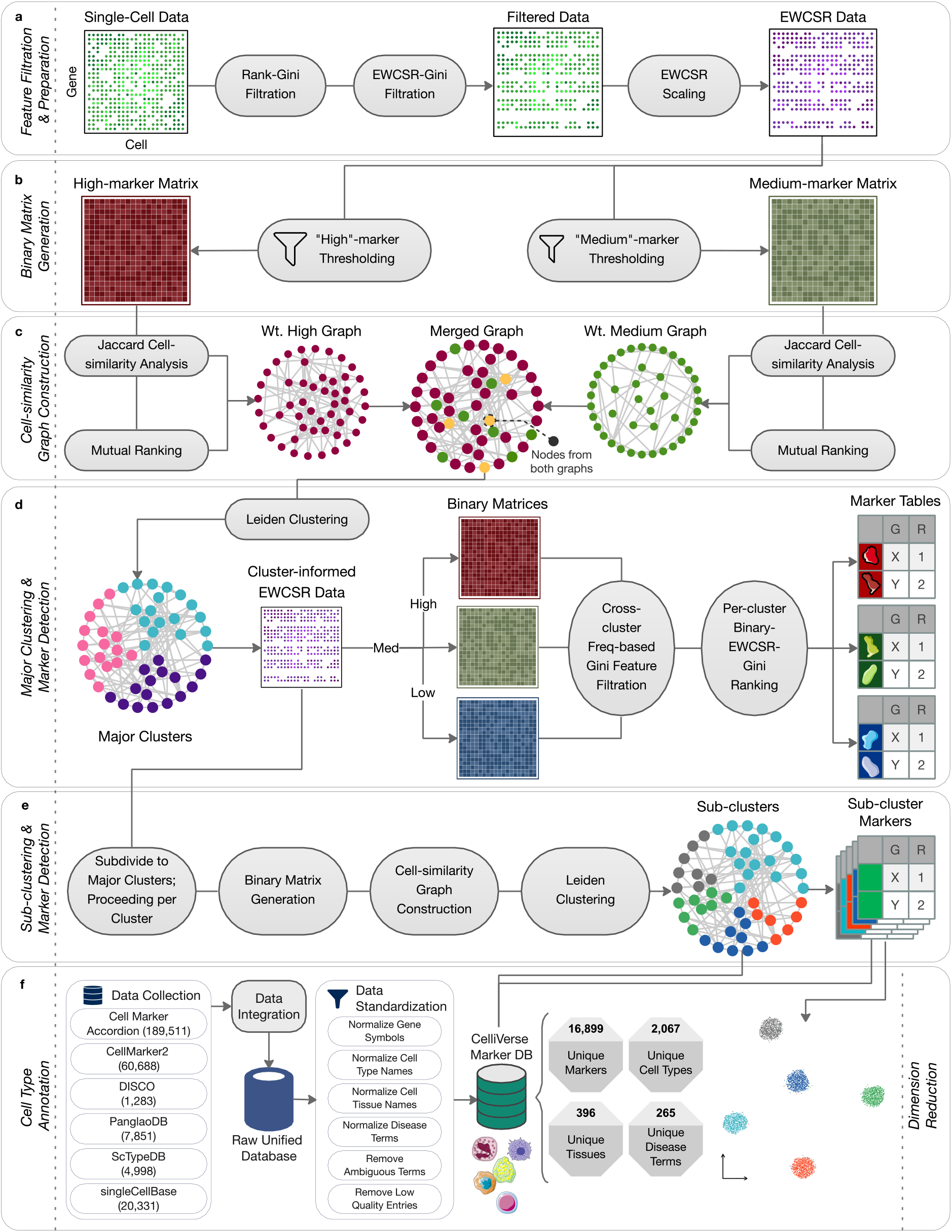
The ClustoCell framework for data-driven cell clustering and marker identification. ClustoCell processes scRNA-seq data through a multi-step pipeline to identify biologically meaningful cell clusters and their defining markers. **a,** Rank-based Gini filtration removes uniformly expressed genes, followed by EWCSR transformation and a subsequent Gini filtration to prioritize cell-intrinsic expression strata. **b,** Data-driven thresholds applied to a second round of EWCSR-transformed data yield binary matrices distinguishing high- and medium-expression strata within each cell; cells lacking high-expression genes are designated as quiescent and excluded. **c,** Jaccard similarity and mutual rank transformation are applied to each binary matrix to construct weighted cell–cell similarity graphs, which are merged into a unified graph. **d,** Leiden clustering defines major cell clusters; clusters below a size threshold are flagged as isolated. Per-cluster Gini assessment ranks positive, medium, and negative markers by purity. **e,** The pipeline is recursively applied within major clusters to resolve subclusters and subcluster-specific markers. **f,** Schematic overview of the CelliVerse Marker DB construction pipeline, showing integration of six public marker resources, standardization of gene symbols, cell type, tissue and disease nomenclature, and removal of inconsistent or low-confidence entries to generate a harmonized, high-confidence marker reference. ClustoCell-derived markers can be used for cell type annotation with CelliVerse Marker DB or any other marker-based annotation resource, as well as for cell type–informative dimensionality reduction.

The method begins by removing genes with uniform expression across cells (Fig. 1a). ClustoCell first applies a rank-based Gini filtration^33^ to discard genes with homogeneous expression, retaining only those with discriminative potential for cell type identification. Retained genes are then subjected to expression-weighted centered scaled rank (EWCSR) transformation, applied per cell by prioritizing genes relative to each cell’s own transcriptome, conferring robustness to global expression shifts, sequencing depth variation, and batch effects. A subsequent Gini filtration applied to the EWCSR-transformed values further refined the data by removing any remaining low-inequality features. A second round of EWCSR transformation is then applied to the expression data of the retained genes (Fig. 1a).

Global data-driven thresholds derived from these transformed values are used to discretize gene expression into “high”, “medium”, and “low” expression strata. This results in three binary matrices of identical dimensions, each representing the presence or absence of a given gene expression stratum, high, medium, or low, across all cells and genes (Fig. 1b). Cells lacking any high-expression stratum markers are designated as quiescent, reflecting resting or metabolically inactive states, and are excluded from downstream analysis^34–36^. Using the high and medium expression strata binary matrices, Jaccard-based cell–cell similarity is then computed, emphasizing shared expression stratum presence over precise expression ranks – an approach that more directly captures shared cell identity programs.

For each of the high and medium expression strata binary matrices, Jaccard similarity followed by mutual rank transformation is used to construct weighted cell–cell similarity graphs. The high- and medium-strata graphs are then merged into a single union graph that integrates complementary similarity information across both expression strata (Fig. 1c). Leiden clustering is then applied to this union graph to define major cell clusters, with clusters falling below a user-defined minimum size threshold designated as isolated cells (Fig. 1d).

Cluster marker identification is performed using complementary binary matrices representing high, medium, and low expression strata. Following removal of features with low inter-cluster inequality, ClustoCell uses per-cluster Gini assessment to rank positive, medium, and negative markers for each cluster, simultaneously quantifying marker purity (Fig. 1d). To resolve finer cellular structure, the full pipeline is recursively applied within each major cluster to identify subclusters and their associated markers (Fig. 1e).

Markers derived from both major clusters and subclusters enable direct cell type and cell state annotation using curated marker databases such as CelliVerse Marker DB, a comprehensive marker gene database developed in this work by integrating six major publicly available resources (Fig. 1f; Methods). These markers additionally serve as cell-type–informative features for enhanced dimensionality reduction and visualization, providing a biologically grounded alternative to HVG-based embeddings.

### ClustoCell yields cell-type-concordant clustering

To assess the ability of ClustoCell to identify cell-type-concordant clusters, we first applied it to 450 scRNA-seq datasets (224 unique studies) spanning diverse tissues and sequencing technologies, obtained from the 3CA^37^, TISCH2^38^, Shi *et al*. pan-cancer atlas^39^, and human Ensemble Cell Atlas (hECA v2.0)^40^ databases (Fig. 2a, Methods). Across these datasets, ClustoCell achieved a median concordance of 92% with original cell type annotations (Fig. 2b), demonstrating that ClustoCell accurately recovers underlying cell type structure across diverse single-cell datasets. We next benchmarked ClustoCell against nine widely used single-cell clustering and cell-labelling methods including Seurat^27^, Scanpy^41^, Scran^42^, scGPT^43^, Accordion^44^, scType^45^, SCINA^46^, AUCell^47^, and UCell^48^. Evaluation was performed across eight well-annotated scRNA-seq datasets: PBMC3K, PBMC10K, peritumoural normal liver (PNL), melanocyte-melanoma, basal cell carcinoma, colon cancer atlas, lung adenocarcinoma (LUAD) cell atlas, and the human breast cell atlas (Methods). These datasets comprised diverse immune, epithelial, and tumour cell types. Cell type annotations from the original studies, hereafter referred to as original manual annotations (OMA), served as ground-truth labels in this analysis.

**Fig. 2:**
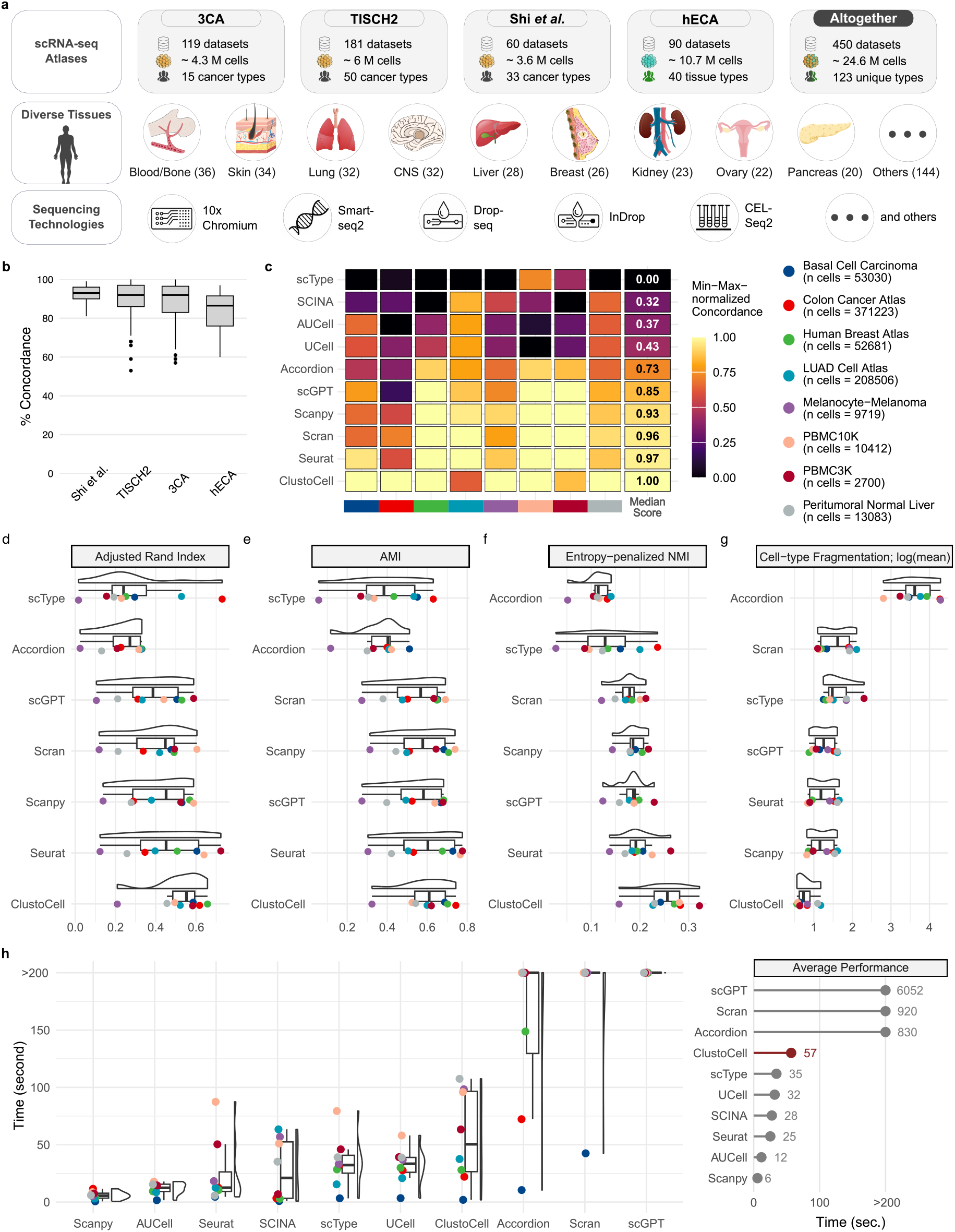
Benchmarking of ClustoCell clustering performance across single-cell RNA-seq atlases. ClustoCell clustering is compared with cell type labels from four single-cell RNA-seq (scRNA-seq) atlases and benchmarked against nine single-cell clustering and cell annotation methods across diverse tissue, cancer, and immune datasets including eight scRNA-seq datasets with expert-curated manual cell type annotations: basal cell carcinoma, colon cancer atlas, human breast cell atlas, LUAD cell atlas, melanocyte-melanoma, PBMC10K, PBMC3K, and PNL. **a,** Curated atlas-level single-cell RNA-seq datasets collected from four major public resources including 3CA, TISCH2, Shi *et al.* pan-cancer atlas, hECA v2.0, were used to evaluate ClustoCell across diverse human tissues, cancer types and sequencing platforms. **b,** Concordance between ClustoCell clustering and cell type labels from the four atlases. **c,** Heatmap of min-max normalised cluster purity – the degree to which each inferred cluster is dominated by a single annotated cell type – quantifying concordance between inferred clusters and original manual annotations (OMA), computed from contingency tables. **d,** Adjusted rand index (ARI) comparing inferred clusters to manual annotations across datasets. **e,** Adjusted mutual information (AMI), measuring information-theoretic agreement between clustering results and OMA while correcting for chance agreement. **f,** Normalised mutual information (NMI) penalized by cluster entropy, quantifying resolution-adjusted agreement. **g,** Cell type fragmentation scores (log-transformed mean fragmentation), measuring the extent to which annotated cell types are subdivided into multiple clusters, where lower fragmentation reflects higher cluster integrity — the representation of each cell type by a compact, interpretable cluster set. **h,** Computational runtime comparison across methods and datasets. Together, these metrics provide a comprehensive assessment of clustering performance by jointly evaluating biological concordance, resolution appropriateness, and cluster integrity beyond purity alone.

Initially, we assessed concordance between inferred cluster labels and OMA using cluster purity derived from contingency tables^49^ (Methods) (Fig. 2c; Supplementary Fig. 1–8). Across datasets, ClustoCell achieved the highest median concordance, with Seurat and Scran ranking second and third, respectively, while the remaining methods exhibited greater variability. These results indicated that ClustoCell robustly identifies clusters that capture dominant cell type-informative signals across diverse biological contexts.

However, partitioning the data into an excessively high number of small clusters may introduce bias in the concordance analysis^50^. To address this, we restricted comparisons to reference-free clustering methods, thereby avoiding bias arising from pre-specified cluster numbers imposed by input gene signatures. We quantified concordance by computing the Adjusted rand index (ARI) between OMA and clusters derived from unsupervised clustering. Across all datasets, ClustoCell consistently achieved the highest median and the lowest variation in ARI compared to the other methods (Fig. 2d). Seurat and Scanpy ranked second and third, respectively, while methods producing highly fragmented solutions showed markedly reduced ARI despite moderate purity-based concordance (Fig. 2c, d).

We next evaluated adjusted mutual information (AMI)^51^, a measure that corrects mutual information for chance agreement and reduces bias associated with differences in cluster structure. ClustoCell achieved the highest median AMI across datasets, followed by Seurat and scGPT, further confirming that its improved concordance with OMA was not driven by chance agreement or resolution inflation (Fig. 2e).

To further assess robustness to over-clustering, we computed normalised mutual information (NMI)^51^ and applied an entropy-based penalty to down-weight clustering solutions with inflated cluster-label entropy. Figure 2f shows that ClustoCell achieved the highest median score across datasets, reflecting accurate recovery of annotated structure without unnecessary fine-grained subdivision. Seurat and scGPT followed closely, whereas methods with inflated cluster entropy exhibited substantial score reduction, demonstrating that their apparent concordance with OMA was driven primarily by resolution inflation rather than accurate recovery of underlying biological structure.

Cell type fragmentation analysis (Methods) further showed that ClustoCell produced the lowest median fragmentation score across all datasets, indicating that the biological cell types were represented by compact and interpretable cluster sets (Fig. 2g). In contrast, over-clustering methods fragmented individual cell types into numerous small clusters, often spanning an order of magnitude more partitions. Scanpy and Seurat ranked second and third, respectively, but consistently exhibited higher fragmentation than ClustoCell. Together, these analyses demonstrate that ClustoCell achieved a favorable balance between biological resolution and cluster integrity.

Finally, we compared computational performance across datasets (Fig. 2h). Although ClustoCell was not the fastest method overall, its runtime scaled favorably with dataset size and remained comparable to Seurat across all datasets, while substantially outperforming Scran and Accordion on large atlases. These results indicate that ClustoCell combines robust biological performance with practical computational efficiency.

### ClustoCell improves cluster marker identification and cell-type separation

To evaluate ClustoCell’s performance in cluster marker identification, we benchmarked it against nine widely used methods: Seurat^27^, MAST^52^, DESeq2^53^, Wilcox-Limma^27^, ROC-based approaches, logistic regression, Scran, Scanpy, and NSForest^32^. Benchmarking was performed across three different datasets, including PBMC3K, PNL, and the prostate cell atlas, covering a wide variety of cell types (Methods).

For each dataset, OMA were considered as individual clusters, and the top 10 ranked markers per cluster were identified using each method. Overlap between identified markers and corresponding entries in the CelliVerse Marker DB (Fig. 1f) was then quantified. ClustoCell achieved the highest median min-max normalised overlap score across all datasets, consistently outperforming all other methods. Scanpy and NSForest frequently ranked among the top four performers, while logistic regression (LR) and Wilcox-Limma fell in the lowest-ranked quartile (Fig. 3a–c).

**Fig. 3:**
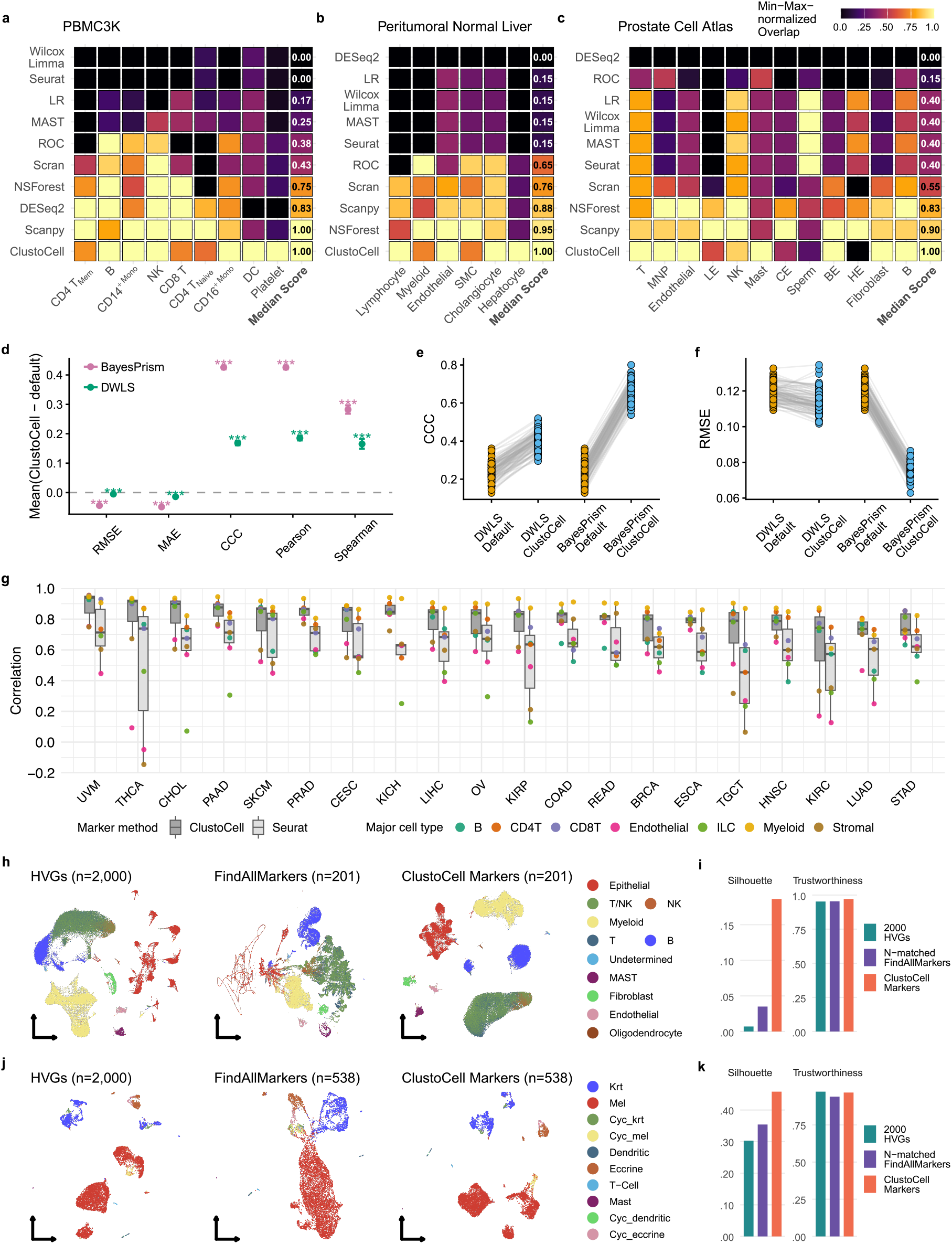
ClustoCell improves cluster marker identification and cell type separation across diverse scRNA-seq datasets. **a–c,** Heatmaps of overlap scores derived from confusion matrices comparing curated cell type markers from the CelliVerse Marker DB against the top ten ranked markers identified by each marker identification method for PBMC3K (**a**), peritumoural normal liver (**b**), and the prostate cell atlas (**c**), with scores normalised to enable cross-method comparison. **d–f**, Benchmarking of cell type deconvolution using published default marker selection strategies and ClustoCell-derived markers across two widely used deconvolution methods, DWLS and BayesPrism. **d**, Forest plot summarising the mean paired differences (ClustoCell − default) across five deconvolution performance metrics, including Pearson correlation, Spearman correlation, concordance correlation coefficient (CCC), root mean squared error (RMSE), and mean absolute error (MAE), with bootstrap 95% confidence intervals. **e,f**, Paired comparisons of concordance correlation coefficient (CCC) (**e**) and root mean squared error (RMSE) (**f**) across 100 matched pseudobulk samples generated from an independent PBMC dataset (PBMC10K). Each pair of connected points represents deconvolution of the same pseudobulk sample using the published default marker selection strategy or ClustoCell-derived markers while keeping the underlying deconvolution algorithm unchanged. **g**, Comparison of ClustoCell- and Seurat-derived cell-type signatures across 20 TCGA cancer types. Box plots show the distributions of Spearman correlation coefficients between cancer-specific cell-type signature scores and ESTIMATE scores for signatures generated from the top 30 markers identified by each method. Colored points indicate individual major cell types. **h,i**, LUAD Cell Atlas. **h**, UMAP embeddings generated using Seurat’s default dimensionality reduction pipeline with three alternative feature sets: 2,000 highly variable genes (HVGs), ClustoCell-derived top markers, and an N-matched set of Seurat *FindAllMarkers* genes (left to right). **i**, Quantification of cluster separability (mean silhouette width) and embedding trustworthiness for each feature set. **j,k**, Melanocyte and melanoma dataset. **j**, UMAP embeddings generated using HVGs, ClustoCell-derived markers, or Seurat-derived markers. **k**, Corresponding silhouette and trustworthiness metrics.

To assess whether ClustoCell-derived markers improve cell type deconvolution for bulk RNA-seq data, we benchmarked two widely used methods, DWLS^54^ and BayesPrism^55^, using either their default marker selection strategy or the top 10 ranked ClustoCell-derived markers across the three benchmark datasets shown in Figures 3a–c (Methods). Across independent query datasets spanning immune, liver and prostate tissues, BayesPrism generally achieved higher concordance with ground-truth cell type proportions than DWLS, and replacing its default marker set with ClustoCell-derived markers consistently further improved deconvolution performance. Performance gains for DWLS were more dataset dependent, indicating that the impact of marker selection varies between deconvolution frameworks (Fig. 3d–f; Supplementary Fig. 9). Together, these results demonstrate that ClustoCell-derived markers provide robust reference signatures for accurate downstream cell type deconvolution across diverse tissue contexts.

Next, we assessed the performance of ClustoCell in identifying cancer-specific markers for major cell types within the tumour microenvironment (TME) (Methods). For this analysis, we first identified cancer-specific TME cell-type markers from the pan-cancer single-cell atlas of Shi *et al*. pan-cancer atlas^39^ using both Seurat and ClustoCell. These marker sets were then used to estimate the abundance of major TME cell types in the corresponding TCGA RNA-seq cohorts. We subsequently evaluated the associations between these estimates and tumour purity, calculated using the ESTIMATE^56^ method. Across 20 TCGA cancer types, ClustoCell-derived signatures showed stronger concordance with ESTIMATE scores than Seurat-derived signatures (Fig. 3g). The median Spearman correlation was 0.80 for ClustoCell compared with 0.60 for Seurat, and ClustoCell produced the stronger correlation in 141 of 157 matched cancer-cell-type comparisons (median paired difference in ρ = 0.120; paired Wilcoxon *P* = 2.39 × 10^−21^). Immune and stromal signatures were generally positively correlated with ESTIMATE scores, consistent with greater non-malignant cell content. These results indicate that ClustoCell-derived markers capture biologically relevant tumour-microenvironment signals with greater concordance to ESTIMATE-derived stromal and immune content than Seurat-derived markers.

Further, we evaluated whether ClustoCell-derived markers improve cell type separation in low-dimensional embeddings, *e.g.,* UMAP. In this analysis, we applied Seurat’s default dimensionality reduction workflow (Methods) to two independent datasets spanning cancer and immune contexts, using three alternative gene sets: highly variable genes identified by the VST approach, top ClustoCell-derived markers of author-defined cell types, and a matched number of markers identified by Seurat’s *FindAllMarkers* function. In both datasets, PCA embeddings generated using ClustoCell-derived markers consistently achieved higher cluster separability, as quantified by mean silhouette width, while maintaining comparable levels of embedding trustworthiness compared with the other two gene sets (Fig. 3h–k). The improvement was most pronounced when there was strong transcriptional similarity between related cell types, where ClustoCell-derived markers yielded clearer inter-cluster boundaries and reduced overlap in low-dimensional space. Together, these results demonstrate that ClustoCell consistently identifies biologically relevant cluster markers across diverse cell types.

### ClustoCell is robust to normalization choices and global batch effects

In previous analyses, ClustoCell demonstrated robust performance when it was applied to raw count data without any normalization. This arises from its rank-based EWCSR transformation and the construction of cell–cell similarity graphs from binary-transformed EWCSR matrices, which emphasize within-cell gene ordering rather than absolute expression magnitudes.

To systematically assess the impact of normalization on ClustoCell’s performance, we applied it to PBMC3K data processed by five normalization strategies: raw count, Seurat log normalization, Scran log normalization, centered log-ratio (CLR; margin = 2), and SCTransform. ClustoCell performed similarly across raw counts and Seurat-, Scran-, and CLR-normalised data. However, performance was modestly reduced on SCTransform-normalised data, with concordance decreasing to 93%, consistent with SCTransform reshaping within-cell gene rank order through variance modeling and regression of technical effects (Extended Data Fig. 1a). Notably, concordance with OMA was preserved after normalization, with only a small decrease observed for SCTransform-normalised data (Extended Data Fig. 1a).

We next evaluated how robust ClustoCell is to data splitting and batch effects. First, the PBMC3K dataset was randomly partitioned into two samples and analyzed independently. In both samples, ClustoCell-derived clusters exhibited near-perfect concordance (98–99%) with clusters obtained from the full dataset and retained comparable concordance with OMA (93–95% versus 94% for the full dataset; Extended Data Figure 1b,c). Second, we analyzed a merged PBMC3K–PBMC10K dataset, in which standard Seurat normalization and dimensionality reduction revealed strong batch-driven separation (Extended Data Fig. 1d). Despite this, ClustoCell clustering on the merged data showed 98% concordance with clusters obtained from the individual datasets analyzed separately (Extended Data Fig. 1e), indicating that ClustoCell largely decouples biological structure from batch-specific technical variation without requiring explicit batch correction. Finally, on a Splatter-simulated dataset comprising five biological groups across two batches, ClustoCell perfectly recovered the ground-truth groups (100% concordance; Extended Data Fig. 1f,g), whereas the standard Seurat pipeline produced batch-stratified clusters even at very low resolution, yielding only 62% concordance with the true groups (Extended Data Fig. 1h). This result further demonstrates that ClustoCell preferentially captures biological signals over batch effects, even under conditions of strong simulated batch confounding. Together, these results demonstrate that ClustoCell is highly robust to most normalization strategies and batch effects, with reduced performance confined to methods such as SCTransform that model and regress technical variation in ways that inadvertently perturb within-cell gene rank structure, the primary signal leveraged by ClustoCell.

We next examined whether ClustoCell-derived markers improve cell type separation in low-dimensional embeddings of multi-batch datasets (Methods). Applied to pancreatic islet (Panc8) and IFN-β–stimulated PBMC datasets, ClustoCell-derived markers exhibited robust resistance to batch effects, grouping cells by biological identity rather than technical batch, whereas HVG- and Seurat-marker–based embeddings showed substantial batch-driven separation (Extended Data Fig. 1i–l). Seurat-derived markers performed least favorably in this setting, reflecting the sensitivity of differential expression–based marker selection to batch-associated variance. Together, these results indicate that ClustoCell-derived markers encode cell-type–specific structure that is both discriminative and robust, enabling more biologically informative dimensionality reduction.

### ClustoCell is scalable to large scRNA-seq datasets

ClustoCell scales efficiently to large single-cell atlases by combining data sketching with cluster label transfer to the full dataset. Applied to three large scRNA-seq atlases of normal and cancer tissues including colon cancer, LUAD, and breast tissue, ClustoCell was run on sketched subsets of 20,000 cells, followed by label transfer to all remaining cells. Despite operating on a small representative subset, this strategy yielded cluster assignments with consistently high concordance with OMA (>90%) across all three datasets (Extended Data Fig. 2).

Sketching and label transfer are natively integrated into the ClustoCell framework, allowing users to specify sketch size and label transfer strategy directly. This enables clustering, annotation propagation, and marker identification to be performed in a single, streamlined workflow, maintaining annotation fidelity while substantially reducing computational cost – making ClustoCell well suited for routine analysis of large-scale single-cell atlases.

### ClustoCell surpasses expert manual annotation of PBMC cell types

To assess ClustoCell’s ability to recover major immune cell populations, we applied it to the PBMC3K dataset, a highly curated and widely used peripheral blood mononuclear cell reference with expert-derived annotations (Extended Data Fig. 3a). Several annotated populations reflect shared developmental origins, including CD8^+^ T cells and natural killer (NK) cells derived from common lymphoid progenitors^57^, naïve and memory CD4^+^ T cells arising from CD4^+^ T cell precursors, and FCGR3A^+^ monocytes, CD14^+^ monocytes and dendritic cells originating from the myeloid lineage via the monocyte–dendritic cell progenitor pathway (Extended Data Fig. 3b).

ClustoCell identified major clusters with 94% concordance with OMA, and only 88 discordant cells across the full dataset after excluding unannotated cells (Extended Data Fig. 3c). To investigate these discrepancies, we examined all discordant subsets containing at least 10 cells, together with cells manually annotated as “NA” (not assigned) (Extended Data Fig. 3d,e). For each subset, ClustoCell-derived marker programs were computed, and the top ten markers were identified (Extended Data Fig. 3f).

Cell types were then assigned to each subset using an integrated marker-based scoring approach, which combines information on markers overlapping with the CelliVerse Marker DB and their expression patterns within each subset (Methods). Using this integrated score, ClustoCell correctly reassigned all discordant subsets and NA cells to their appropriate cell types, outperforming the original expert annotations (Extended Data Fig. 3g). The sole exception was a subset of 14 cells (from C3) originally annotated as NA, which were assigned as NK cells rather than CD4^+^ T cells. These cells exhibited low marker purity — with top NK markers expressed in only 21% of cells in the subset — and a markedly lower combined score (22) compared to all other subsets (maximum 2,723), suggesting reduced data quality or ambiguous transcriptional state within this subset rather than a limitation of the ClustoCell framework.

### ClustoCell resolves cell subtypes and cell states

We assessed ClustoCell’s ability to resolve biologically meaningful cell types and cell states across normal and cancer conditions, using the PBMC3K dataset and the malignant compartment of a LUAD single-cell atlas.

In the PBMC3K dataset, ClustoCell identified five major lineage-level clusters (Fig. 4a) that collectively captured all nine OMA-annotated cell types (Extended Data Fig. 3c), with 12 subclusters nested within them. Examination of cell type fractions per subcluster revealed that most annotated cell types mapped predominantly to a single subcluster, whereas B cells, CD14^+^ monocytes and platelets were each partitioned into two distinct subclusters (Fig. 4b), suggesting finer-grained stratification by ClustoCell within these lineages. ClustoCell-derived marker programs for these subclusters indicated distinct functional cellular states. To validate this, we curated subtype- and state-specific gene signatures for each lineage (Supplementary Data 1) and computed per-cell module scores for the corresponding subclusters. Signature-based analyses showed that subclusters belonging to the same OMA cell type were transcriptionally distinct, and that the finer-grained B cell, CD14^+^ monocyte and platelet subclusters each corresponded to a different functional state (Fig. 4c–e).

**Fig. 4:**
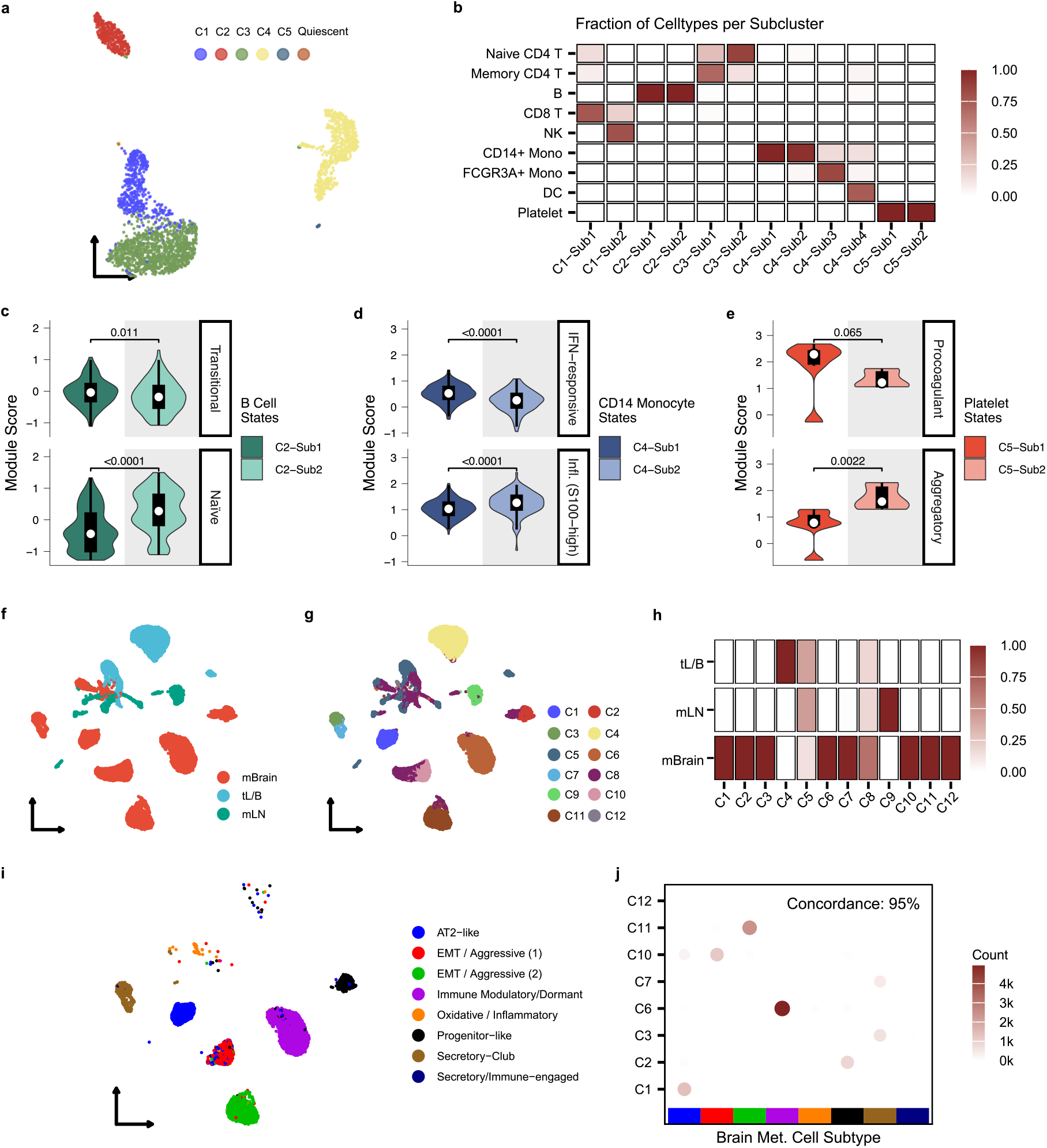
ClustoCell resolves biologically coherent cell types and states in normal and disease contexts. **a**, UMAP of the PBMC3K dataset colored by ClustoCell clusters. **b**, Confusion matrix showing the fraction of OMA within each ClustoCell subcluster. 12 subclusters are nested within these 5 major clusters. **c–e**, Violin plots of module scores for lineage-specific subtype or state signatures in PBMC3K: B cell states (**c**), CD14^+^ monocyte states (**d**), and platelet states (**e**), shown for their respective ClustoCell subclusters. **f**, UMAP of malignant cells from the LUAD atlas colored by anatomical origin. **g**, UMAP of LUAD malignant cells colored by ClustoCell cluster assignment. **h**, Confusion matrix showing the fraction of each anatomical origin within each LUAD malignant cell cluster. **i**, UMAP of LUAD brain metastatic cells colored by assigned malignant cell subtypes or states. **j**, Concordance dot plot comparing ClustoCell brain metastasis–related clusters with malignant cell subtypes/states.

We next applied ClustoCell to the malignant compartment of the LUAD atlas, comprising primary tumour cells as well as lymph node and brain metastases (Fig. 4f). ClustoCell identified 12 malignant cell clusters (Fig. 4g), with each cluster predominantly corresponding to a single anatomical origin, except for clusters C5 and C8, and eight clusters enriched for brain metastases (Fig. 4h). Marker programs for clusters C5 and C8 revealed a shared signature indicative of functional plasticity and metastatic potential, consistent with a hybrid EMT and stress-adapted state, marked by expression of *STMN1*, *S100A4*, *GLO1*, *SEPP1*, *IFI6*, and *IFI27*. Focusing on brain metastatic cells, we defined eight malignant cell subtype or state signatures informed by ClustoCell marker programs (Supplementary Data 1) and assigned a subtype or state to each cell using module scoring. This analysis revealed two closely related but functionally distinct epithelial–mesenchymal transition (EMT)/aggressive states sharing a core identity defined by *S100A4*, *S100P* and *STMN1*, but diverging in programs associated with invasive growth and angiogenesis (HGF/MET-driven) versus hyperproliferation (EGFR/keratin-driven) (Fig. 4i). Concordance analysis demonstrated 95% agreement between ClustoCell clusters and brain metastasis cell subtypes or states, indicating high accuracy in resolving malignant cell states at metastatic sites (Fig. 4j).

### ClustoCell identifies malignant cells in cancer scRNA-seq data

To assess whether ClustoCell can distinguish malignant cells from non-malignant cells, we applied it to 12 scRNA-seq datasets spanning 10 different cancer types (Methods). OMA in the source studies were generated through expert curation and, where applicable, informed by canonical marker expression and copy-number inference. Across datasets, ClustoCell distinguished malignant from non-malignant populations with excellent agreement with OMA (Supplementary Fig. 10-21), achieving an average concordance of 96% (Fig. 5a), despite using no marker genes, reference labels or copy-number information. These results indicate that ClustoCell resolves malignant cells directly from transcriptional structure.

**Fig. 5:**
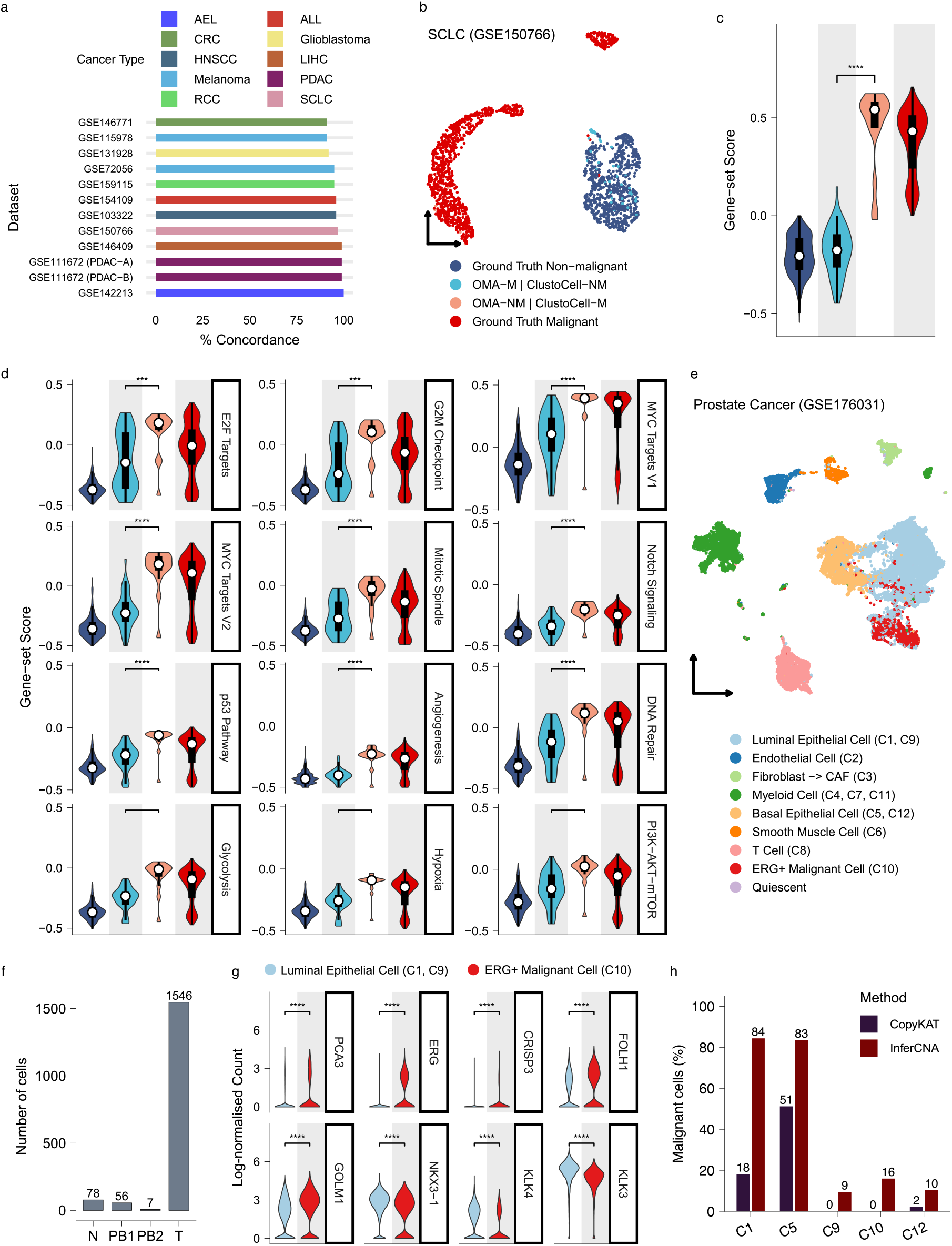
ClustoCell discriminates malignant and non-malignant cells. **a,** Concordance between ClustoCell-derived clusters and OMA of malignant and non-malignant cell populations across 12 scRNA-seq datasets spanning 10 cancer types including Acute Erythroid Leukaemia (AEL); Acute Lymphoblastic Leukaemia (ALL); Colorectal Cancer (CRC); Glioblastoma; Head and Neck Squamous Cell Carcinoma (HNSCC); Liver Hepatocellular Carcinoma (LIHC); Melanoma; Pancreatic ductal adenocarcinoma (PDAC); Renal cell carcinoma (RCC); Small Cell Lung Cancer (SCLC). Each dataset was clustered into two major groups using ClustoCell, and concordance was calculated relative to OMA. **b,** UMAP of the SCLC dataset colored by concordant malignant and non-malignant cell populations (considered as ground truth sets) as well as discordant subsets, cells annotated as non-malignant by OMA but assigned to the malignant ClustoCell cluster (OMA-NM | ClustoCell-M) or vice versa (OMA-M | ClustoCell-NM). **c,** Distribution of differential expression (DE)-based malignancy scores across SCLC ground-truth (concordantly grouped into the same cluster) and discordant cell populations. **d,** Distribution of MsigDB cancer hallmark gene sets scores across SCLC ground-truth (concordant) and discordant cell populations. **e**, UMAP of the prostate cancer scRNA-seq dataset (GSE176031) showing the 12 ClustoCell-derived clusters annotated into nine cell types based on cluster marker genes. **f**, Distribution of sample sources contributing to the ERG^+^ malignant cell cluster (C10), with 92% of cells originating from tumour samples. PB1 and PB2 denote cells derived from the first and second prostate biopsy patient groups, respectively; T denotes cells from radical prostatectomy tumour specimens; and N denotes cells from matched normal prostate tissue collected during radical prostatectomy. **g**, Expression of representative prostate cancer marker genes in the ERG positive malignant cell cluster compared with luminal epithelial cells. **h**, Percentage of cells classified as malignant/aneuploid by CopyKAT and InferCNA across epithelial ClustoCell clusters, including the ERG positive malignant cell cluster (C10).

Discrepancies between ClustoCell and OMA may reflect transcriptionally mixed cells, transitional states between normal and malignant identities, or epithelial-adjacent non-malignant populations that cannot be confidently classified. To investigate these discordant cells, we focused on the Small Cell Lung Cancer (SCLC) dataset, a particularly informative model given its highly distinct neuroendocrine transcriptional identity, strong proliferative programs, large copy number burden, and minimal overlap between malignant and normal lung epithelial states (Fig. 5b). Scoring discordant cells using DEGs between cells concordantly classified as malignant and non-malignant by both ClustoCell and OMA demonstrated higher accuracy of ClustoCell assignment relative to OMA in this dataset (Fig. 5c). Consistently, scoring cells against 12 SCLC-relevant cancer hallmark programs further confirmed the superior accuracy of ClustoCell in discriminating malignant from non-malignant cells (Fig. 5d).

Further, given that prostate cancer is among the most transcriptome-altered and copy-number-driven malignancies, making it a particularly suitable benchmark for CNA-based malignancy inference^58, 59^, we next compared ClustoCell with CopyKAT^60^ and InferCNA on a prostate cancer scRNA-seq dataset comprising biopsy and radical prostatectomy tumour samples together with matched normal tissues. ClustoCell resolved 12 clusters that were annotated into nine distinct cell types based on their marker genes (Fig. 5e), including an ERG^+^ malignant cell cluster derived predominantly from tumour samples (92%; Fig. 5f) that exhibited a distinct ERG-associated malignant expression profile, with higher expression of ERG-associated and cancer markers and lower expression of several mature luminal differentiation markers than luminal epithelial clusters (Fig. 5g). In contrast, InferCNA and CopyKAT classified 16% and none of the cells from this malignant cluster as aneuploid/malignant, respectively (Fig. 5h).

Together, these analyses show that ClustoCell achieves high global concordance with expert annotations while providing biologically coherent reinterpretation of discordant cells, in several cases supporting refined discrimination between malignant and non-malignant states beyond what manual annotation captured.

### ClustoCell identifies a conserved effector–exhaustion axis predictive of PD-1–driven clonal expansion in breast cancer

To evaluate ClustoCell’s ability to resolve T cell states relevant to immune checkpoint therapy, we analyzed the breast cancer cohort of Bassez *et al*.^61^, which includes detailed T/NK annotations and TCR-based clonal expansion measurements. Applying ClustoCell to this dataset, we resolved the T/NK compartment into 19 subclusters that provided substantially finer granularity than the author-defined categories (Fig. 6a, b). Broad phenotypes such as CD4_EX and CD8_EX (exhausted T cells), and CD4_EM (effector-memory T cells) segregated into molecularly coherent states capturing intermediate/effector-like exhausted CD8 cells, activated/early-effector-like exhausted CD8 cells with interferon-licensed HLA-DR^+^ activation, CXCL13^+^ T peripheral helper (TPH)-like CD4 exhausted cells, and cytotoxic effector-memory CD4 differentiation. Subcluster-annotation concordance confirmed that ClustoCell recovered these states at higher resolution than the original annotations (Fig. 6b), with multiple subclusters differentially enriched with respect to TCR-derived expansion phenotypes (Fig. 6c).

**Fig. 6:**
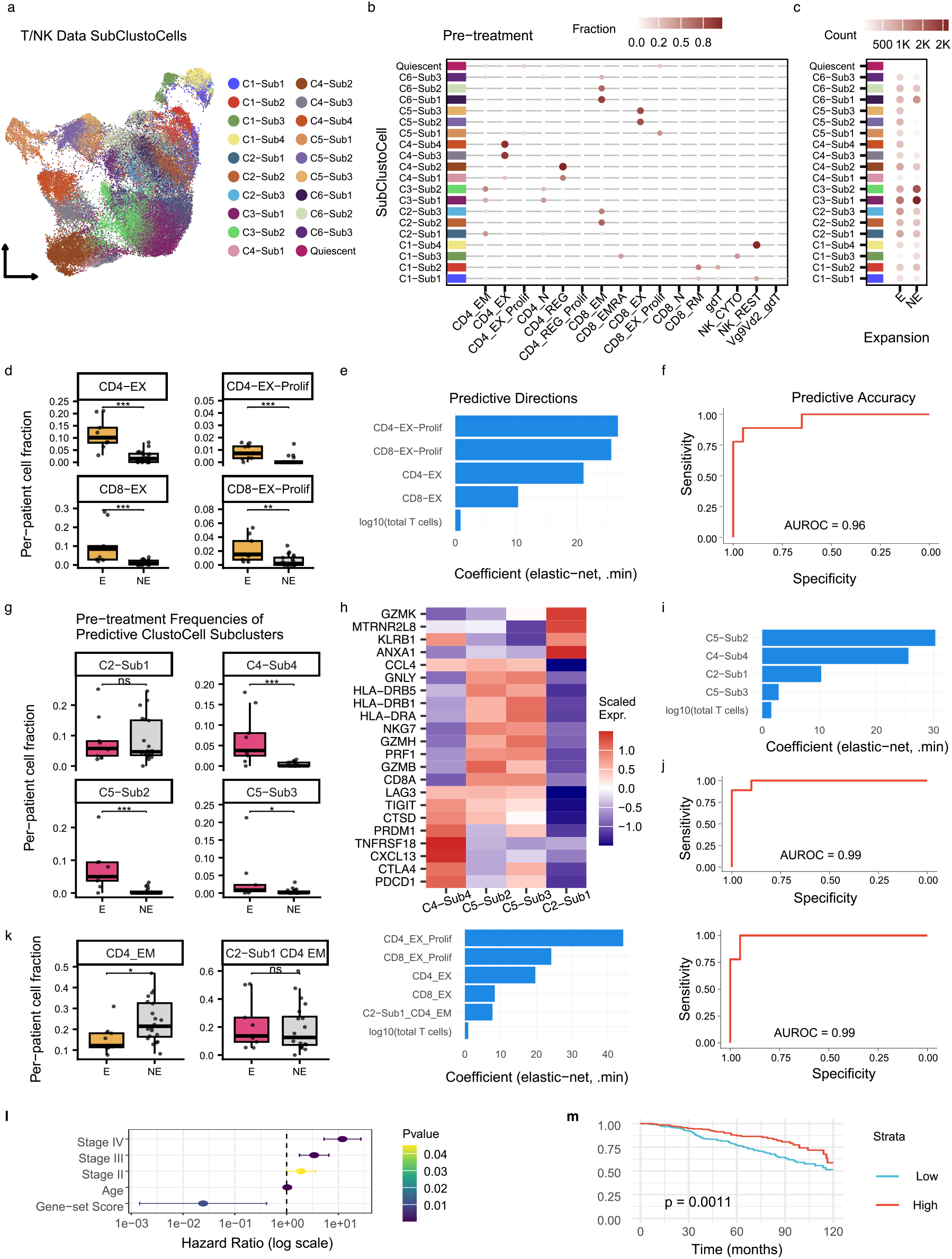
Predictive T-cell states comprising the Coordinated Effector–Exhaustion Circuit in pre-treatment breast cancer. **a**, UMAP of the T/NK compartment annotated by 19 ClustoCell subclusters. **b**, Fractions of author-defined T/NK annotations within each subcluster in pre-treatment biopsies. **c**, Mapping of subclusters to clonal expansion status. **d**, Per-patient frequencies of author-defined predictive cell types in Expander and Non-expander cases. **e**, Elastic-net coefficients indicating predictive direction across author-defined predictive cell types and T yield. **f**, ROC curve for author-defined predictive states (AUROC = 0.96). **g**, Per-patient frequencies of predictive ClustoCell subclusters C5-Sub3, C4-Sub4, C5-Sub2, and C2-Sub1. **h**, Scaled expression heatmap of ClustoCell-derived defining markers for the four expansion-enriched predictive subclusters: C4-Sub4 (CXCL13^+^ TPH-like CD4 exhausted), C5-Sub2 (HLA-DR^+^ NR4A1^+^ activated / early-effector-like exhausted CD8), C5-Sub3 (HLA-DR^+^ CX3CR1^+^ intermediate effector-like exhausted CD8), and C2-Sub1 (IL7R^+^ GZMK^+^ CD4 effector-memory). An expanded panel of canonical lineage and activation/exhaustion markers across the same four subclusters is shown in Supplementary Figure 22. **i**, Elastic-net coefficients indicating predictive direction across the four subclusters and T yield. **j**, ROC curve for prediction using ClustoCell subclusters (AUROC = 0.99). **k**, Per-patient frequencies of CD4_EM and the CD4_EM subset within C2-Sub1, corresponding elastic-net coefficients, and ROC curve for prediction (AUROC = 0.99). **l**, Forest plot showing univariable Cox proportional hazards model for overall survival in TCGA breast cancer (BRCA) patients using a composite gene set score derived from Singscore (upregulated genes and CD4 effector-memory (EM) signature (*CD4*, *CD40LG*, *IL7R*, *ITGB1*, *ANXA1*) minus downregulated genes). **m**, Kaplan–Meier overall survival curves for TCGA BRCA patients stratified by median split of the composite Singscore (upregulated genes and CD4 EM signature minus downregulated genes).

As an initial benchmark, we re-evaluated the author-defined predictors – CD4_EX, CD4_EX_prolif, CD8_EX, and CD8_EX_prolif – which constituted the strongest signals in the original study. Consistent with prior findings, these categories were markedly enriched in expander patients in the pre-treatment setting (Fig. 6d) and together with log_10_ T yielded a high-performing elastic-net model (AUROC = 0.96; Fig. 6e,f), confirming that our re-analysis faithfully recapitulates the principal predictive signals of the original study and provides a robust baseline for evaluating higher-resolution state definitions.

We next asked whether operating directly at the resolution of ClustoCell-defined subclusters, rather than author-defined cell type categories, could improve predictive performance. Classifying subclusters by their association with TCR-derived expansion labels (Fig. 6c; Methods), the four most significantly enriched subclusters, C5-Sub3, C4-Sub4, C5-Sub2, and C2-Sub1, were enriched among expander patients before therapy (Fig. 6g), with marker performance revealing strong, non-redundant signatures (Fig. 6h; Supplementary Fig. 22): C5-Sub2 corresponded to an HLA-DR^+^, NR4A1^+^, interferon-licensed activated/early-effector-like exhausted CD8 population, and C5-Sub3 to an HLA-DR^+^ CX3CR1^+^ intermediate effector-like exhausted CD8 state – both mapping to the author-defined CD8_EX category; C4-Sub4 to a CXCL13^+^ CD4 TPH/TEX-like program mapping to a subset of author-defined CD4_EX; and C2-Sub1 to a transcriptionally distinct CD4 effector-memory state representing a subset of author-defined CD4_EM. Together, these states define a coordinated pre-treatment transcriptional module spanning effector-memory and effector-exhaustion arms across both CD4 and CD8 lineages, which we term the Coordinated Effector–Exhaustion Circuit. Pseudotime analysis confirmed that these states are embedded within continuous lineage-specific differentiation trajectories rather than representing isolated transcriptional endpoints: within the CD8 compartment, C6-Sub1 was ordered upstream of C5-Sub2 and C5-Sub3 along an effector-memory to HLA-DR^+^ effector-exhausted continuum (Extended Data Fig. 4a; Supplementary Data 1), while within the CD4 lineage, C2-Sub1 was positioned upstream of the CXCL13^+^ TPH-like CD4 exhausted state C4-Sub4 (Extended Data Fig. 4b), together summarised as a coordinated pre-treatment transcriptional circuit (Extended Data Fig. 4c). A consistent circuit architecture was identified in the BCC dataset, where ClustoCell resolved an analogous set of expansion-associated states organized along coherent lineage trajectories (Supplementary Note 1; Supplementary Fig. 23,24).

The combined fractions of these four subclusters, together with log_10_ T yield, correlated positively with expansion (Fig. 6i) and enabled highly accurate discrimination of expander versus non-expander cases (AUROC = 0.99; Fig. 6j), outperforming models based on author-defined cell types alone (Fig. 6f).

To determine whether ClustoCell-defined subcluster resolution could augment the author’s existing predictive model without replacing its canonical predictors, we examined the contribution of transcriptionally refined subpopulations to predictive performance. The author-defined CD4_EM category, while not among the original predictors, maps across multiple ClustoCell subclusters, including the expansion-enriched C2-Sub1 and the non-expander-enriched C3-Sub1 and C3-Sub2, suggesting that its heterogeneity dilutes its predictive signal at coarse resolution. Including C2-Sub1 CD4_EM alongside the author-defined exhausted and proliferative predictors retained high predictive performance (AUROC = 0.99; Fig. 6k), demonstrating that subcluster-level resolution can unmask expansion-relevant signals obscured by coarse annotation. These results demonstrate that ClustoCell-defined subcluster resolution can both replace and augment author-defined predictors, motivating further investigation of the biological and prognostic basis of the key discriminating population, C2-Sub1.

To validate the biological specificity of the C2-Sub1 CD4 EM population, we contrasted it with a transcriptionally related but non-expander-enriched CD4 EM subcluster, C3-Sub2 (Fig. 6b,c). Differential expression analysis identified prominent upregulation of cytotoxic lymphocyte genes, such as *GZMA*, *GZMK*, CCL4 and CCL5, in C2-Sub1 relative to C3-Sub2 (Supplementary Data 1). To assess whether this transcriptional distinction carries prognostic relevance, we scored TCGA breast cancer samples against a composite program comprising these DEGs augmented with CD4 EM lineage-anchoring markers (*CD4*, *CD40LG*, *IL7R*, *ITGB1*, *ANXA1*) to mitigate potential NK or CD8 T cell confounding. This score was significantly associated with improved overall survival (Fig. 6l,m), with a consistent hazard ratio direction after multivariate adjustment (HR = 0.05; P = 0.08), confirming the prognostic association of this transcriptionally distinct CD4 EM state. The C2-Sub1 score also correlated positively with CIBERSORT-inferred, M1 macrophages, resting memory CD4 T cells, CD8 T cells and regulatory T cells, indicating that this CD4 effector-memory program is embedded within a broader immune-infiltrated, T-cell-inflamed tumour microenvironment rather than reflecting an isolated cell population (Supplementary Fig. 25).

## Discussion

ClustoCell introduces a unified, data-driven framework for single-cell analysis that integrates feature filtration, clustering, and marker discovery, without relying on global variance detection, external references, or preferred gene signatures. ClustoCell operates on cell-intrinsic expression strata, a data-driven, per-cell stratification of gene expression into high-, medium-, and low-expression tiers. This prioritizes within-cell transcriptional identity over population-level statistical patterns, enabling accurate recovery of cell types across diverse tissues, disease contexts, and experimental platforms without explicit batch correction or normalization standardization.

Existing annotation strategies either depend on globally variable genes that conflate technical and biological variation, or on curated references that limit applicability to well-characterized cell types. ClustoCell requires only expression data as input, no reference atlas, no predefined marker genes, no gene signatures, making it applicable to any cell identity regardless of prior characterization. Applied without modification across 224 independent datasets spanning diverse tissues, cancers, and platforms, it recovered original cell type annotations with high concordance, demonstrating generalizability at scale. In benchmarking against CelliVerse Marker DB, a harmonized marker reference integrating six public resources developed here, ClustoCell prioritized cell-type–specific markers more effectively than differential expression or feature-ranking approaches. In clustering evaluations across eight diverse datasets, ClustoCell consistently recovered annotated cell types as stable, unfragmented populations, achieving superior concordance, ARI, AMI, and fragmentation scores compared to leading methods, including the pre-trained single-cell foundation model scGPT, while maintaining practical computational efficiency and requiring no pretraining.

The robust performance of ClustoCell stems from rank-based, binary-transformed features that decouple biological from technical variation, recovering ground-truth biological groups with perfect accuracy while standard pipelines produce batch-stratified clusters. ClustoCell-derived markers further improve dimensionality reduction over HVG or DE-based feature sets, and sketch-based label propagation extends the framework to atlases of millions of cells with minimal loss of resolution.

In curated immune datasets, ClustoCell went beyond reproducing expert annotations by resolving ambiguous and unannotated cells into lineage-consistent identities. Across 10 tumour types, ClustoCell distinguished malignant from non-malignant cells with 96% average concordance from transcriptional structure alone, without copy-number inference. Notably, in prostate cancer, a canonical copy-number-driven malignancy, it identified a tumour-derived ERG positive malignant population that dedicated copy-number-based callers failed to robustly detect, underscoring the sensitivity of transcriptional-structure-based malignancy discrimination.

Applied to immunotherapy cohorts, ClustoCell delineates a mechanistically cohesive pre-treatment immune circuit, the Coordinated Effector–Exhaustion Circuit, integrating inhibitory checkpoint expression, interferon-stimulated gene licensing, cytotoxic effector programs, and TPH-driven CXCL13-mediated recruitment into a single framework linking T cell states to PD-1 response. ClustoCell further revealed that canonical cell type categories dilute predictive signal by collapsing transcriptionally heterogeneous subpopulations with opposing enrichment directions, a resolution-dependent limitation that subcluster-level analysis can overcome. The prognostic relevance of this refined resolution was confirmed in the TCGA breast cancer data, where a C2-Sub1-derived score was associated with improved survival and co-enriched with a broader immune-engaged microenvironment; a consistent circuit architecture in basal cell carcinoma demonstrated cross-tumour generalizability.

Some limitations merit consideration. ClustoCell’s performance is modestly reduced on data processed with SCTransform, which reshapes the within-cell gene rank order it leverages; raw or log-normalised counts are preferable in such cases. Additionally, the predictive immune circuits identified here were derived from relatively small immunotherapy cohorts and require validation in larger datasets to establish clinical utility, and the biological interpretation of the associated differentiation trajectories is further constrained by the absence of tumour-draining lymph nodes from these datasets.

Collectively, ClustoCell provides a reference-free, biologically grounded framework, unifying clustering, marker discovery, and annotation. By linking transcriptional resolution to biological and clinical relevance, it offers a generalizable tool for dissecting cellular heterogeneity, refining disease phenotypes, and informing precision interventions in immunotherapy and beyond.

## Methods

### ClustoCell model architecture

ClustoCell operates on an input scRNA-seq expression matrix *X* ∈ *R^d×n^*, where *d* denotes the number of genes and *n* the number of cells.

#### Rank-based and EWCSR-based feature filtration

For each cell *j*, genes are ranked according to expression levels, yielding rank values *R_i,j_* for gene *i*. For each gene, inequality of rank distributions across cells is quantified using the Gini coefficient:

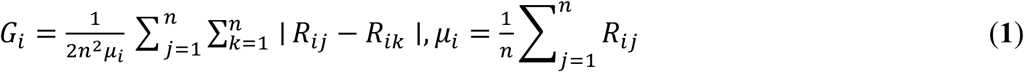

Genes with *G_i_* below a threshold *θ_gini_* are filtered out.

Next, an expression-weighted centered scaled rank (EWCSR) transformation is applied. For each cell *j*, let *G*_j_ denote the set of expressed genes, 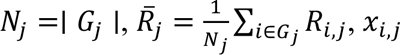 the raw expression count of gene *i* in cell *j*, and *L*_%_ the library size. EWCSR values are computed as:

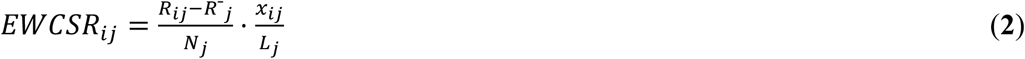

After EWCSR transformation (Eq. 2), ClustoCell defines expression-state thresholds using the empirical distributions of positive and negative EWCSR values separately, rather than assuming a symmetric or unified distribution.

Let

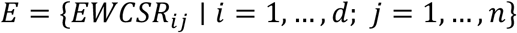

denote the set of all EWCSR values across features and cells. We partition this set into positive and negative components:

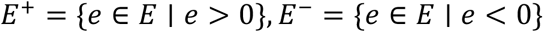

The high-expression EWCSR threshold is defined as the upper α-quantile of the positive EWCSR distribution:

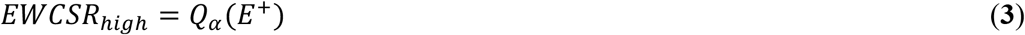

Similarly, the low-expression EWCSR threshold is defined as the upper β-quantile of the absolute negative tail (equivalently, the β-quantile of the negative EWCSR distribution):

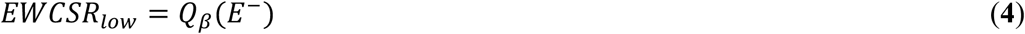

where *Q*(⋅) denotes the empirical quantile function, and α and β are user-defined parameters controlling the stringency of high and low expression states, respectively.

This asymmetric, tail-specific thresholding ensures that highly overexpressed and strongly under-expressed genes are identified relative to their respective empirical distributions, rather than assuming symmetry around zero. This design preserves sensitivity to cell-type–specific activation and repression programs while remaining robust to global shifts in expression scale and sparsity.

Binary matrices encoding high and medium expression states are constructed as:

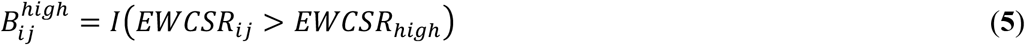

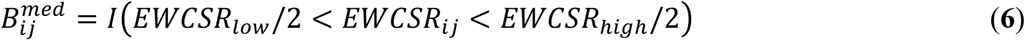

Binary features are further filtered using the Gini coefficient, which for binary data simplifies to 1 – *p_i_*, where *p_i_* is the proportion of cells exhibiting the expression state. These prioritized features are then utilized to derive a finalized EWCSR sparse matrix from raw count data (Eq. 2). Subsequently, the *EWCSR_high_* and *EWCSR_low_*: thresholds are recalibrated against this finalized matrix according to Eq. 3 and Eq. 4, respectively. This iterative adjustment enables the high-fidelity reconstruction of binary matrices that encode high and medium expression states.

#### Cell–cell similarity graph construction

For each binary matrix, Jaccard similarity between cells *j* and *k* is computed as:

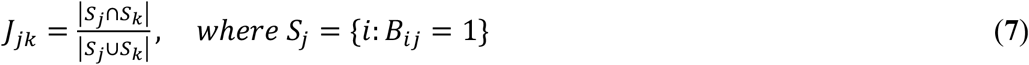

Mutual rank is then defined as:

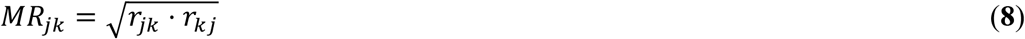

where *r*_jk_ denotes the rank of *J*_jk_ among the Jaccard similarities in row *j* (*i.e.*, cell *j*’s similarities to all other cells), and *r*_)%_ denotes the corresponding rank of *J*_jk_ in row *k*. Adjacency matrices are obtained by thresholding mutual ranks:

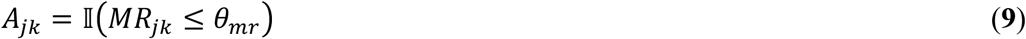

where *θ*_mr_ is a mutual rank threshold (defaulting to 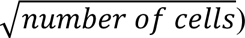 that controls the sparsity of the similarity graph.

High- and medium-marker graphs are constructed using these adjacency matrices and Jaccard weights and merged into a single union graph by summing shared edge weights. Leiden clustering is applied to identify major clusters; clusters smaller than the user-defined minimum size threshold (*θ*_C8DEF=B_, default = 5) are labelled as isolated cells.

#### Marker identification and subclustering

A low-expression binary matrix is defined as:

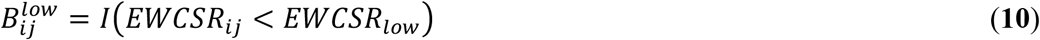

Let *k* denote the total number of clusters identified by Leiden clustering and let C_k_ denote the set of cells assigned to cluster *k*, with *n*_k_ = |C_k_|.

For each binary matrix, cluster-level feature frequencies are computed:

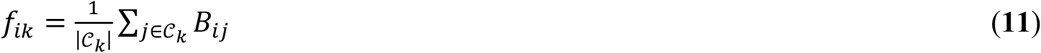

and features with low inequality across clusters are removed using a cluster-level Gini coefficient:

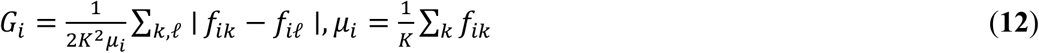

Within each cluster, per-feature impurity is quantified using a binary Gini formulation:

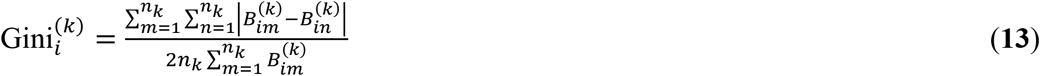

Within which simplifies to 1 – *p*_i_, where *p*_i_ is the proportion of cells exhibiting the expression state in cluster *k*. Features are ranked in ascending order of 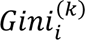 to prioritize positive, medium, and negative markers separately.

The full pipeline is recursively applied within each major cluster to identify subclusters and subcluster-specific markers.

### CelliVerse Marker Database

We created CelliVerse Marker DB, a comprehensive and high-quality database of marker genes for annotation of cell type and cell state. CelliVerse Marker DB integrated and harmonized data from six major cell type marker gene resources, including Cell Marker Accordion v1.0^44^, CellMarker2^62^, DISCO^63^, PanglaoDB^64^, ScTypeDB^45^, and singleCellBase^65^. For each database, the information on marker gene symbols, corresponding cell types, tissue sources, and disease conditions was extracted. The initial database comprised 284,662 marker genes, each associated with a specific cell type, 22,764 unique marker genes, 4,585 cell types, 496 tissues, and 335 disease conditions (Supplementary Data 1).

We then applied a systematic standardization pipeline to unify gene symbols, cell type nomenclature, tissue labels, and disease condition terms across databases, while removing inconsistent or low-confidence entries. Gene symbols were harmonized using the latest HGNC (HUGO Gene Nomenclature Committee) reference^66^. Also, cell type, tissue, and disease condition labels underwent lexical normalization and semantic harmonization. This involved standardizing nomenclature by resolving discrepancies in capitalization, consolidating abbreviated and elaborated forms, and merging synonymous identities into a unified vocabulary. Entries containing ambiguous, low-quality, or inconsistent annotations were excluded. The resulting database comprises 275,367 entries, 16,899 marker genes, 2,067 cell types, 396 tissues, and 265 disease conditions (Supplementary Data 2). This refinement establishes the CelliVerse Marker DB as a unified, interoperable, high-confidence resource and the largest reference for cell-type and cell state marker identification (Fig. 1f).

### Concordance analysis

Throughout this study, we evaluated the agreement between two sets of cell labels (for example, cluster labels generated by ClustoCell and the manual cell type annotations provided in the original study) using contingency tables.

For each comparison, we constructed a contingency table that summarised the overlap between the two label sets. To ensure unbiased comparison across methods with differing intrinsic resolutions, contingency tables were oriented adaptively: when the number of inferred clusters exceeded the number of annotated cell types, clusters were placed on rows and annotations on columns, reflecting fine-grained subdivisions of broader cell types; when fewer clusters were inferred, annotations were placed on rows and clusters on columns, reflecting coarser groupings. This symmetric formulation avoided imposing a fixed one-to-one correspondence between clusters and annotations. Concordance was calculated as the sum of the largest value in each row of the table, divided by the total number of cells. This value was reported as a percentage.

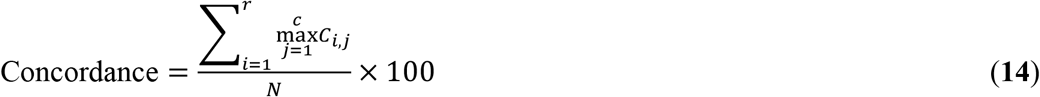

where *r* is the number of rows, *c* is the number of columns, and *N* is the total number of cells. *C*_i,j_ represents the count of instances assigned to row *i* and column *j*.

### Concordance analysis against public scRNA-seq atlases

We evaluated ClustoCell across all scRNA-seq datasets from the 3CA^37^, TISCH2^38^, Shi *et al*. pan-cancer atlas^39^, and human Ensemble Cell Atlas (hECA v2.0)^40^ databases, all of which were retrieved in May 2026. These resources differ in their preprocessing pipelines, quality control procedures, and cell type annotation strategies, providing a diverse collection of datasets for evaluating clustering performance across heterogeneous data processing and annotation frameworks. Datasets containing only a single annotated cell type were excluded. For the hECA atlas, datasets with highly granular cell type annotations were converted to broader, higher-level annotations because distinguishing closely related cell types and states is inherently sensitive, and overly granular annotations can artificially suppress concordance scores. Study accession identifiers were cross-referenced across databases to identify overlapping studies. Within the Shi *et al*. pan-cancer atlas, cohorts comprising multiple cancer types were split into individual cancer-specific datasets to match the study-level granularity used by 3CA and TISCH2. This resulted in a final collection of 450 datasets (224 unique studies). All datasets were converted to Seurat objects. Datasets originating from multiple source databases were intentionally retained, allowing direct comparison of ClustoCell performance on the same biological study processed using different preprocessing and annotation pipelines.

ClustoCell was applied to each dataset using default parameters. For datasets containing more than 40,000 cells, sketch-based clustering was enabled. The appropriate input assay layer and log1p transformation were determined dynamically by inspecting the expression matrix using a series of statistical criteria, including the proportion of integer-valued entries, the maximum expression value, the distribution of values above predefined thresholds, and the mean expression. This procedure distinguished raw count matrices from normalised TPM/CPM or log-transformed expression matrices, allowing raw count datasets to be processed using the *counts* layer with internal log1p transformation, whereas normalised datasets were analysed directly using the *data* layer without additional transformation. To avoid concordance suppression arising from the lower granularity of major ClustoCell clusters relative to the reference cell-type annotations, concordance was calculated using ClustoCell sub-clusters against the database-derived annotations (Eq. 14).

### Benchmarking ClustoCell against existing clustering methods

Next, we benchmarked ClustoCell against nine widely used single-cell clustering and cell-labelling methods, including Seurat^27^, Scanpy^41^, Scran^42^, scGPT^43^, Accordion^44^, scType^45^, SCINA^46^, AUCell^47^, and UCell^48^, across eight scRNA-seq datasets with expert-curated manual cell type annotations: basal cell carcinoma, colon cancer atlas, human breast cell atlas, LUAD cell atlas, melanocyte-melanoma, PBMC10K, PBMC3K, and PNL. For the concordance analysis between clusters and expert annotations, the manual T and NK labels were merged into a single T/NK category in the LUAD cell atlas. This consolidation was necessary because the manually defined T/NK super-cluster contains 64,403 cells, markedly larger than the isolated T (21,971) and NK (4,853) subsets. ClustoCell further subdivided this broad lymphoid compartment into two coherent subclusters, one enriched for T-cell features and the other for NK-cell features, capturing fine-grained structure rather than diverging from the manual taxonomy. Treating T and NK as separate manual categories would therefore inflate discordance by artificially fragmenting what, biologically and numerically, is a single dominant compartment. Collapsing these labels into a unified T/NK group provides a size-balanced and biologically faithful basis for evaluating agreement between unsupervised clustering methods and expert annotations.

All methods were applied using their recommended workflows and default parameters unless otherwise specified. To ensure comparability, we utilized the same number of principal components (PCs) for all methods, determined via the elbow method. The sole exception was Scran, which employs its own internal procedure for defining the required number of PCs. For scGPT, cell embeddings were generated using the recommended pretrained whole-human model and were subsequently used as input to the Scanpy KNN graph construction and Leiden clustering workflow. We used specific configurations for scType to ensure biological relevance. For the PBMC datasets, the scType reference tissue was restricted to ‘Immune system’; for all other datasets, we specified the corresponding tissue type (*e.g.*, ‘Liver’ for the PNL dataset) alongside ‘Immune system’. In cases where an exact match was unavailable in the scTypeDB, the most biologically proximal tissue was selected (*e.g.*, ‘Brain’ for the melanocyte–melanoma dataset) in conjunction with ‘Immune system’. For datasets exceeding 50,000 cells, we applied Seurat’s Uniform sketching procedure to subsample 20,000 cells, as several methods required >10 minutes to process full datasets. All methods performed clustering on the sketched data, after which cluster labels were transferred to the full dataset using Seurat’s *ProjectData* function to ensure consistent evaluation across methods. For ClustoCell, labels were transferred using its internal EWCSR-based procedure (ewcsr-cor) for datasets containing fewer than 100,000 cells to maximize accuracy. In this approach, cluster labels identified by ClustoCell in the sketched dataset are transferred to the full dataset by computing the correlation between the gene EWCSR profile of each cell in the full (non-sketched) expression matrix and the EWCSR centroid representing each cluster derived from the sketched data. Each cell is assigned the label of the cluster whose centroid shows the highest correlation. For larger datasets, the same label-transferring method applied to other clustering methods (*i.e*., *ProjectData*) was used to balance accuracy and computational efficiency. Cluster purity was assessed by calculating the concordance of clustering results with OMA (Eq. 14). Cluster purity quantifies how strongly each cluster is composed of cells belonging to a single ground truth class, with values ranging from 0 to 1 where higher scores reflect better clustering performance^49^.

For SCINA^46^, AUCell^47^, and UCell^48^, which require predefined gene signatures, we supplied dataset-specific marker gene sets corresponding to the manually annotated cell types (Supplementary Data 3). As a result, these methods were constrained to produce exactly one cluster per annotated cell type. For datasets containing multiple annotated subtypes of the same lineage (*e.g.*, Tumor_1 and Tumor_2, or B_cells_1 and B_cells_2 in the basal cell carcinoma dataset), we refined signature gene sets using Seurat’s *FindMarkers* function to identify subtype-specific markers.

To complement purity-based concordance, we evaluated biological resolution and cluster integrity using three complementary metrics. **Adjusted rand index (ARI)** was computed using the *adjustedRandIndex* function from the mclust R package^67^ v6.1.2. ARI quantifies agreement between clustering results and OMA while correcting for chance, with values ranging from 0 (random agreement) to 1 (perfect agreement). **Adjusted mutual information (AMI)** was computed using the *AMI* function from the aricode R package v1.1.0. AMI quantifies information-theoretic agreement between clustering results and OMA while correcting for chance agreement. **Normalised mutual information (NMI)** was computed using symmetric normalization of mutual information between cluster labels and annotations. To penalize over-clustering, we adjusted NMI by cluster entropy, defined as:

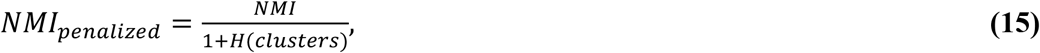

where entropy (*H*) was computed from the empirical distribution of cluster labels.

Cell type fragmentation^68^ was quantified by counting, for each annotated cell type, the number of clusters in which it constituted at least 10% of the cells. Mean fragmentation scores were computed across cell types, with lower values indicating greater biological coherence.

### Benchmarking of cluster marker identification

Benchmarking of cluster marker identification was conducted using nine methods across three scRNA-seq datasets including PBMC3K, PNL, and the prostate cell atlas. The methods included Seurat (Wilcoxon rank-sum test)^27^, MAST^52^, DESeq2^53^, Wilcox-Limma (Seurat’s *wilcox_limma* method)^27^, ROC analysis, LR (Logistic Regression), Scran^42^, Scanpy^41^, and NSForest^32^. Each method was implemented using its default parameters, as specified by the original developers. The datasets span diverse tissue contexts and cellular compositions. They include high-resolution immune subsets as well as parenchymal and stromal populations including hepatocytes, cholangiocytes, and contractile smooth muscle cells. Cell type annotations provided by each study (referred to as OMA) were treated as the ground-truth clusters for marker identification benchmarking. To reduce ambiguity arising from overlapping transcriptional programs within the hepatic myeloid compartment, macrophage populations in the PNL dataset were aggregated into a unified Myeloid cluster prior to analysis.

All datasets were normalised using Seurat *LogNormalize* method before applying the marker identification methods. For each expert-annotated cell type, gene markers were identified and ranked by each method, and the top ten ranked genes were retained. Performance was evaluated by quantifying the overlap between the top high-ranking genes and the corresponding marker sets in the CelliVerse Marker DB. For cross-dataset comparison, overlap counts were min–max normalised within each cell type to the range [0,1], where 1 indicates the highest-performing method.

Benchmarking of cell type deconvolution was performed using two widely used reference-based deconvolution methods, DWLS^54^ and BayesPrism^55^. For each method, two reference signatures were evaluated: (i) the published default marker selection strategy implemented by the original method, and (ii) an equivalent reference signature constructed using the top 10 ClustoCell-derived markers while keeping all other components of the deconvolution pipeline unchanged. Reference signatures were generated independently from the PBMC3K, peritumoural normal liver (PNL) and Prostate Cell Atlas scRNA-seq datasets. Evaluation was performed using independent query scRNA-seq datasets comprising PBMC10K, GSE115469 for liver and GSE185344 for prostate. For each query dataset, one hundred pseudobulk samples were generated by randomly sampling 1,000 cells per sample and summing their raw gene expression counts. Ground-truth cell type proportions were calculated directly from the sampled cells. Deconvolution performance was assessed by comparing estimated and true cell type proportions using Pearson correlation, Spearman correlation, concordance correlation coefficient (CCC), root mean squared error (RMSE) and mean absolute error (MAE). Performance differences between the default and ClustoCell-derived marker signatures were evaluated using paired statistical tests across matched pseudobulk samples, and bootstrap confidence intervals were calculated for the mean paired differences.

We used the pan-cancer single-cell RNA-sequencing dataset from Shi *et al*. to identify cancer-specific markers for individual tumour microenvironment (TME) cell types including B cells, CD4 T cells, CD8 T cells, endothelial cells, innate lymphoid cells, myeloid cells, and stromal cells. For each cell type within each cancer type, the top 30 marker genes were independently identified using Seurat and ClustoCell. These cancer- and cell-type-specific marker sets were subsequently used to calculate gene-set scores in the corresponding TCGA cancer type. The gene-set scores were calculated using the singscore R package (v1.26.0).

To find markers with Seurat, a Seurat object was created from the raw counts and normalised by *LogNormalize* with a scale factor of 10,000. Positive markers were detected using *FindAllMarkers* with the Wilcoxon rank-sum test, a minimum detection fraction of 0.01, a log2 fold-change threshold of 0.1, and sparse-matrix processing. Markers were required to have a positive average log2 fold change and a Benjamini-Hochberg-adjusted *P*-value below 0.05. Within each group, genes were ordered by decreasing average log2 fold change, followed by adjusted *P*-value and gene symbol; duplicate genes were removed, and up to 30 markers were retained.

TCGA RNA-sequencing data were downloaded using the TCGAbiolinks R package (v2.34.0), and upper-quartile-normalised FPKM (FPKM-UQ) expression values were used for downstream analyses. For each TCGA project, raw unstranded counts and upper-quartile-normalised fragments per kilobase per million mapped reads (FPKM-UQ) were obtained from the harmonized SummarizedExperiment object. Gene filtering was determined from the aggregated raw-count matrix using the default settings of *filterByExpr* from the edgeR R package (v4.4.2); the resulting filter was then applied to the FPKM-UQ matrix. Across projects, 15,294–20,587 gene symbols remained after filtering.

Tumour purity for individual TCGA samples was estimated using the ESTIMATE^56^ method from the R package tidyestimate (v1.1.1). For each cancer type, Spearman’s correlation coefficients were calculated between the cancer-specific cell-type scores and estimated tumour purity to assess the relationship between inferred cell-type abundance and tumour content. Analyses were performed separately for marker sets identified using Seurat and ClustoCell.

### Dimensionality reduction using ClustoCell-derived markers

For each dataset, ClustoCell was applied using its default parameters to identify cluster-specific positive markers, treating the original manually annotated cell types as clusters. Top-ranked positive markers were extracted for each cluster and aggregated across clusters to form the ClustoCell-derived feature set. The number of markers selected per cluster was set to *N* = 25 for the LUAD cell atlas, pancreatic islet (Panc8), and IFN-β PBMC datasets, and *N* = 90 for the melanocyte and melanoma dataset, which required a larger marker set due to high transcriptional similarity among multiple cell types. Because markers may be shared across clusters, the total number of unique ClustoCell-derived markers was generally less than *N* multiplied by the number of clusters.

For comparison, Seurat-derived marker sets were generated using the *FindAllMarkers* function (Wilcoxon rank-sum test). Only significant markers (adjusted *P* < 0.05) were retained and ranked by log_2_ fold change. The top *N* markers were selected to match the total number of features prioritized by ClustoCell, ensuring balanced representation across clusters and enabling direct benchmarking.

Dimensionality reduction was performed using Seurat’s standard workflow, performing PCA (*RunPCA*) followed by UMAP (*RunUMAP*) on all feature sets. Also, the same number of principal components was used for PCA, UMAP visualization, silhouette score calculation, and trustworthiness estimation within each dataset.

Cluster separability in PCA space was quantified using mean silhouette width (R package bluster v1.18). Trustworthiness^69^ between PCA and UMAP embeddings was computed using the scikit-learn implementation (Python 3.12.8, scikit-learn v1.7.0) on a random subset of 10,000 cells per dataset to ensure computational efficiency, following established best practices in large-scale single-cell benchmarking.

### Assessment of robustness to normalization and batch effects

ClustoCell was applied to all datasets using default parameters, except that *log1p = FALSE* was used when data were already normalised and log-transformed. For normalization robustness analyses, the PBMC3K dataset was analyzed in raw form and after normalization using Seurat, Scran, CLR normalization (margin = 2), and SCTransform^70^.

To assess robustness to subsampling and batch partitioning, PBMC3K cells were randomly assigned to two samples using random sampling with replacement, and ClustoCell was applied independently to each sample and to the full dataset. Concordance was computed between clusters obtained from subsampled and full data, and relative to OMA.

For cross-dataset batch analysis, PBMC3K and PBMC10K datasets were merged and analyzed jointly and separately. Concordance was computed between ClustoCell clusters derived from merged and individual datasets.

Simulation-based batch robustness was evaluated using a Splatter-generated dataset comprising two batches and five biological groups. Simulation parameters included batch-specific expression shifts and group-specific differential expression (*batchCells = c(500, 800), nGenes = 2000, group.prob = c(0.1, 0.2, 0.3, 0.05, 0.35), de.prob = 0.6, batch.facLoc = 0.3, batch.facScale = 0.3*). ClustoCell was applied with *leiden_resolution = 0.6* to recover five major clusters. For comparison, Seurat clustering was performed with resolution set to 0.001. Overall concordance was computed as the sum of the maximum cell count in each row of the contingency table divided by the total number of cells (Eq. 14).

### Scalable implementation of ClustoCell

For each dataset, we selected a representative subset of 20,000 cells using Seurat’s *Uniform* sketching procedure^71^. ClustoCell was performed on this subset to obtain clusters. The resulting cluster labels were then transferred to the full dataset. Similar to benchmarking of cell clustering, for datasets containing fewer than 100,000 cells, label transfer was performed using ClustoCell’s EWCSR-based method (*ewcsr-cor*) to maximize accuracy.

For larger datasets, label transfer was performed using the *clustoCell_TransferLabel* function from the celliverse R package with *method = “count-project”* and *dims = 20* to improve computational efficiency while maintaining accuracy. The *count-project* method implements Seurat’s *ProjectData* pipeline. Briefly, the sketched dataset is added as a new assay to the full dataset object. Both assays are processed using a standard workflow including *LogNormalize* normalization (*NormalizeData*), identification of 2,000 variable features using the VST method (*FindVariableFeatures*), scaling (*ScaleData*), and principal component analysis (*RunPCA*). Label transfer is then performed using *ProjectData* with *sketched.reduction = “pca”*, *normalization.method = “LogNormalize”*, and *dims = 1:20*. The reference labels (*refdata*) correspond to the ClustoCell cluster assignments of the cells in the sketched dataset. These choices were made to ensure a consistent analysis strategy across all datasets and sections of the study. After label transfer, concordance between ClustoCell-derived clusters and OMA was calculated on the full dataset (Eq. 14).

### ClustoCell’s clusters of PBMC3K data

ClustoCell was applied to the PBMC3K dataset using default parameters to identify major cell clusters. OMA were used as an external reference for concordance analysis, quantified as described in Eq. 14.

The identity of discordant cell subsets — those assigned to a different cluster than their OMA label — was evaluated by comparing the top 10 ClustoCell-derived markers per subset against curated cell type markers in the CelliVerse Marker DB. For each subset-cell type pair, two intermediate quantities were computed. The **purity-adjusted marker count** reflects the net contribution of overlapping markers: purity-weighted counts of markers concordantly identified (where both ClustoCell and CelliVerse Marker DB agree on the positive or negative status) for the candidate cell type are summed, whereas purity-weighted counts of discordant markers (where the tool and reference provide opposing annotations) are subtracted. **The purity-adjusted marker occurrence** reflects concordance across the independent resources compiled in CelliVerse Marker DB: purity-weighted occurrence counts of concordant positive and negative markers are summed, and occurrences corresponding to conflicting annotations are subtracted. Both quantities can therefore take positive or negative values, depending on the balance of supporting and conflicting evidence.

To integrate these two signals into a single metric while preserving the directionality of evidence, a sign-preserving combined score was defined as:

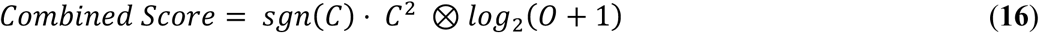

where *C* denotes the purity-adjusted marker count and *O* denotes the purity-adjusted marker occurrence. The operator ⊗ indicates sign-consistent multiplication, such that the magnitude of the product is given by the standard product of the two terms, while the sign is negative also when both inputs are negative, thereby preventing artificial sign inversion when both quantities are conflicting (*i.e.*, negative).

Each cell subset was assigned to the cell type with the highest combined score. Average marker purity values reported in Extended Data Figure. 3g represent the mean expression purity of cell type-relevant overlapping markers within each subset.

### Identification of cell identity in healthy and disease conditions

ClustoCell was applied to the PBMC3K dataset and to the malignant compartment of the LUAD atlas using default parameters. For PBMC3K, subclusters were identified within major lineage-level clusters. For LUAD, clustering was performed on malignant cells spanning primary tumours and metastatic sites.

Subtype- and state-specific gene signatures were curated based on ClustoCell-derived marker programs and prior biological knowledge (Supplementary Data 1). Per-cell module scores were computed using Seurat’s *AddModuleScore* function. Cells were assigned to the subtype or state corresponding to the highest module score. Overall concordance between ClustoCell clusters and assigned cell subtypes/states was computed (Eq. 14).

### Identification of malignant and non-malignant cells

ClustoCell was applied to 12 scRNA-seq datasets spanning 10 cancer types, including Acute Erythroid Leukaemia (AEL), Acute Lymphoblastic Leukaemia (ALL), Colorectal Cancer (CRC), Head and Neck Squamous Cell Carcinoma (HNSCC), Liver Hepatocellular Carcinoma (LIHC), Pancreatic ductal adenocarcinoma (PDAC), Renal cell carcinoma (RCC), Small Cell Lung Cancer (SCLC), Melanoma, and Glioblastoma.

For each dataset, ClustoCell was run on the full dataset. For datasets containing both malignant and non-malignant epithelial cells, we first used ClustoCell to identify the cluster corresponding to the entire epithelial population. ClustoCell was then applied again within this epithelial cluster to distinguish malignant from normal epithelial cells. This two-step strategy was used because ClustoCell initially groups cells based on overall transcriptomic similarity: normal and malignant epithelial cells share more similar gene expression profiles and markers with each other than with other cell types, such as immune cells, and therefore tend to cluster together before being separated into malignant and normal subpopulations. The resolution parameter was adjusted in each dataset to obtain two major clusters, enabling direct comparison between malignant and non-malignant compartments. All other parameters were kept at their default settings unless stated otherwise. Lastly, the cluster–annotation concordance was quantified (Eq. 14).

The discrepancies between ClustoCell labels and OMA of SCLC were examined in two alternative ways. First, up- and down-regulated genes between concordant malignant and non-malignant cell populations (ground truth labels), identified using Seurat’s *FindMarkers* function, were used to score all cells. Scoring was done using the Singscore rank-based single-sample scoring method^72^ as implemented in the singscore R package v1.28.1. Second, top SCLC-relevant hallmark gene sets were obtained from the MSigDB database using the msigdbr R package (v25.1.1), and discordant and concordant cell populations were scored using the Singscore method.

To further compare ClustoCell with copy-number-based malignancy detection methods, we analysed the prostate cancer scRNA-seq dataset GSE176031 containing biopsy and radical prostatectomy tumour samples together with matched normal tissues. ClustoCell was applied using default parameters. Cell types were assigned based on the CelliVerse Marker DB using *typoClust* (*tissue = “Prostate”*, *condition = “Prostate Cancer”*) function of the celliverse R package. The ERG^+^ malignant cluster identified by ClustoCell was compared with luminal epithelial clusters by examining the expression of established prostate cancer marker genes.

CopyKAT (v1.2.5) was applied to epithelial cell populations comprising the ERG^+^ malignant cluster together with luminal epithelial and additional non-malignant epithelial clusters to provide reference diploid cells, using default parameters unless otherwise stated (*id.type = “S”*, *cell.line = FALSE*, *distance = “euclidean”*). Cells predicted as “aneuploid” were considered malignant.

InferCNA R package (v1.0.0) was applied to count per million (CPM) normalised data from the epithelial cell populations comprising the ERG^+^ malignant cluster together with luminal epithelial and additional non-malignant epithelial clusters. Per-cell CNA signal and CNA correlation were calculated using the package functions *cnaSignal* and *cnaCor*, respectively, and cells with both metrics exceeding their corresponding dataset means were classified as malignant. The proportion of malignant cells identified by each method was then compared across epithelial clusters, with particular focus on the ERG^+^ malignant cluster identified by ClustoCell.

### Analysis of effector–exhaustion programs and clonal expansion in breast cancer

The T/NK cells of the scRNA-seq dataset from Bassez *et al.* (breast cancer) were downloaded from https://lambrechtslab.sites.vib.be/en/single-cell and processed using Seurat v5.3.1. Data were log-normalised. ClustoCell was applied to identify all clusters and subclusters within this dataset.

Expansion status of the cells provided by Bassez *et al.* was binarized at the single-cell level, yielding a dataset-wide expansion prevalence of 0.695. This value was used as the background probability against which subcluster-level expansion was assessed. For each subcluster, the proportion of expanding cells was computed, and subclusters were classified as expansion-enriched if their expansion proportion exceeded the global baseline, thereby accounting for intrinsic class imbalance in the dataset.

For each expansion-enriched subcluster, a 2×2 contingency table was constructed comparing subcluster membership to expansion status, and a two-sided Fisher’s exact test was performed. *P*-values were used as the primary ranking metric, as they provide stable statistical evidence even when subcluster sizes are uneven or contingency-table entries are small. Odds ratios were computed but not used for prioritization due to potential inflation under cell-number imbalance. The four most significantly associated subclusters were selected based on minimum *p*-value and carried forward for patient-level modeling.

Predictive modeling was restricted to pre-treatment biopsies. For each patient, the fractional abundance of selected subclusters and log_10_ T count were provided as features to elastic-net logistic regression (R package glmnet^73^ v4.1, α = 0.5) with 5-fold cross-validation. ROC curves were generated using the R package pROC^74^ v1.19. For comparison, identical modeling was performed using author-defined predictive cell types (CD4_EX, CD4_EX_prolif, CD8_EX, CD8_EX_prolif).

Marker expression was summarised using *Seurat::AverageExpression*, row-scaled, and visualized as heatmaps. For the four expansion-enriched predictive subclusters (C5-Sub2, C5-Sub3, C4-Sub4 and C2-Sub1), state assignments were made by jointly evaluating the original ClustoCell-derived markers (Fig. 6h) and a canonical-marker reference panel covering: inhibitory checkpoints and CD4 exhaustion (*PDCD1, LAG3, CTLA4, TIGIT, TNFRSF18, PRDM1, CXCL13*); cytotoxic effector program (*GZMB, GZMH, PRF1, NKG7, GNLY, CCL4*); class II MHC / interferon licensing (*HLA-DRA, HLA-DRB1, HLA-DRB5*); CD8 stem-like / progenitor exhaustion (*TCF7, LEF1, BCL2*); early TCR-driven activation and survival (*NR4A1, IL2RA, ICOS, TNFRSF4/OX40, BCL2L1, BCL2A1*); TPH-like CD4 exhausted program, including CXCR5^lo^/− and BCL6^lo^/− (*CXCR5*, *BCL6*, *IL21*, *CD40LG*, *ICOS*, *TNFRSF4*, *MAF*, *CCR2*, *CCR5*); terminal versus intermediate/effector-like exhaustion (*TOX*, *ENTPD1*/*CD39*, *CD101*, *CXCR6*, *GZMK*, *CX3CR1*); and CD4 effector-memory lineage anchoring (*CD4, IL7R, ITGB1, ANXA1, KLRB1, CD40LG*). The full expanded panel is shown in Supplementary Figure 22 and informed the cell-state labels used throughout the Results.

Subcluster-level pseudotime trajectories were inferred using Monocle3^75^ and were used exclusively for biological interpretation rather than for predictive modeling. Pseudotime was computed separately within CD8 and CD4 compartments to summarise transcriptional continuity among effector-memory and effector-exhaustion states.

### Differential gene expression and survival analysis of CD4 EM subsets

Differential gene expression analysis was performed within the CD4 effector-memory compartment using Seurat’s *FindMarkers* function, comparing C2-Sub1 and C3-Sub2 subclusters in pre-treatment samples. Cells were subset to CD4 EM annotations, and differential expression was assessed using default Wilcoxon rank-sum testing with Benjamini–Hochberg correction. Genes with adjusted *P* < 0.05 and |log_2_ fold change| > 0.585 were considered significant.

For survival analysis, upregulated and downregulated gene sets were defined from the differential expression results. All significantly upregulated genes meeting the above thresholds were retained, whereas the 20 most strongly downregulated genes were selected based on log2 fold change magnitude. TCGA breast invasive carcinoma (TCGA-BRCA) gene-level normalised RNA-seq expression data (assay “*BRCA_RNASeq2GeneNorm-20160128*”) and matched clinical annotations were obtained using the curatedTCGAData R package (v1.3)^76^ via the MultiAssayExperiment framework. Analysis was restricted to primary tumour samples (TCGA sample code 01) matched to available clinical follow-up; samples with missing survival time, missing vital status, or missing Singscore output were excluded, yielding 936 primary tumours with 119 deaths available for downstream Cox modelling.

Rank-based single-sample scoring was performed using the Singscore method. Gene expression values were first ranked within each sample, and a composite score was calculated by contrasting the average rank of upregulated genes augmented with an additional CD4 effector-memory signature (*CD4*, *CD40LG*, *IL7R*, *ITGB1*, *ANXA1*) to reduce potential confounding by cytotoxic lymphocyte infiltration against the average rank of downregulated genes, yielding a directional measure of the C2-Sub1 transcriptional program.

Overall survival (OS) was constructed from TCGA clinical annotations, with time-to-event calculated as days from initial pathologic diagnosis to death (for uncensored patients) or to last follow-up (for censored patients), and converted to months; observations were administratively censored at 120 months (10 years) to reduce the influence of very long-term follow-up. Cox proportional hazards models were fitted using the survival R package (v3.8; https://CRAN.R-project.org/package=survival). Univariable Cox models were fitted separately for the Singscore-derived C2-Sub1 gene-set score and for each clinical covariate (age at initial pathologic diagnosis and AJCC pathologic tumour stage collapsed into four ordered categories: Stage I, II, III and IV). A multivariable Cox model was then fitted with the gene-set score, age, and pathologic stage entered jointly as predictors, allowing the hazard ratio for the C2-Sub1 gene-set score to be estimated after adjustment for age and stage. Univariable forest plots were generated using the forestmodel R package (v0.6.2; https://CRAN.R-project.org/package=forestmodel). For Kaplan–Meier visualization, samples were dichotomized at the median C2-Sub1 gene-set score and survival curves generated with the survminer R package (v0.5.1; https://github.com/kassambara/survminer), with statistical significance assessed by log-rank testing.

To assess whether the prognostic C2-Sub1 transcriptional program reflected a broader immune-inflamed tumour microenvironment, relative immune cell abundances were estimated for TCGA-BRCA samples using the CIBERSORT LM22 leukocyte signature matrix (https://cibersortx.stanford.edu/)^77^. Spearman rank correlations were calculated between the C2-Sub1 Singscore-derived gene-set score and each inferred immune cell population.

### Analysis of immune circuits predictive of PD-1 response in basal cell carcinoma

scRNA-seq and TCR-seq data from Yost *et al*. were processed using the same pipeline applied to the Bassez cohort. Data were log-normalised. ClustoCell clustering was performed to derive transcriptionally stable subclusters.

Per-patient clonal expansion was defined by comparing pre- and post-treatment clone frequencies using Fisher’s exact test (*P* < 0.05; log₂ fold change > 0.5). Cells belonging to expanded clones were labelled Expanded; patients with at least one expanded clone were classified as Expanders. Expansion labels were binarized at the single-cell level, yielding a dataset-wide expansion prevalence of 0.76, which was used as the background expectation for subcluster-level enrichment.

For each subcluster, expansion enrichment was assessed by comparing its proportion of Expanded cells to the global baseline, followed by Fisher’s exact testing. Subclusters were ranked by *P*-value, and the three most significantly associated states were selected for patient-level modeling. Predictive models were restricted to pre-treatment biopsies and used elastic-net logistic regression (R package glmnet, α = 0.5; 5-fold cross-validation), with AUROC computed using R package pROC.

Lineage-restricted pseudotime analyses were performed separately for CD8 and CD4 T-cell compartments using the same log-normalised expression matrix underlying ClustoCell subclustering. Trajectories were inferred using graph-based pseudotime ordering, with root states defined a priori based on transcriptional root-node inference and biological plausibility. For the CD8 axis, C5-Sub1 (CD8 cytotoxic effector/effector-memory-like state) was selected as the starting point on the basis of its effector-memory transcriptional profile and position in the UMAP embedding. For the CD4 axis, C2-Sub1 — a CXCL13^+^ IL-21^+^ MAF^+^ CD40LG^+^ TPH-like CD4 state — was designated as the root based on transcriptional root-node inference and biological plausibility. Pseudotime values were projected onto the original UMAP embedding for visualization and interpreted qualitatively to assess transcriptional continuity. Pseudotime orderings are interpreted as transcriptional continuity rather than as evidence of *in vivo* developmental conversion.

For cell-state assignment of the six annotated BCC T-cell subclusters (C3-Sub2, C4-Sub3, C5-Sub1, C7-Sub3, C2-Sub2, C2-Sub1), marker expression was summarised using *Seurat::AverageExpression* on log-normalised counts, and each gene was z-scored across all six subclusters for visualization. The panel comprised 52 canonical genes covering: CD8 cytotoxic/effector program (*CD8A, CD8B, GZMB, GZMK, GNLY, NKG7, SLC7A5, TBX21, IFNG*); MHC class II and activation (*HLA-DRA, HLA-DRB1, HLA-DRB5, NR4A1*); CD8 effector-memory survival and chemotaxis (*BCL2A1, TCF7, CCR2, CCR5*); inhibitory receptors and exhaustion markers (*PDCD1, TIGIT, LAG3, TOX, ENTPD1*); regulatory T cell core with memory coat (*FOXP3, CTLA4, IL2RA, TNFRSF4, TNFRSF18, BATF, CD27, LTB, LEF1, BCL2, BCL2L1, ICOS, MAF, GATA3*); Th17 lineage core (*RORC, RORA, IL17A, IL17F, IL22, IL23R, CCR6, KLRB1, BHLHE40, ID2*); and Th17-like helper/TFH exclusion module (*CXCL13, CXCR6, CD40LG, IL21, CXCR5, BCL6*). Exclusion markers for competing lineages were included to confirm lineage specificity: *FOXP3, GATA3, TBX21*, and *IFNG* for Th17 identity; *CD8A, CD8B, GZMB*, and *NKG7* for Treg identity; *RORC* and *IL17A/F* for CD8 effector identity; and *CXCR5* and *BCL6* for GC-TFH exclusion.

## Data availability

All publicly available marker resources and the HGNC gene nomenclature database used to construct the CelliVerse MarkerDB are listed in Supplementary Data 1. The CelliVerse MarkerDB is distributed as a data object (markerDB) within the celliverse R package, available from GitHub (https://github.com/asalavaty/celliverse) and the Comprehensive R Archive Network (CRAN; https://cran.r-project.org/package=celliverse). In addition, the CelliVerse MarkerDB is deposited on GitHub as both dense and sparse matrix files at https://github.com/asalavaty/CelliVerse-Project. All publicly available bulk and single-cell RNA-seq datasets analyzed in this study are listed in Supplementary Data 4 and are additionally archived on Zenodo (DOI: 10.5281/zenodo.20550512).

## Code availability

ClustoCell is implemented as a function within the celliverse R package. The celliverse package also includes an LLM-powered Agent that provides a natural-language interface for applying ClustoCell to user data, facilitating its use by researchers without programming or bioinformatics expertise. The complete source code, documentation, and tutorials for *celliverse* are available from GitHub (https://github.com/asalavaty/celliverse) and CRAN (https://cran.r-project.org/package=celliverse) under the GPL-3 license. All scripts and workflows used to generate the results presented in this study are available at https://github.com/asalavaty/CelliVerse-Project and enable full reproduction of the reported analyses. A lightweight browser-based demonstration of ClustoCell outputs is also available as a Hugging Face Space (https://huggingface.co/spaces/asalavaty/celliverse).

## Acknowledgements

This work was supported by the Prostate Cancer Foundation Challenge Award (2023CHAL4233). S.S. is supported by a National Health and Medical Research Council (NHMRC) Investigator Grant (APP2010070). R.M. was supported by a Prostate Cancer Foundation Young Investigator Award. This work was also supported by the Peter MacCallum Cancer Centre, The University of Melbourne, Dana-Farber Cancer Institute, and Harvard Medical School. We thank Terence P. Speed for valuable discussions and methodological advice that helped inform the development of the rank-based scoring strategy used in ClustoCell. We also thank Nicole Haynes for her valuable feedback on the manuscript.

## Extended Data Figures

**Extended Data Fig. 1:**
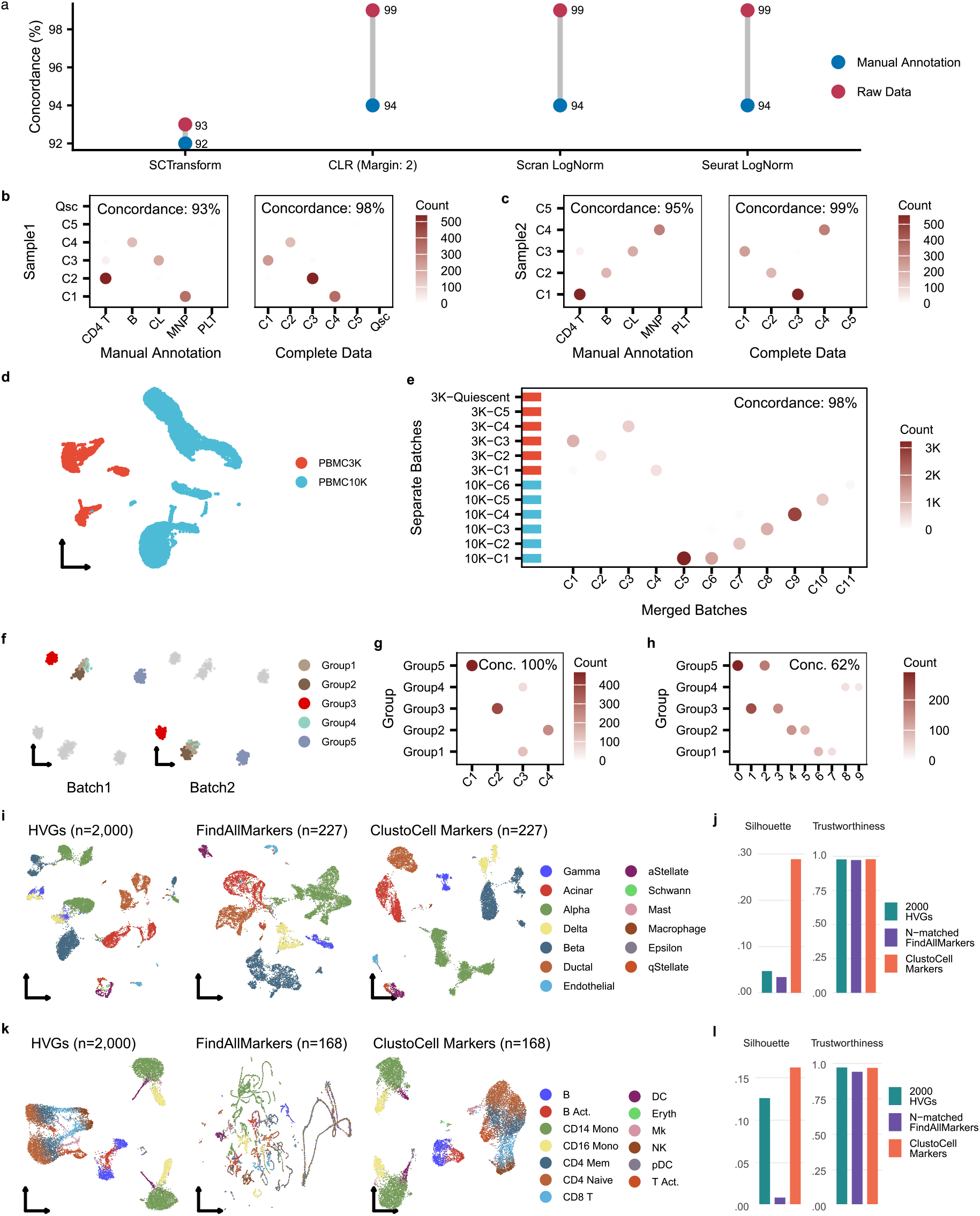
ClustoCell is robust to normalization choices and global batch effects. **a**, Concordance of ClustoCell clusters derived from PBMC3K data normalised using SCTransform, CLR (margin = 2), Scran, and Seurat with those derived from the raw PBMC3K data and with expert manual cell-type annotations. **b**, Random PBMC3K Sample 1: concordance of ClustoCell clusters obtained from the subsampled data with clusters derived from the full dataset, and with manual annotations. **c**, Random PBMC3K Sample 2: same analyses as in panel b. **d**, UMAP of merged PBMC3K and PBMC10K datasets colored by dataset of origin, showing batch-driven separation after standard normalization. **e**, Concordance between ClustoCell clusters obtained from the merged PBMC3K–PBMC10K dataset and clusters obtained from the two datasets analyzed separately. **f**, UMAPs of a Splatter-simulated scRNA-seq dataset colored by ground-truth biological groups across two batches. **g**, Concordance between ClustoCell clusters and ground-truth groups in the simulated dataset. **h**, Concordance between Seurat-derived clusters and ground-truth groups in the simulated dataset. **i,j**, Pancreatic islet cells (Panc8). **i**, UMAP embeddings generated using Seurat’s default dimensionality reduction pipeline with three alternative feature sets: 2,000 highly variable genes (HVGs), ClustoCell-derived top markers, and an N-matched set of Seurat *FindAllMarkers* genes (left to right). **j**, Quantification of cluster separability (mean silhouette width) and embedding trustworthiness for each feature set. **k,l**, IFN-β–stimulated PBMC dataset. **k**, UMAP embeddings generated using HVGs, ClustoCell-derived markers, or Seurat-derived markers. **l**, Silhouette and trustworthiness metrics across methods. Across both datasets, ClustoCell-derived markers yield higher cluster separability while preserving embedding trustworthiness and reducing batch-driven structure.

**Extended Data Fig. 2:**
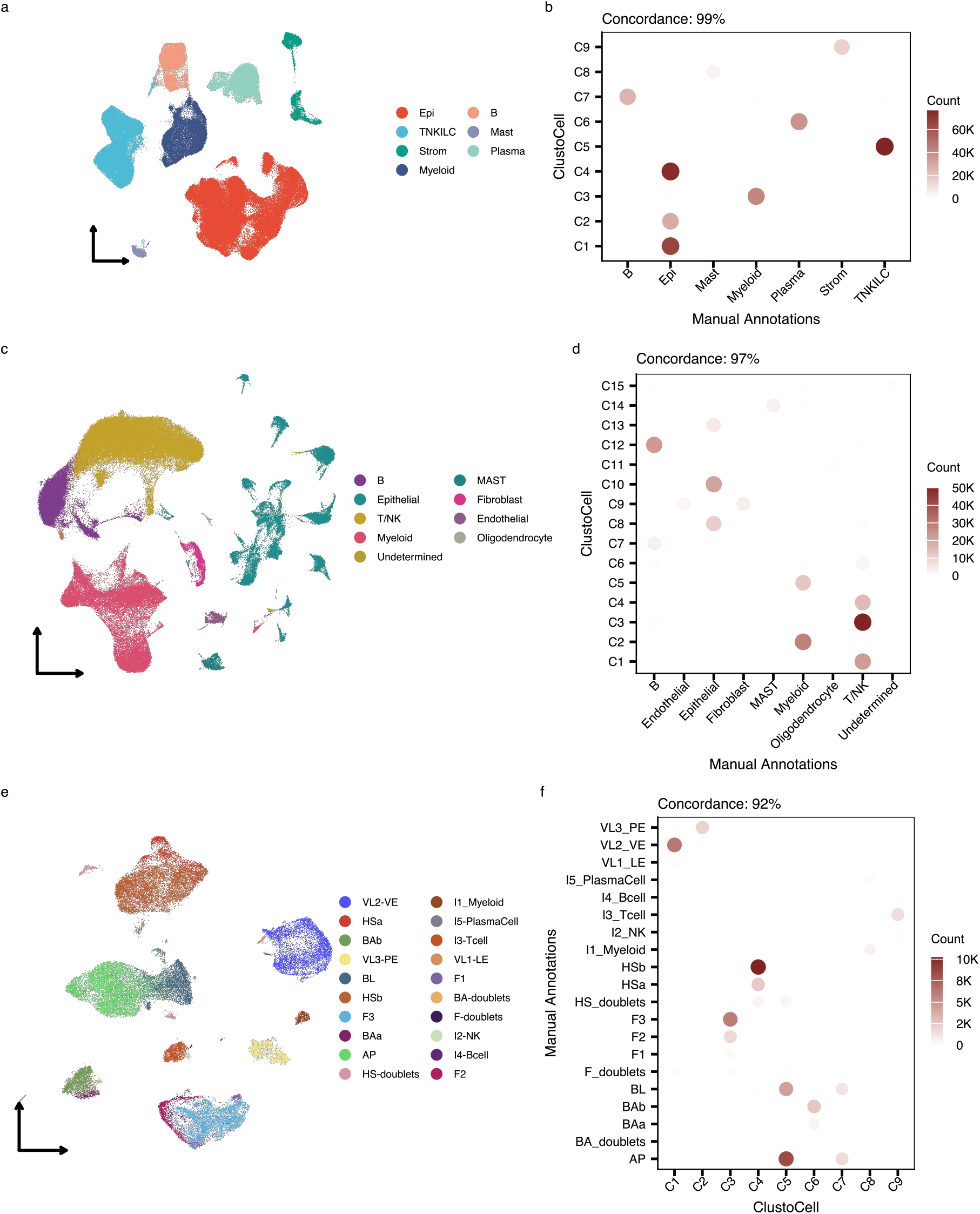
Scalable application of ClustoCell to large single-cell atlases. **a,b**, Human Colon Cancer Atlas. **a**, UMAP visualization of the full dataset colored by expert manual cell type annotations. **b**, Concordance between ClustoCell-derived clusters—obtained by clustering a 20,000-cell sketch followed by label transfer—and expert annotations. **c,d**, LUAD Cell Atlas. **c**, UMAP visualization of the dataset colored by expert annotations. **d**, Concordance between ClustoCell clusters and expert cell type labels following sketch-based clustering and label transfer. **e,f**, Human Breast Cell Atlas. **e**, UMAP visualization colored by expert annotations. **f**, Concordance between ClustoCell-derived clusters and expert annotations.

**Extended Data Fig. 3:**
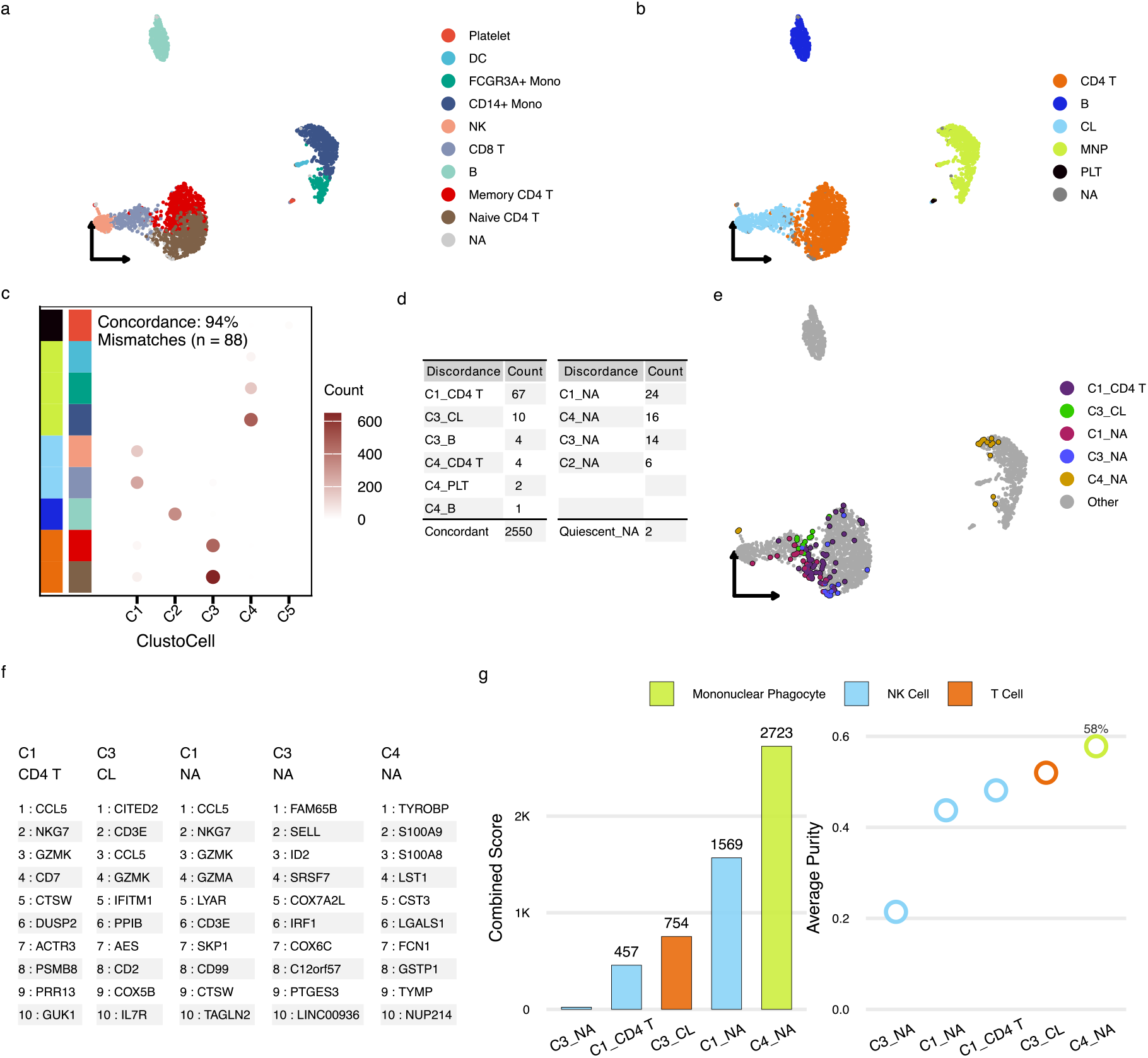
ClustoCell surpasses expert manual annotation of PBMC cell types. **a**, UMAP visualization of the PBMC3K dataset colored by original manual annotations (OMA). **b**, UMAP colored by major lineage-level groupings reflecting shared developmental origins. **c**, Concordance dot plot comparing ClustoCell major clusters with OMA at both fine-grained and major cell type levels. **d**, Summary table of discordant cell subsets between ClustoCell clusters and OMA, including manually annotated NA subsets, with corresponding cell counts. **e**, UMAP highlighting discordant and NA subsets containing at least ten cells. **f**, Top ten ClustoCell-derived markers for each discordant subset. **g**, Marker-based cell type reassignment of discordant and NA subsets, showing assigned cell types, combined scores, and average marker purity within each subset.

**Extended Data Fig. 4:**
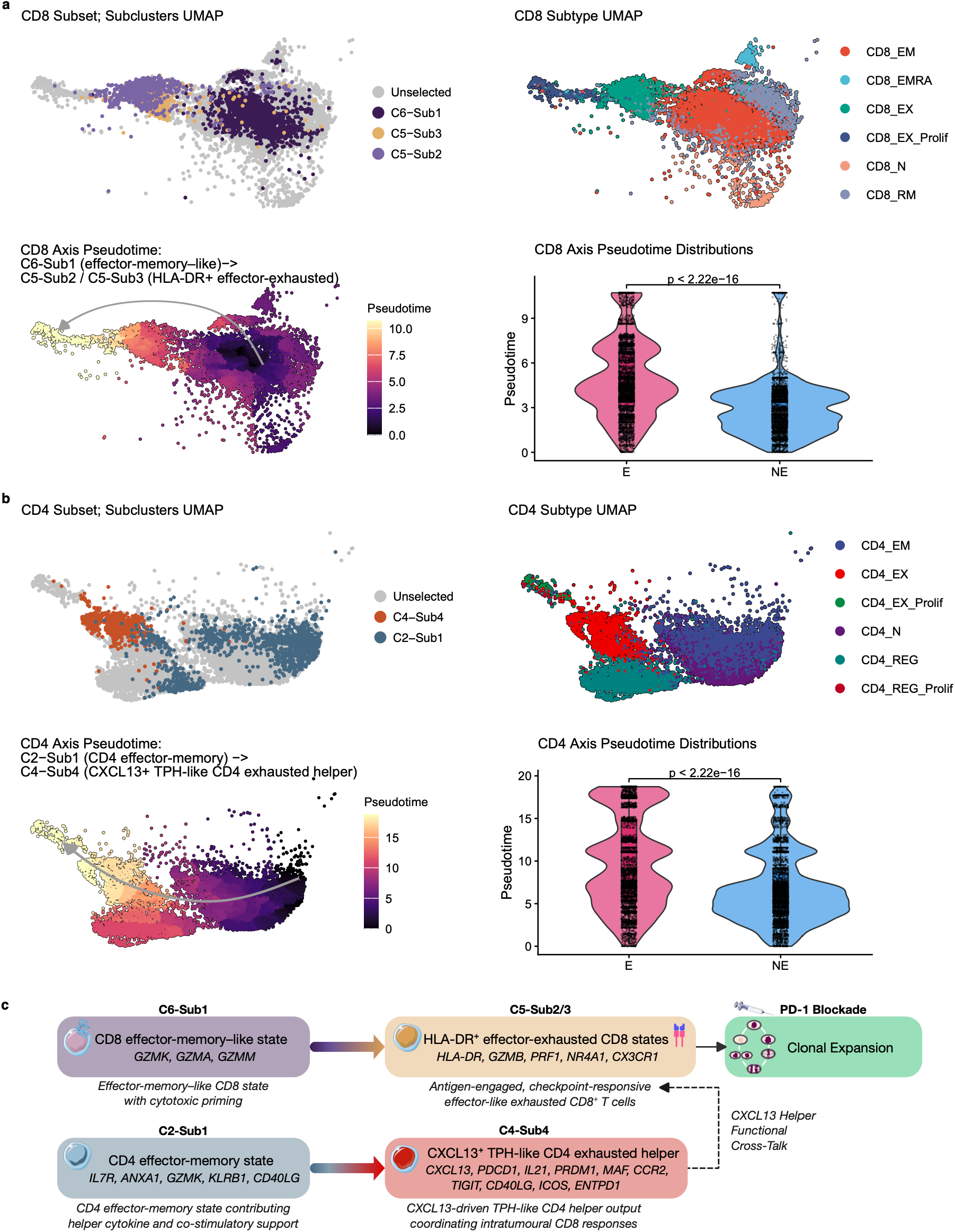
Subcluster-resolved transcriptional continua summarising a coordinated pre-treatment immune circuit in breast cancer. **a,** CD8-compartment pseudotime ordering. Top-left, UMAP coloured by ClustoCell subclusters; top-right, the same UMAP coloured by author-defined CD8 subtypes; bottom-left, the same embedding coloured by inferred pseudotime, illustrating a continuous transcriptional gradient from an effector-memory–like CD8 state (C6-Sub1) toward HLA-DR⁺ effector-exhausted CD8 states (C5-Sub2 and C5-Sub3); bottom-right, CD8-compartment pseudotime distributions in expander (E) versus non-expander (NE) cells, P from two-sided Wilcoxon rank-sum test. **b,** CD4-compartment pseudotime ordering. Top-left, UMAP coloured by ClustoCell subclusters; top-right, the same UMAP coloured by author-defined CD4 subtypes; bottom-left, the same embedding coloured by inferred pseudotime, showing a transcriptional gradient from a CD4 effector-memory state (C2-Sub1) toward a CXCL13⁺ T peripheral helper (TPH)-like CD4 exhausted state (C4-Sub4); bottom-right, CD4-compartment pseudotime distributions in expander versus non-expander cells. **c,** Schematic summarising the inferred pre-treatment Coordinated Effector–Exhaustion Circuit in breast cancer, comprising a CD8 effector-memory–like state, HLA-DR⁺ effector-exhausted CD8 states (combining activated / early-effector-like and CX3CR1⁺ intermediate effector-like exhausted populations), a CD4 effector-memory state, and a CXCL13⁺ TPH-like CD4 exhausted state. Grey arrows denote inferred transcriptional continuity or hypothesised functional coordination.

