## Supplementary Information for "ClustoCell reveals cell states and their markers from single-cell transcriptomes"

### Supplementary Notes

#### Supplementary Note 1

##### A distinct but mechanistically convergent immune circuit predicts PD-1–driven clonal expansion in basal cell carcinoma

To assess cross-tumour generalizability, we applied the same unsupervised ClustoCell workflow to the basal cell carcinoma (BCC) dataset from Yost *et al.* (Yost et al., 2019). Subclustering resolved transcriptionally stable naïve/memory, regulatory, helper, and cytotoxic states with strong concordance to the original annotations (Supplementary Fig. 23a,b), confirming framework robustness across tumour types. Unlike breast cancer — where pre-treatment expansion was associated with a CXCL13$^{+}$ CD4 T peripheral helper (TPH)-like exhausted helper state, an HLA-DR$^{+}$ CX3CR1$^{+}$ intermediate effector-like exhausted CD8 state, an HLA-DR$^{+}$ NR4A1$^{+}$ activated/early-effector-like exhausted CD8 state, and an IL7R$^{+}$ GZMK$^{+}$ CD4 effector-memory state (Fig. 6h; Supplementary Fig. 22) — the BCC dataset emphasized pre-treatment states associated with clonal replacement rather than in-situ reinvigoration of resident exhausted cells (Yost et al., 2019).

Applying the same expansion-aware screening strategy as in Fig. 6, we integrated subclusters with TCR-derived expansion labels and classified expansion-enriched states relative to the global expansion prevalence (0.76). Three subclusters emerged as the most significantly associated with expansion: a CD8 cytotoxic effector/effector-memory-like state (C5-Sub1), an activated effector regulatory T cell (eTreg) state (C3-Sub2), and a CXCL13$^{+}$ IL-17A/F$^{+}$ RORC$^{+}$ Th17-like tissue-resident CD4 state (C4-Sub3) (Supplementary Fig. 23c,d,g). Compositionally, these BCC predictors were distinct from the CXCL13$^{+}$ TPH-like CD4 exhausted helper and intermediate effector-like exhausted CD8 states enriched in expanders in breast cancer, indicating that the ClustoCell-defined expansion-associated circuit takes tumour-specific compositional forms while retaining a shared architecture of coordinated cytotoxic, helper, and regulatory components.

At the patient level, combined pre-treatment fractions of these three subclusters, together with $\log_{10}$ T yield, enabled perfect discrimination between Expander and Non-expander patients (AUROC = 1.00; Supplementary Fig. 23e,f). This performance likely reflects the limited cohort size and should be interpreted as a discovery-cohort observation rather than a validated predictor. The directionality of model coefficients was internally consistent: elevated pre-treatment abundance of the eTreg (C3-Sub2) and Th17-like (C4-Sub3) states, and lower abundance of the CD8 cytotoxic effector/effector-memory-like state (C5-Sub1), together predicted patient-level expansion, whereas none of the three subclusters was individually sufficient. As with the breast-cancer analysis, we interpret this pattern as compositional rather than mechanistic: our cross-sectional pre-treatment data can identify subcluster combinations associated with subsequent clonal expansion but cannot distinguish which are causal versus correlative.

These BCC predictive states are consistent with the clonal-replacement paradigm described by Yost *et al.*, whereby therapeutic response is driven by recruitment and expansion of novel CD8$^+$ clonotypes following PD-1 blockade (Yost et al., 2019). In this context, we identify a distinct pre-treatment immune composition associated with subsequent expansion, comprising increased activated eTreg and tissue-resident Th17-like populations together with reduced abundance of a CD8 cytotoxic effector/effector-memory-like state. C3-Sub2 exhibits a canonical activated effector Treg (eTreg) signature (*FOXP3*, *CTLA4*, *IL2RA*, *BATF*, *TNFRSF4*/*OX40*, *TNFRSF18*/*GITR*, *ICOS*) together with a memory/central-memory transcriptional coat (*LEF1*, *BCL2*, *BCL2L1*, *CD27*, *LTB*, *MAF*) (Supplementary Fig. 23g; Supplementary Fig. 24). Elevated pre-treatment eTreg abundance in expanders may indicate either a pre-existing suppressive niche whose composition permits productive downstream clonal replacement, or a homeostatic negative-feedback response to ongoing inflammation and immune activity in the same tumours — two non-mutually-exclusive interpretations that our cross-sectional pre-treatment data cannot discriminate (Imianowski et al., 2024; Itahashi et al., 2022; Shan et al., 2023). C4-Sub3 co-expresses the full Th17 lineage core (*RORC*, *RORA*, *IL17A*, *IL17F*, *IL22*, *IL23R*, *CCR6*, *KLRB1*) together with tissue-resident helper markers (*CXCL13*, *CXCR6*, *CD40LG*, *IL21*) and exclusion of Th1, Th2, and Treg programs (Supplementary Fig. 23g; Supplementary Fig. 24), consistent with a Th17-like tissue-resident CD4 state; whether this state supports or antagonizes anti-tumour immunity remains disputed in the literature, and our cross-sectional data cannot adjudicate. Finally, C5-Sub1 exhibits a canonical CD8 cytotoxic effector/effector-memory profile (*CD8A*, *CD8B*, *GZMB*, *GZMK*, *GNLY*, *NKG7*, *IFNG*, *TBX21*) with additional expression of HLA-DR class II molecules, *TCF7*, *CCR2*, and *CCR5* (Supplementary Fig. 23g; Supplementary Fig. 24); its reduced pre-treatment abundance in future expanders is compatible with the clonal-replacement paradigm, consistent with the possibility that successful PD-1 responses rely on recruitment and expansion of newly arriving clonotypes rather than the presence of a large pre-existing differentiated cytotoxic compartment; however, our data do not establish whether these cells directly contribute to the expanded post-treatment repertoire. Together, these observations show that ClustoCell identifies compact expansion-associated immune circuits in both breast cancer and BCC, whose composition differs by tumour type while sharing a common architecture of cytotoxic, helper, and regulatory components.

Pseudotime analysis positioned these predictive states along coherent transcriptional continua rather than as isolated endpoints. Within the CD8 compartment, C5-Sub1 (CD8 cytotoxic effector/effector-memory-like state) was positioned at one end of the CD8 pseudotime axis, with an activated effector CD8 state (C7-Sub3) at an intermediate position and a terminally differentiated exhausted CD8 state (C2-Sub2) at the downstream end (Supplementary Fig. 23h). Within the CD4 compartment, C2-Sub1 — a $CXCL13^{+}$ $IL\text{-}21^{+}$ $MAF^{+}$ $CD40LG^{+}$ TPH-like CD4 helper state selected as the transcriptional root node on the basis of root-node inference and biological plausibility — was positioned upstream of the $CXCL13^{+}$ Th17-like tissue-resident CD4 state (C4-Sub3) along the CD4 pseudotime axis (Supplementary Fig. 23i). As in the breast-cancer analysis, these pseudotime orderings are interpreted as transcriptional continuity rather than proof of *in vivo* developmental conversion, given the cross-sectional pre-treatment design and the absence of longitudinal T-cell-receptor sharing across pseudotime endpoints. Taken together, ClustoCell identifies compact expansion-associated immune circuits in both breast cancer and BCC, whose compositional forms differ by tumour type while sharing a common architecture of cytotoxic, helper, and regulatory components.

### Supplementary Figures

#### Supplementary Figure 1

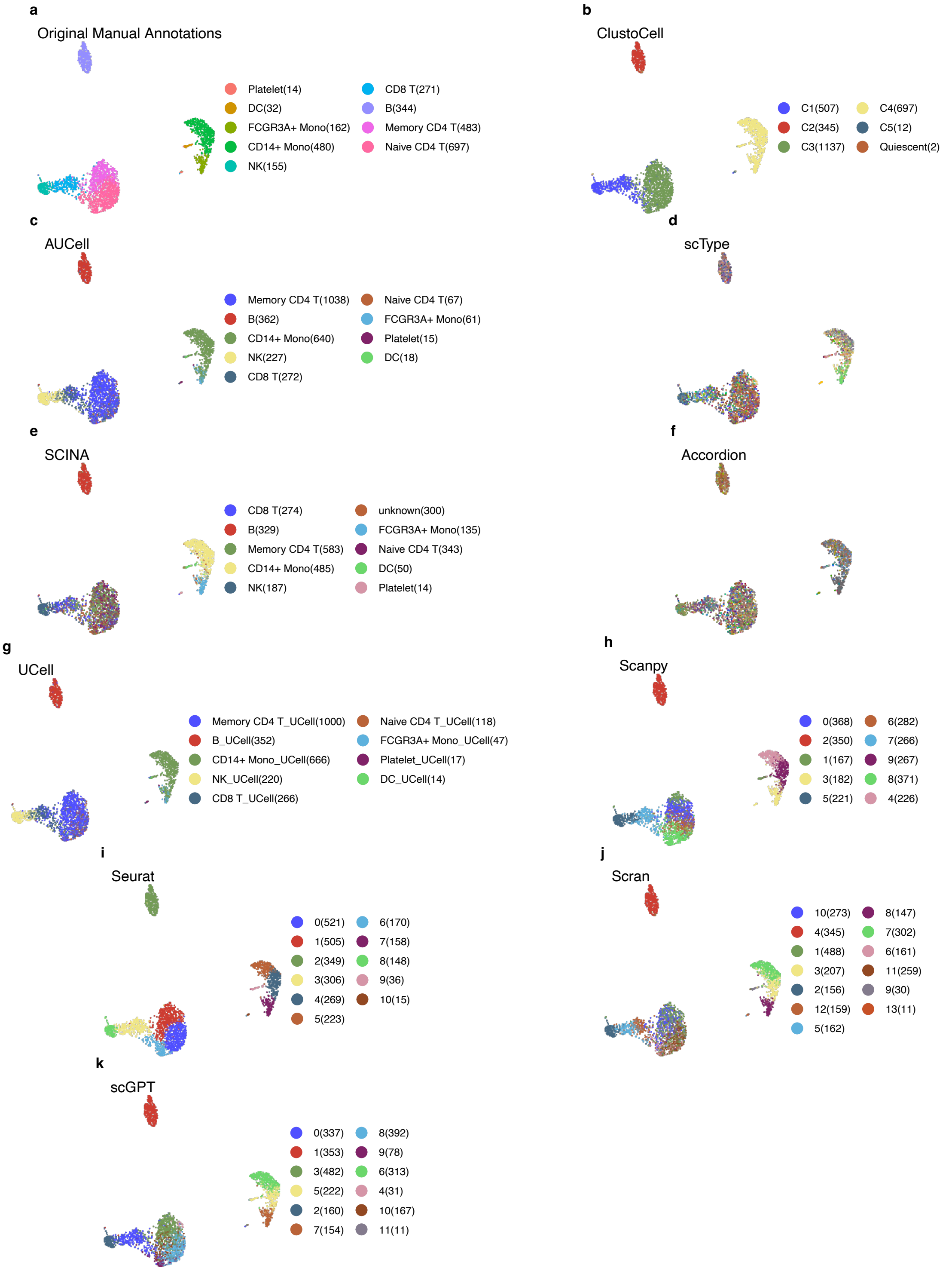

**Supplementary Fig. 1: UMAP Plots of PBMC3K Dataset Colored by Original Manual Annotations and Ten Clustering and Cell Labeling Methods.**

UMAP embeddings of the 10x Genomics PBMC3K dataset, with each panel colour-coding cells by a different annotation/clustering assignment to enable side-by-side comparison of method-specific recovery of canonical PBMC cell types. **a**, original manual annotations. **b**, ClustoCell. **c**, AUCell. **d**, scType. **e**, SCINA. **f**, Accordion. **g**, UCell. **h**, Scanpy. **i**, Seurat. **j**, Scran. **k**, scGPT. Legends are omitted for panels in which a method produced more than 20 clusters.

### Supplementary Figure 2

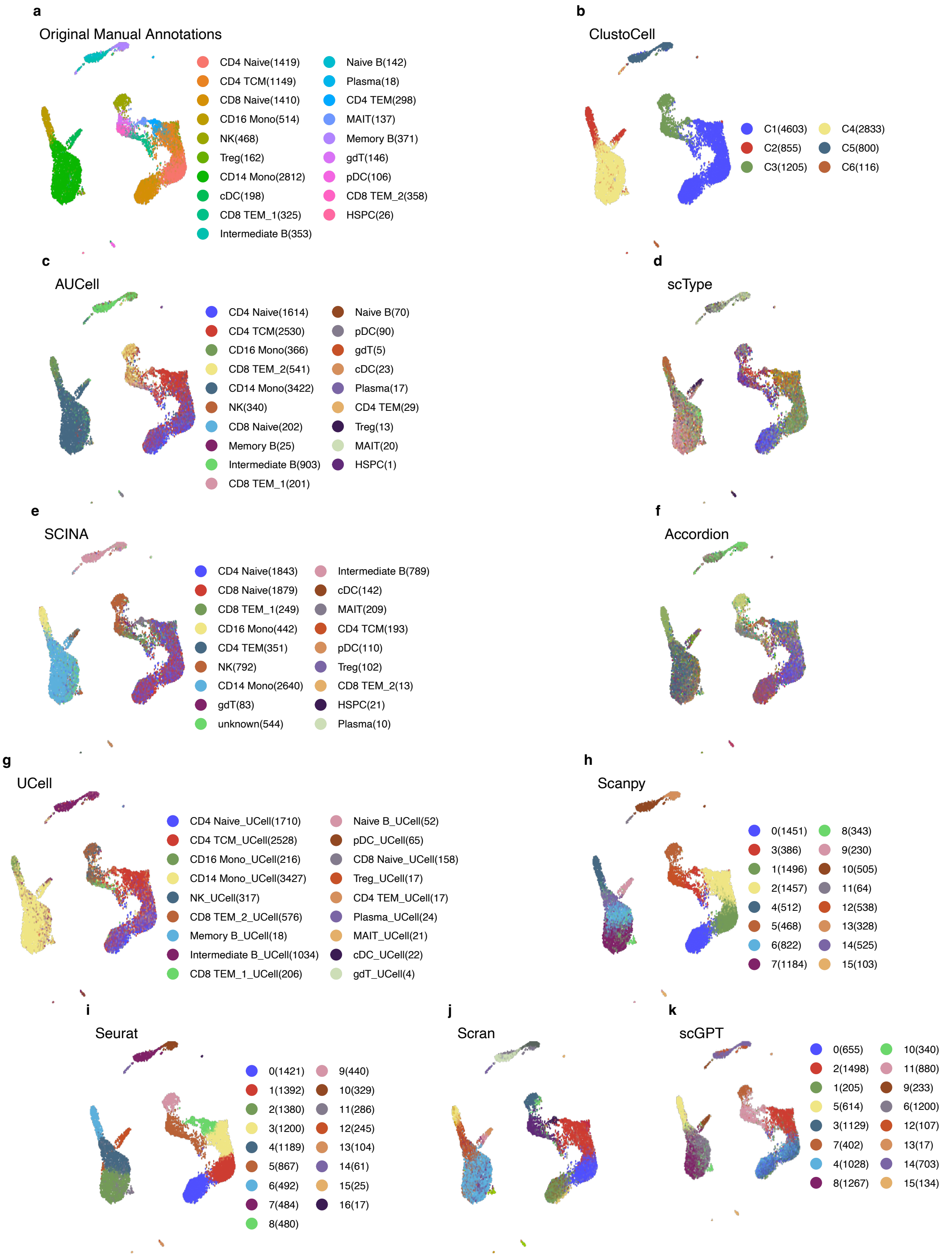

**Supplementary Fig. 2: UMAP Plots of PBMC10K Dataset Colored by Original Manual Annotations and Ten Clustering and Cell Labeling Methods**

UMAP embeddings of the 10x Genomics PBMC10K dataset, with each panel coloured by a different annotation/clustering assignment for side-by-side comparison of method-specific recovery of canonical PBMC cell types. **a**, original manual annotations. **b**, ClustoCell. **c**, AUCell. **d**, scType. **e**, SCINA. **f**, Accordion. **g**, UCell. **h**, Scanpy. **i**, Seurat. **j**, Scran. **k**, scGPT. Legends are omitted for panels in which a method produced more than 20 clusters.

### Supplementary Figure 3

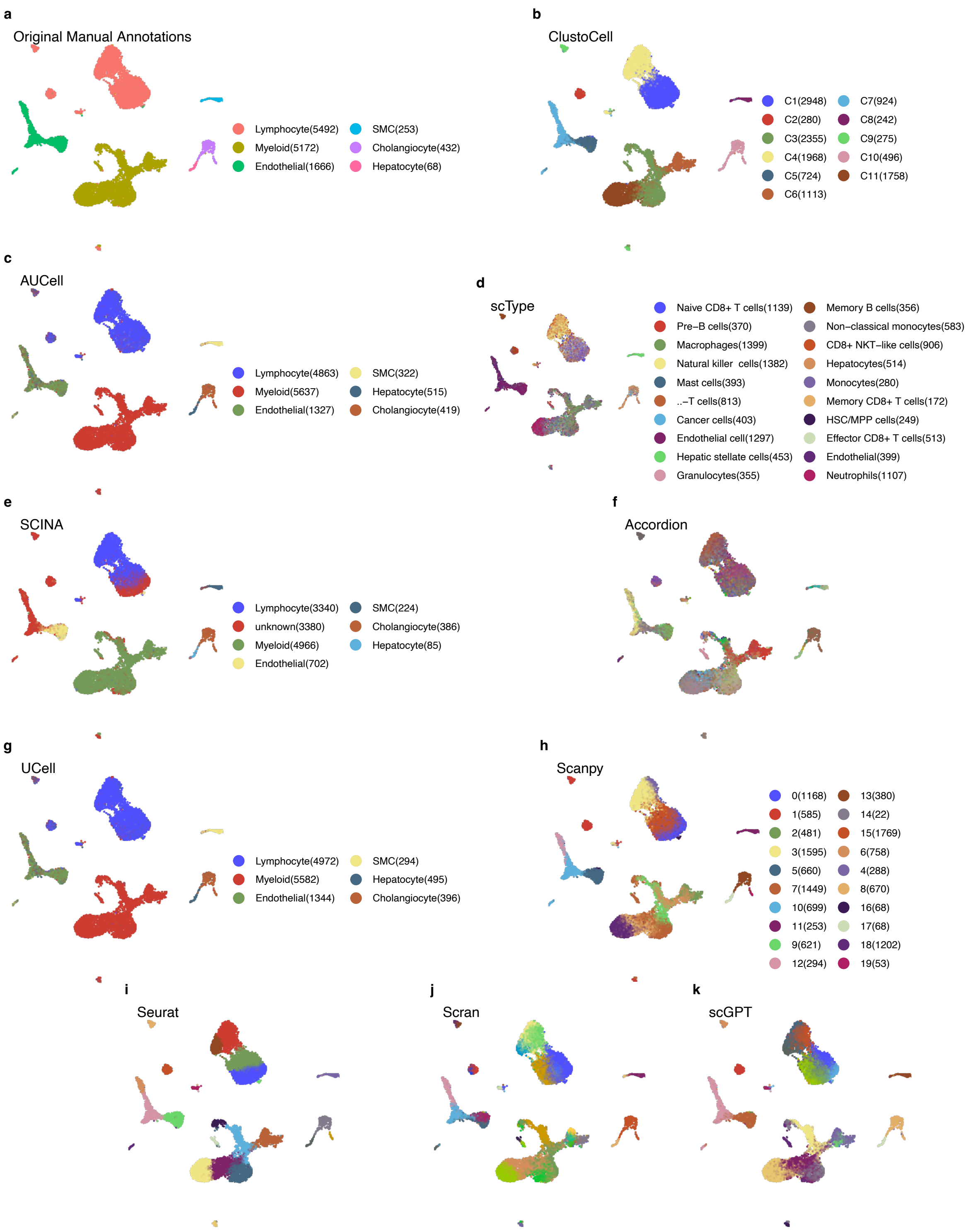

**Supplementary Fig. 3: UMAP Plots of Peritumoral Normal Liver Dataset Colored by Original Manual Annotations and Ten Clustering and Cell Labeling Methods.**
UMAP embeddings of the peritumoral (tumour-adjacent) normal liver scRNA-seq dataset, with each panel coloured by a different annotation/clustering assignment to compare method-specific recovery of hepatic and immune cell states. **a**, original manual annotations. **b**, ClustoCell. **c**, AUCell. **d**, scType. **e**, SCINA. **f**, Accordion. **g**, UCell. **h**, Scanpy. **i**, Seurat. **j**, Scran. **k**, scGPT. Legends are omitted for panels in which a method produced more than 20 clusters.

### Supplementary Figure 4

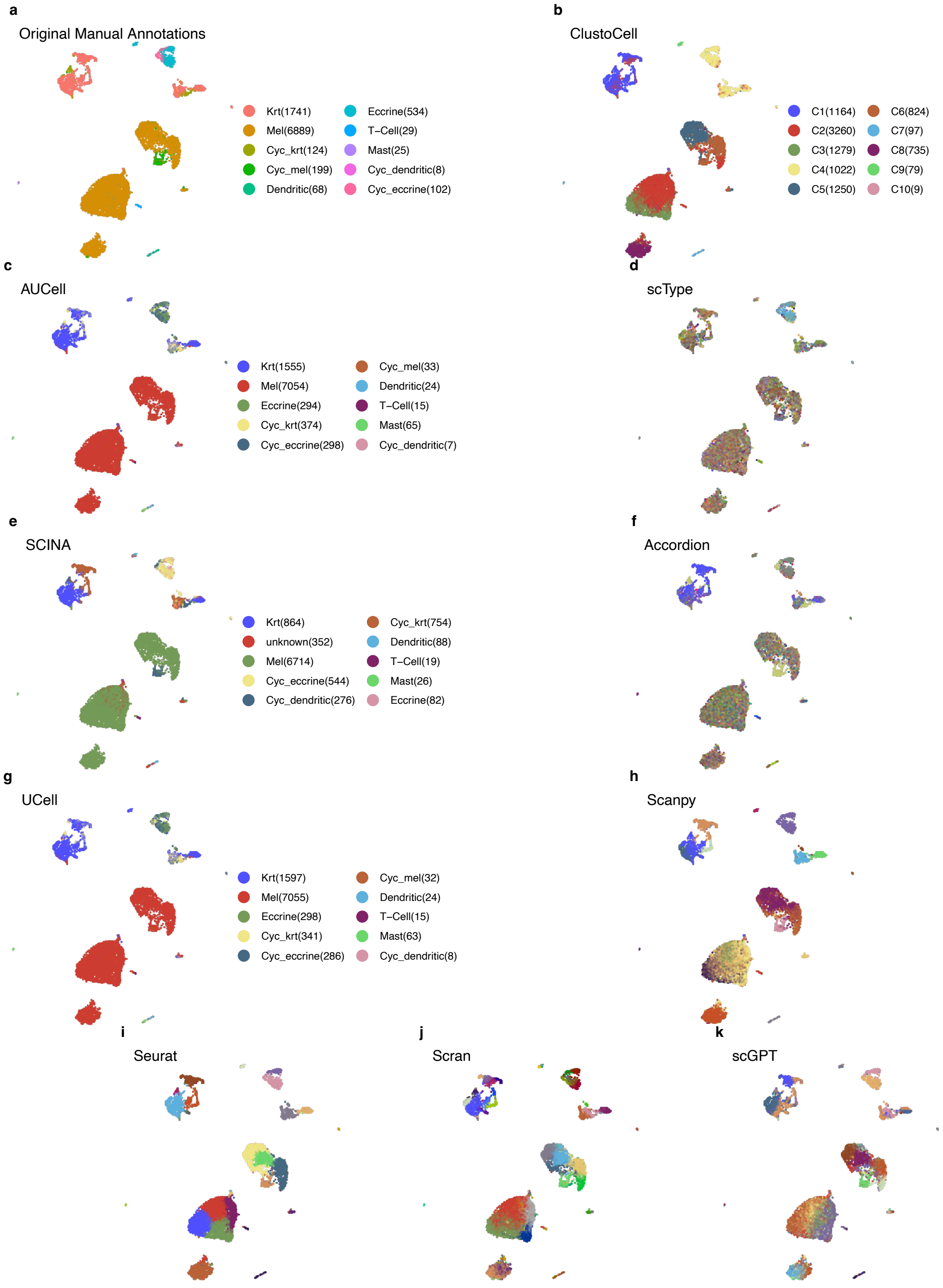

**Supplementary Fig. 4: UMAP Plots of Melanocyte−Melanoma Dataset Colored by Original Manual Annotations and Ten Clustering and Cell Labeling Methods.**
UMAP embeddings of the human melanocyte–melanoma scRNA-seq dataset, with each panel coloured by a different annotation/clustering assignment to compare method-specific recovery of melanocytic and tumour-microenvironment populations. **a**, original manual annotations. **b**, ClustoCell. **c**, AUCell. **d**, scType. **e**, SCINA. **f**, Accordion. **g**, UCell. **h**, Scanpy. **i**, Seurat. **j**, Scran. **k**, scGPT. Legends are omitted for panels in which a method produced more than 20 clusters.

#### Supplementary Figure 5

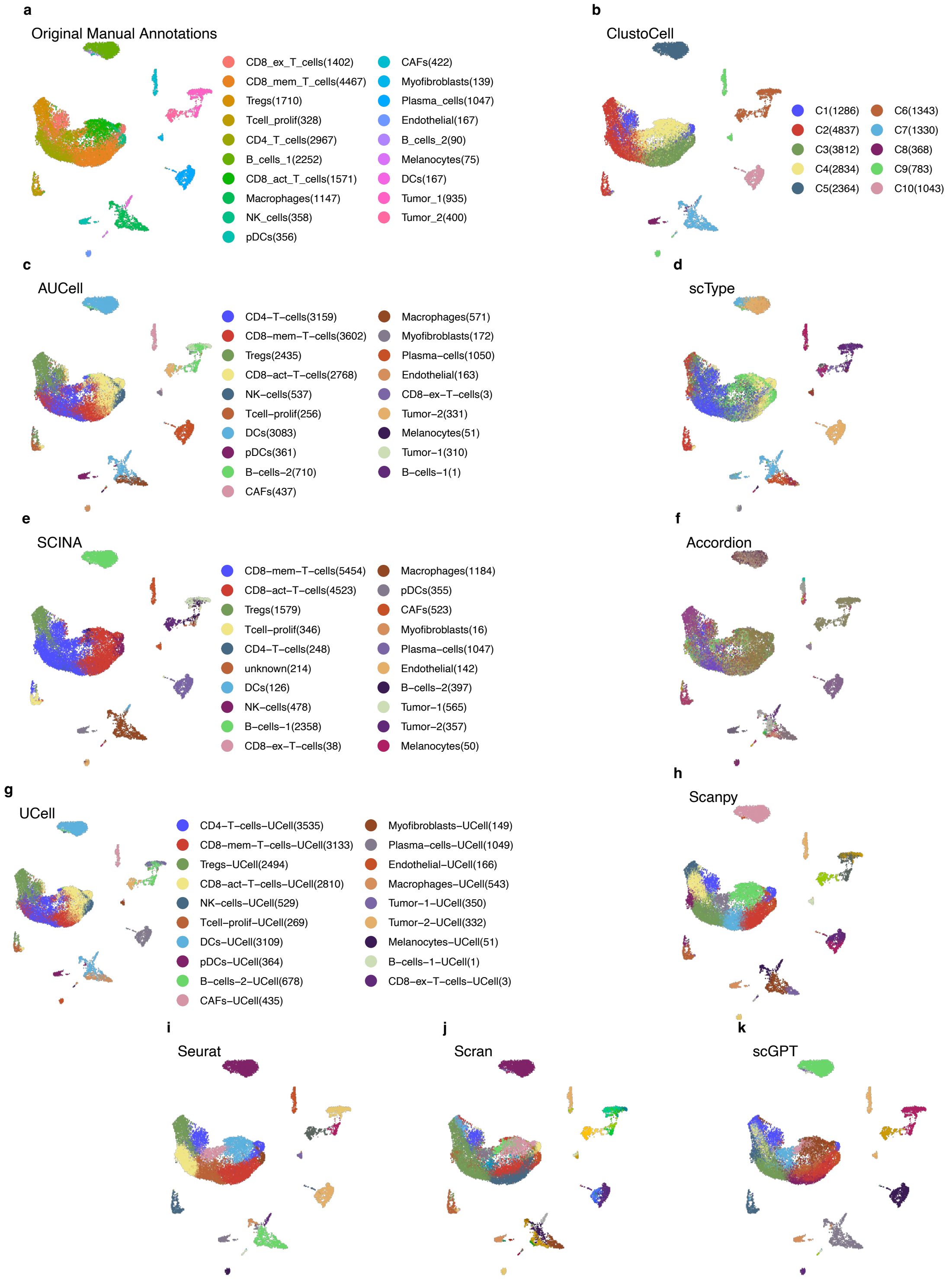

**Supplementary Fig. 5: UMAP Plots of Basal Cell Carcinoma Dataset Colored by Original Manual Annotations and Ten Clustering and Cell Labeling Methods.**
UMAP embeddings of the basal cell carcinoma (BCC) scRNA-seq dataset, with each panel coloured by a different annotation/clustering assignment to compare method-specific recovery of malignant and stromal/immune populations. **a**, original manual annotations. **b**, ClustoCell. **c**, AUCell. **d**, scType. **e**, SCINA. **f**, Accordion. **g**, UCell. **h**, Scanpy. **i**, Seurat. **j**, Scran. **k**, scGPT. Legends are omitted for panels in which a method produced more than 20 clusters.

#### Supplementary Figure 6

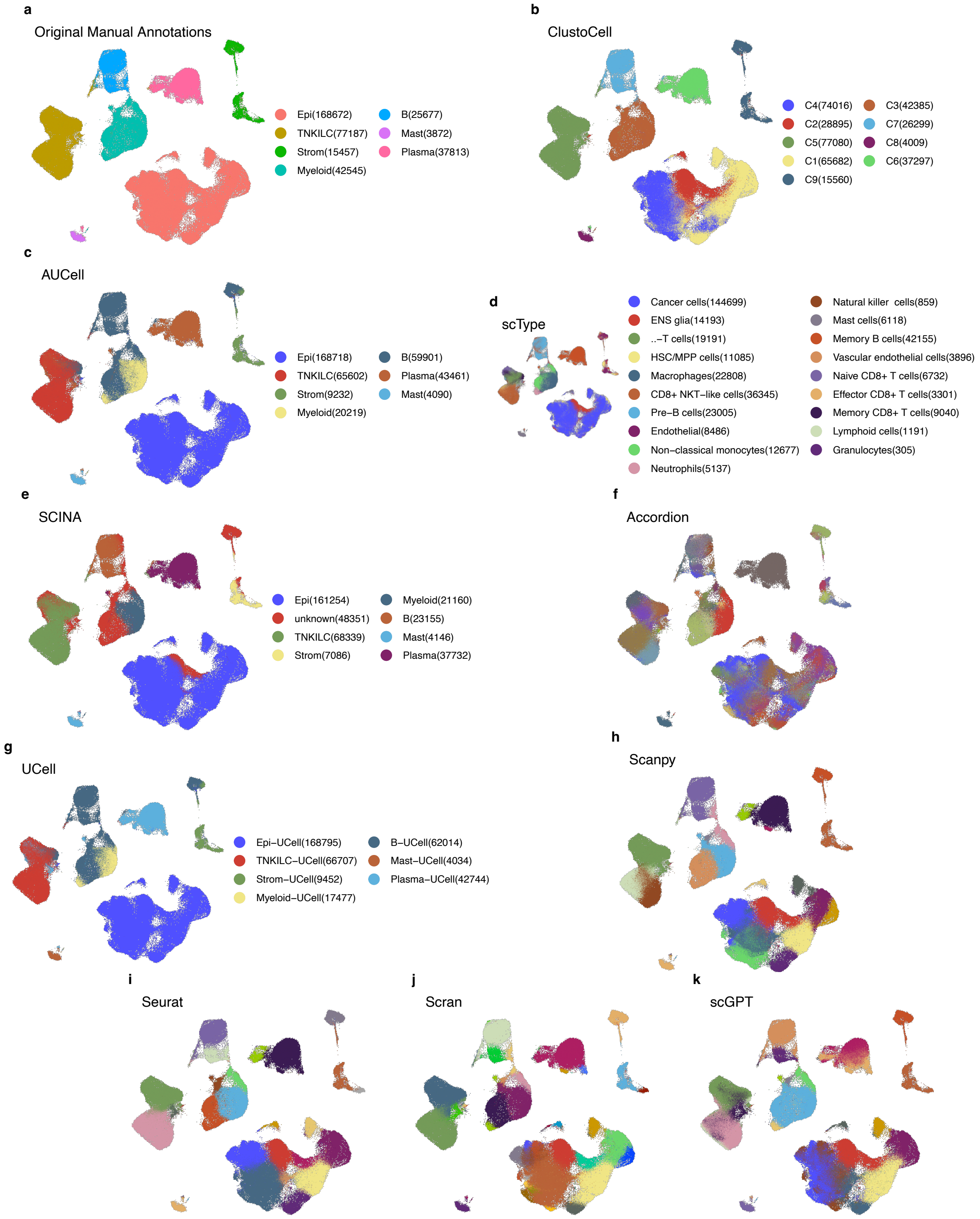

**Supplementary Fig. 6: UMAP Plots of Colon Cancer Atlas Dataset Colored by Original Manual Annotations and Ten Clustering and Cell Labeling Methods.**
UMAP embeddings of the human colon cancer cell atlas scRNA-seq dataset, with each panel coloured by a different annotation/clustering assignment to compare method-specific recovery of epithelial, stromal, and immune populations. **a**, original manual annotations. **b**, ClustoCell. **c**, AUCell. **d**, scType. **e**, SCINA. **f**, Accordion. **g**, UCell. **h**, Scanpy. **i**, Seurat. **j**, Scran. **k**, scGPT. Legends are omitted for panels in which a method produced more than 20 clusters.

### Supplementary Figure 7

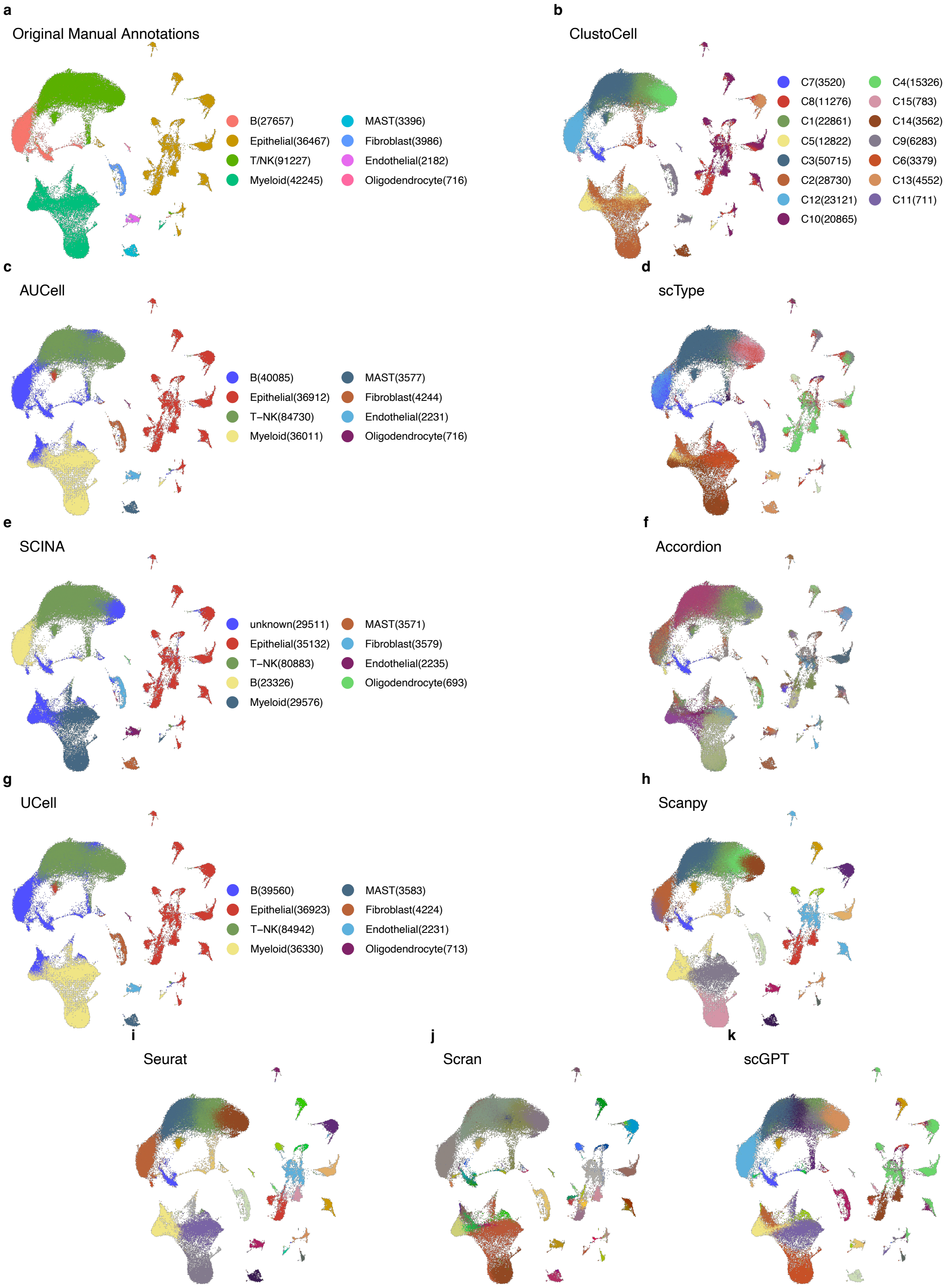

**Supplementary Fig. 7: UMAP Plots of LUAD Cell Atlas Dataset Colored by Original Manual Annotations and Ten Clustering and Cell Labeling Methods.**
UMAP embeddings of the lung adenocarcinoma (LUAD) cell atlas scRNA-seq dataset (GSE131907), with each panel coloured by a different annotation/clustering assignment to compare method-specific recovery of malignant, stromal, and immune populations. **a**, original manual annotations. **b**, ClustoCell. **c**, AUCell. **d**, scType. **e**, SCINA. **f**, Accordion. **g**, UCell. **h**, Scanpy. **i**, Seurat. **j**, Scran. **k**, scGPT. Legends are omitted for panels in which a method produced more than 20 clusters.

### Supplementary Figure 8

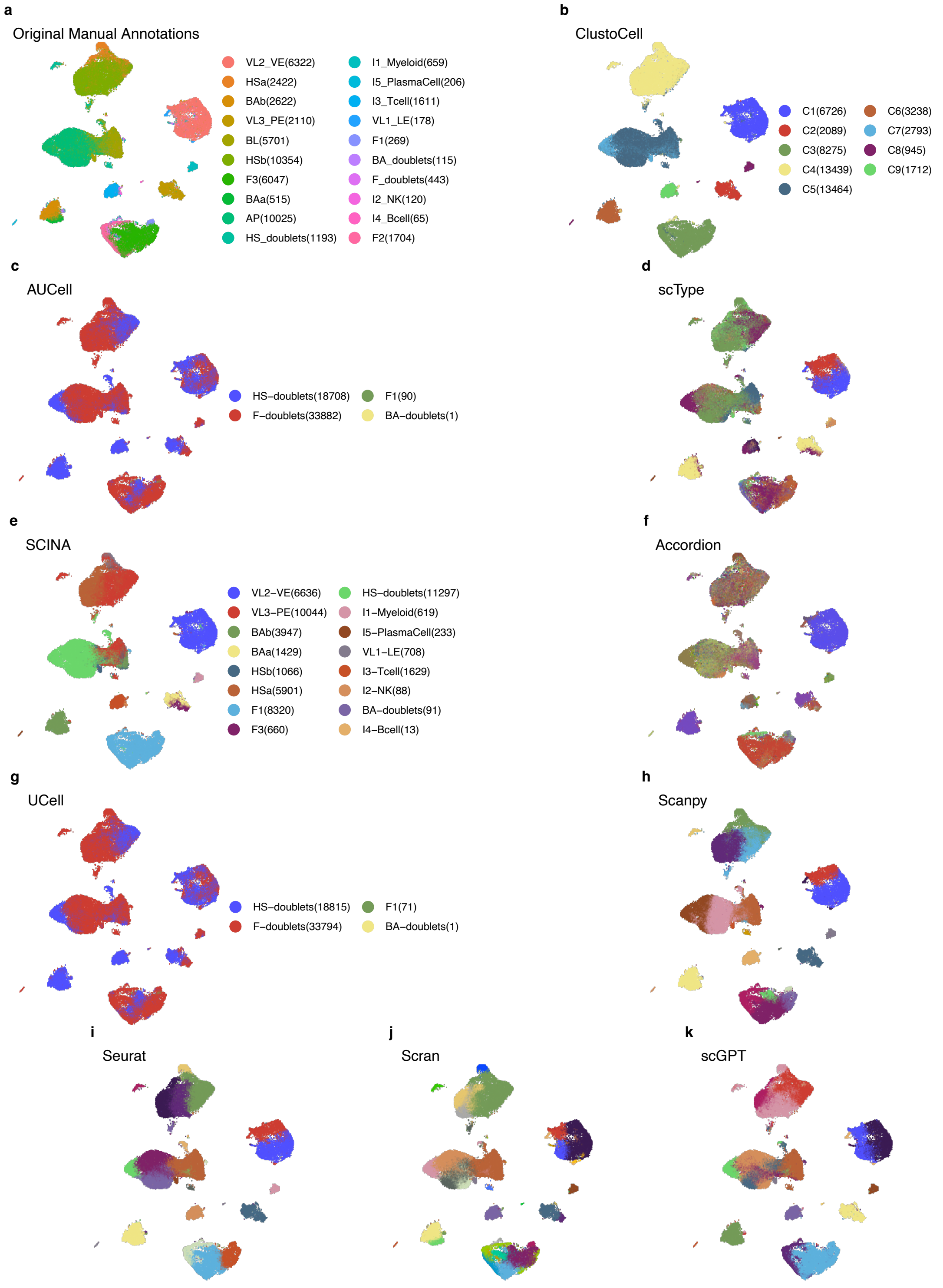

**Supplementary Fig. 8: UMAP Plots of Human Breast Cell Atlas Dataset Colored by Original Manual Annotations and Ten Clustering and Cell Labeling Methods.**
UMAP embeddings of the human breast cell atlas scRNA-seq dataset, with each panel coloured by a different annotation/clustering assignment to compare method-specific recovery of epithelial, stromal, and immune populations of the breast. **a**, original manual annotations. **b**, ClustoCell. **c**, AUCell. **d**, scType. **e**, SCINA. **f**, Accordion. **g**, UCell. **h**, Scanpy. **i**, Seurat. **j**, Scran. **k**, scGPT. Legends are omitted for panels in which a method produced more than 20 clusters.

### Supplementary Figure 9

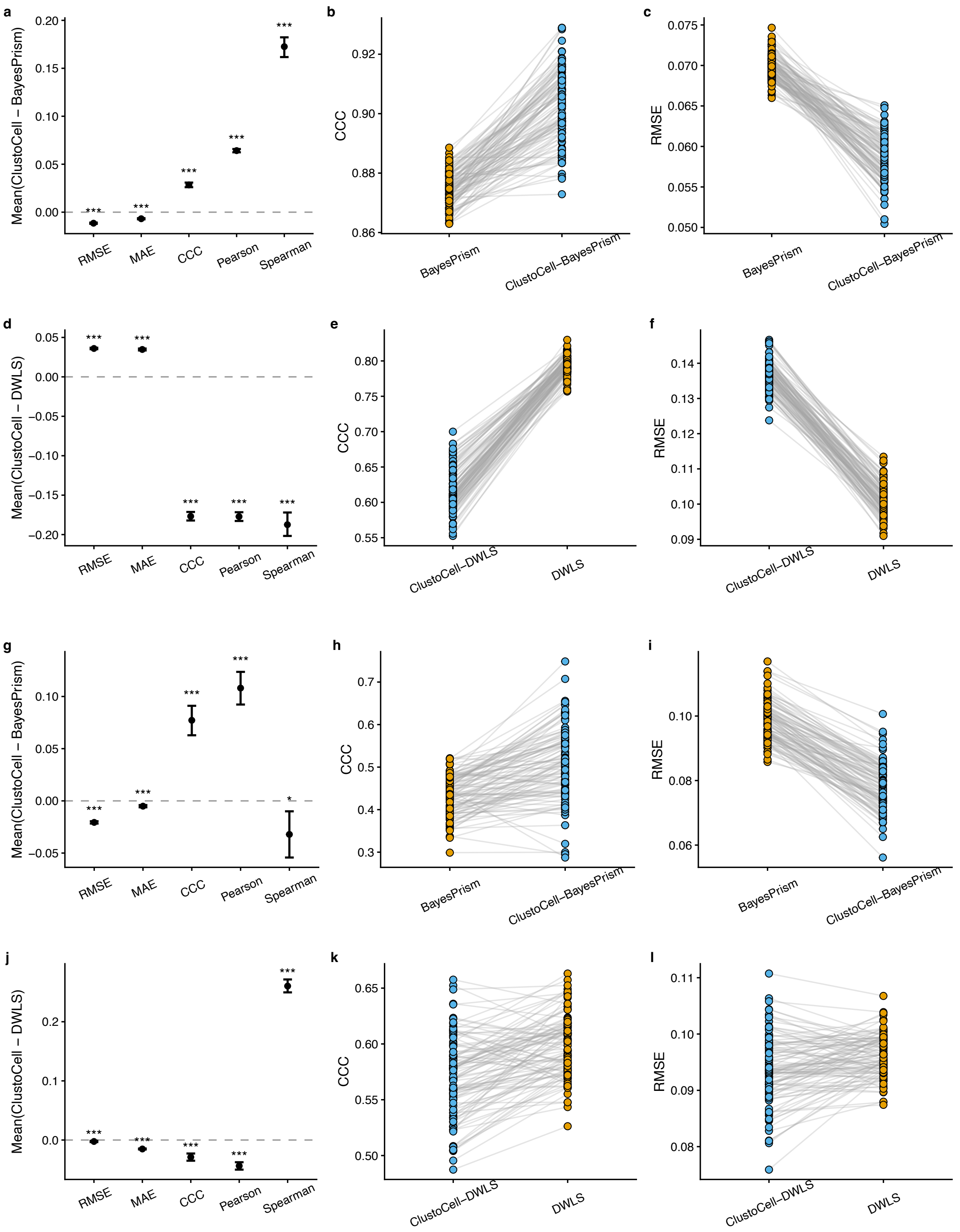

**Supplementary Fig. 9: Benchmarking of cell type deconvolution using ClustoCell-derived markers across liver and prostate datasets.**

**a–f**, Liver benchmark. Benchmarking of cell type deconvolution using published default marker selection strategies and ClustoCell-derived markers with BayesPrism (**a–c**) and DWLS (**d–f**). **a**,**d**, Forest plots summarizing the mean paired differences (ClustoCell − default) across five deconvolution performance metrics, including Pearson correlation, Spearman correlation, concordance correlation coefficient (CCC), root mean squared error (RMSE), and mean absolute error (MAE), with bootstrap 95% confidence intervals. **b**,**e**, Paired comparisons of concordance correlation coefficient (CCC). **c**,**f**, Paired comparisons of root mean squared error (RMSE). Each pair of connected points represents deconvolution of the same pseudobulk sample using the published default marker selection strategy or ClustoCell-derived markers while keeping the underlying deconvolution algorithm unchanged. **g–l**, Prostate benchmark. Benchmarking of cell type deconvolution using published default marker selection strategies and ClustoCell-derived markers with BayesPrism (**g–i**) and DWLS (**j–l**). g,j, Forest plots summarizing the mean paired differences (ClustoCell − default) across five deconvolution performance metrics with bootstrap 95% confidence intervals. **h**,**k**, Paired comparisons of concordance correlation coefficient (CCC). **i**,**l**, Paired comparisons of root mean squared error (RMSE). Each pair of connected points represents deconvolution of the same pseudobulk sample using the published default marker selection strategy or ClustoCell-derived markers while keeping the underlying deconvolution algorithm unchanged.

#### Supplementary Figure 10

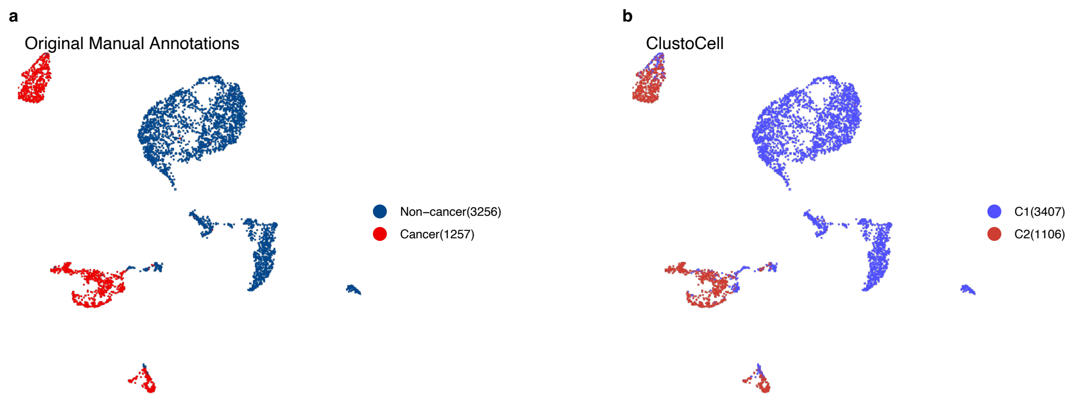

**Supplementary Fig. 10: UMAP Plots of GSE72056 Dataset Colored by Original Manual Annotations and ClustoCell.**
Side-by-side UMAP embeddings of the GSE72056 melanoma scRNA-seq dataset showing concordance between expert manual annotations and ClustoCell-derived clusters. **a**, original manual annotations. **b**, ClustoCell clusters.

#### Supplementary Figure 11

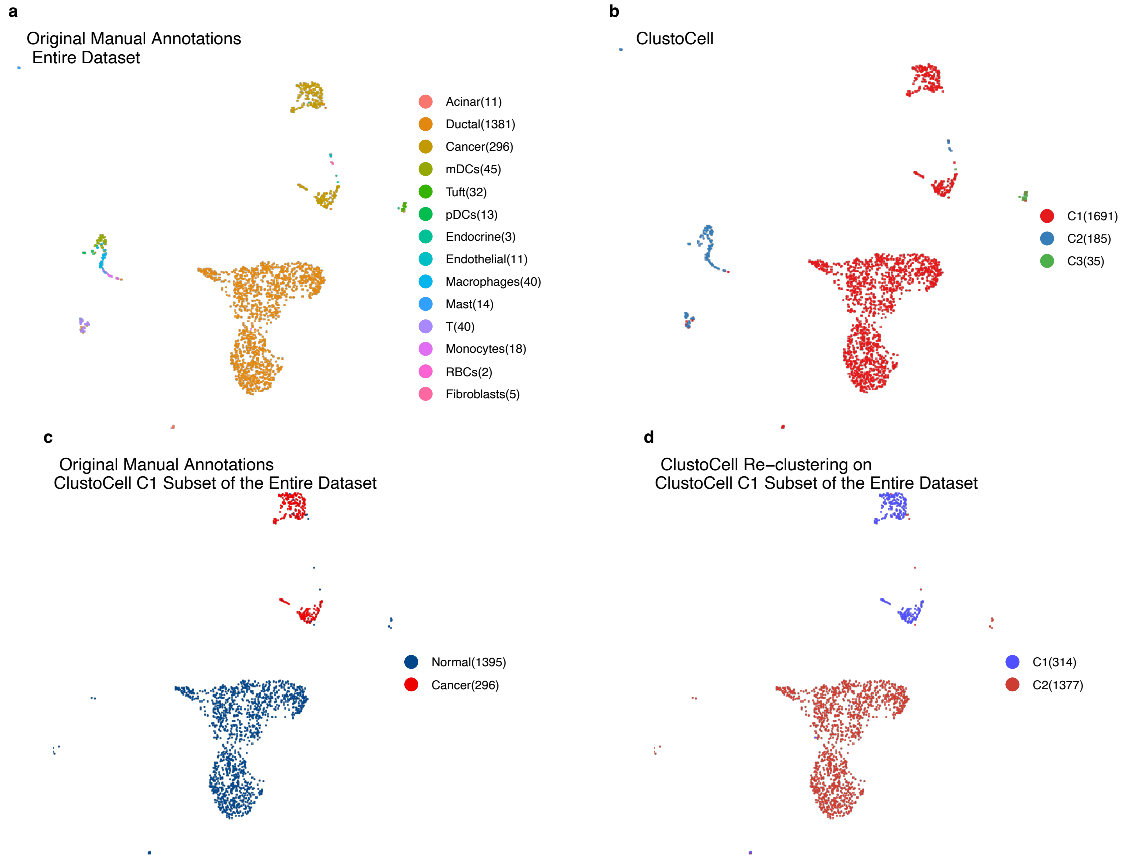

**Supplementary Fig. 11: UMAP Plots of GSE111672 (PDAC−A) Dataset Colored by Original Manual Annotations and ClustoCell.**
UMAP embeddings of the GSE111672 PDAC-A pancreatic ductal adenocarcinoma scRNA-seq dataset, comparing expert manual annotations with ClustoCell clustering at the full-dataset level and after ClustoCell-driven re-clustering of a selected first-level subset (cluster C1). **a**, original manual annotations of the entire dataset. **b**, ClustoCell clusters of the entire dataset. **c**, original manual annotations restricted to the ClustoCell C1 subset. **d**, ClustoCell re-clustering of the same C1 subset, illustrating the within-cluster substructure recovered at higher resolution.

#### Supplementary Figure 12

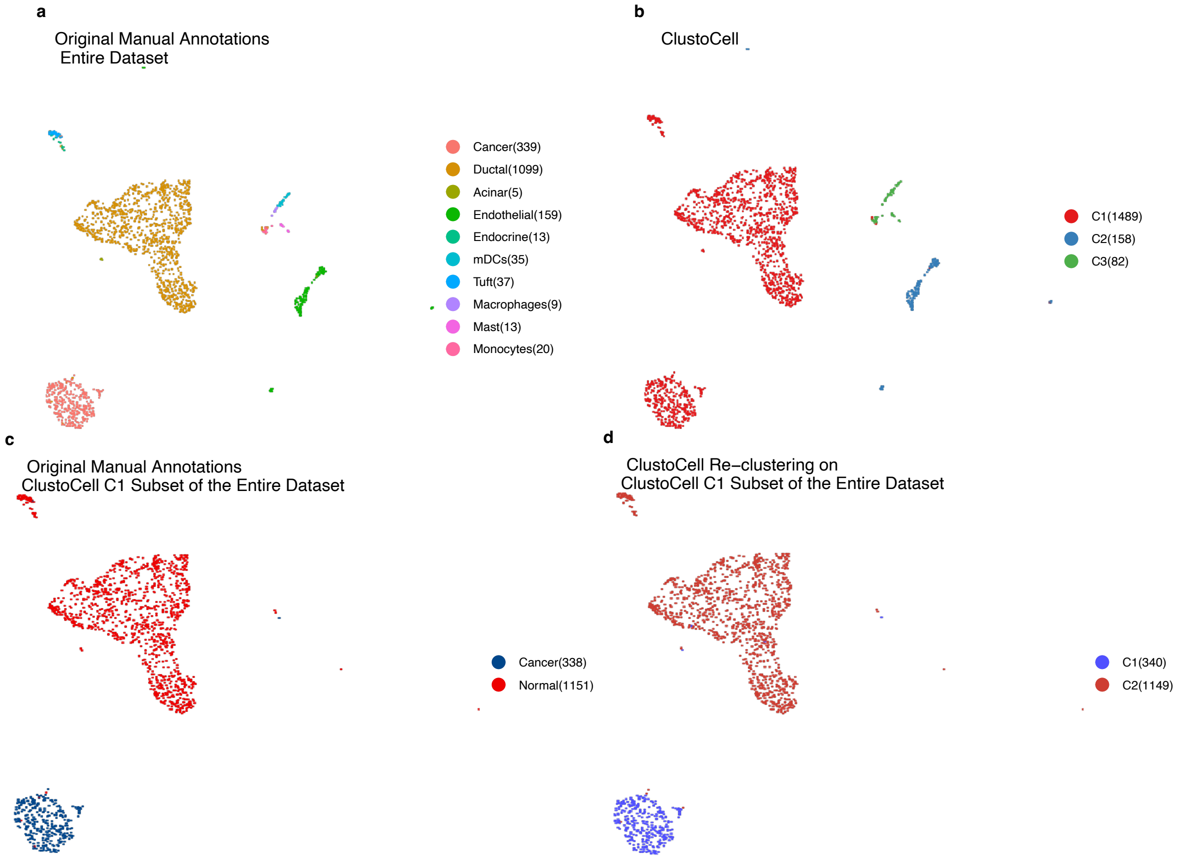

**Supplementary Fig. 12: UMAP Plots of GSE111672 (PDAC−B) Dataset Colored by Original Manual Annotations and ClustoCell.**
UMAP embeddings of the GSE111672 PDAC-B pancreatic ductal adenocarcinoma scRNA-seq dataset, comparing expert manual annotations with ClustoCell clustering at the full-dataset level and after ClustoCell-driven re-clustering of a selected first-level subset (cluster C1). **a**, original manual annotations of the entire dataset. **b**, ClustoCell clusters of the entire dataset. **c**, original manual annotations restricted to the ClustoCell C1 subset. **d**, ClustoCell re-clustering of the same C1 subset, illustrating the within-cluster substructure recovered at higher resolution.

#### Supplementary Figure 13

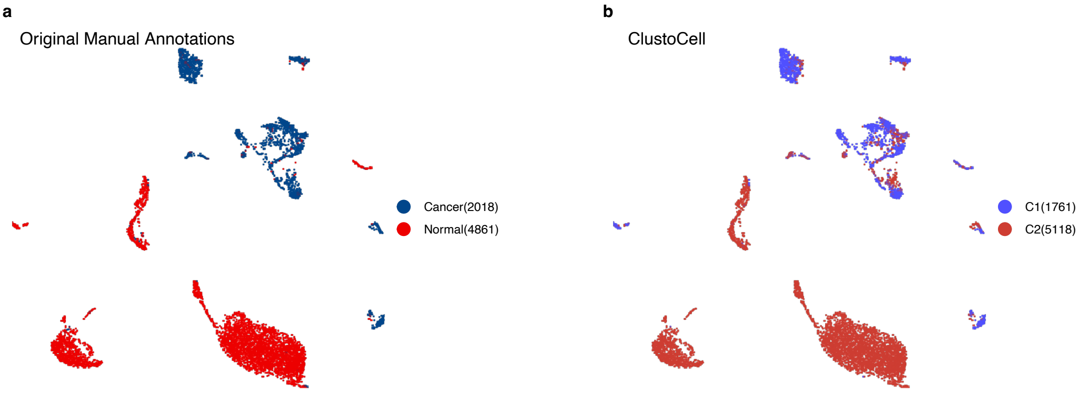

**Supplementary Fig. 13: UMAP Plots of GSE115978 Dataset Colored by Original Manual Annotations and ClustoCell.**
Side-by-side UMAP embeddings of the GSE115978 melanoma scRNA-seq dataset showing concordance between expert manual annotations and ClustoCell-derived clusters. **a**, original manual annotations. **b**, ClustoCell clusters.

#### Supplementary Figure 14

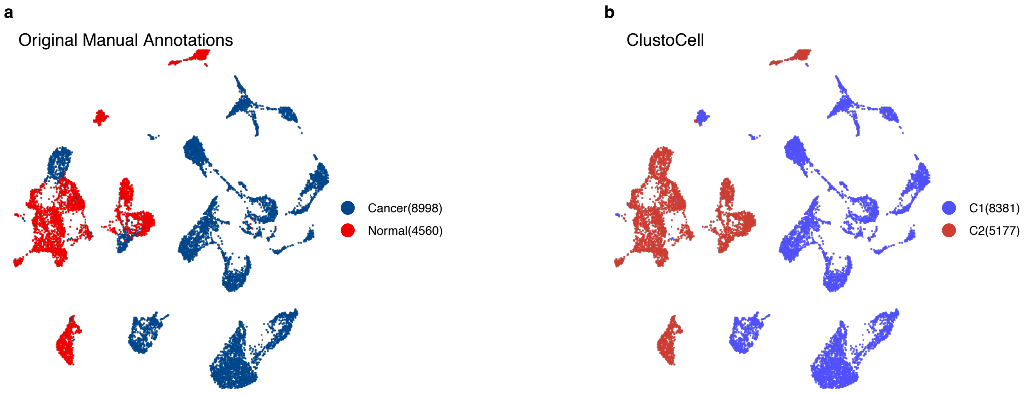

**Supplementary Fig. 14: UMAP Plots of GSE131928 Dataset Colored by Original Manual Annotations and ClustoCell.**
Side-by-side UMAP embeddings of the GSE131928 glioblastoma scRNA-seq dataset showing concordance between expert manual annotations and ClustoCell-derived clusters. **a**, original manual annotations. **b**, ClustoCell clusters.

#### Supplementary Figure 15

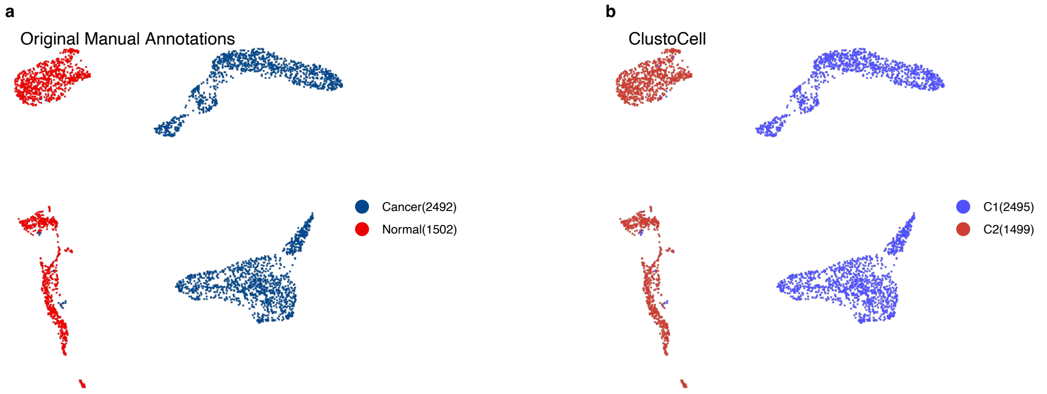

**Supplementary Fig. 15: UMAP Plots of GSE142213 Dataset Colored by Original Manual Annotations and ClustoCell.**
Side-by-side UMAP embeddings of the GSE142213 scRNA-seq dataset showing concordance between expert manual annotations and ClustoCell-derived clusters. **a**, original manual annotations. **b**, ClustoCell clusters.

#### Supplementary Figure 16

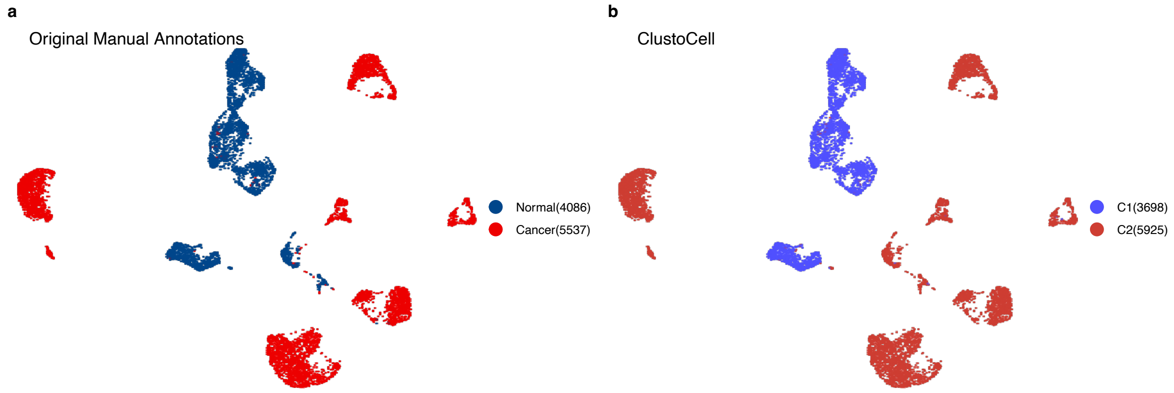

**Supplementary Fig. 16: UMAP Plots of GSE154109 Dataset Colored by Original Manual Annotations and ClustoCell.**
Side-by-side UMAP embeddings of the GSE154109 scRNA-seq dataset showing concordance between expert manual annotations and ClustoCell-derived clusters. **a**, original manual annotations. **b**, ClustoCell clusters.

#### Supplementary Figure 17

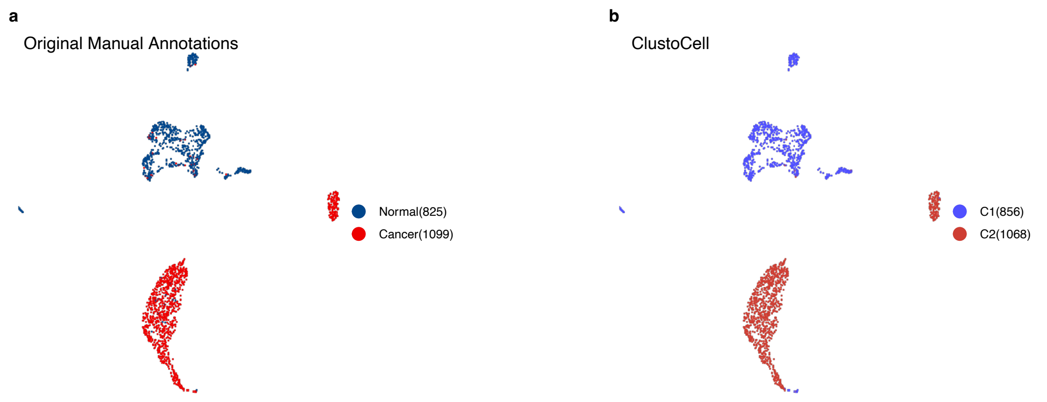

**Supplementary Fig. 17: UMAP Plots of GSE150766 Dataset Colored by Original Manual Annotations and ClustoCell.**
Side-by-side UMAP embeddings of the GSE150766 scRNA-seq dataset showing concordance between expert manual annotations and ClustoCell-derived clusters. **a**, original manual annotations. **b**, ClustoCell clusters.

#### Supplementary Figure 18

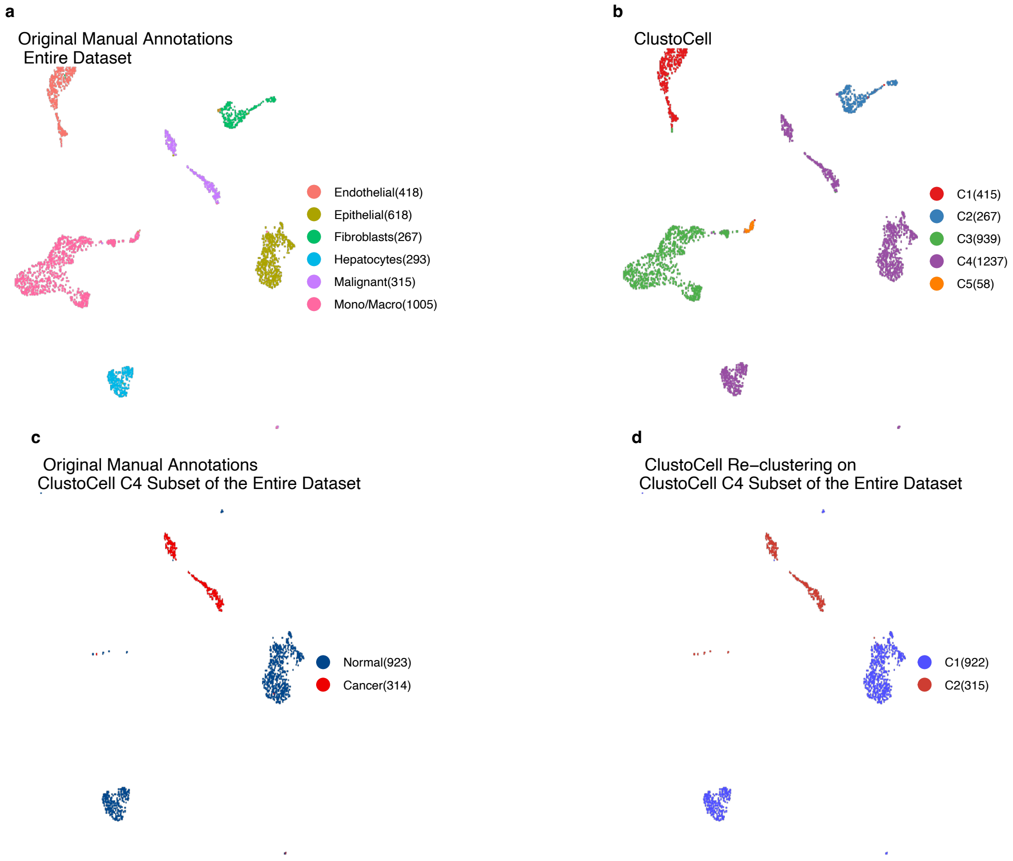

**Supplementary Fig. 18: UMAP Plots of GSE146409 Dataset Colored by Original Manual Annotations and ClustoCell.**

UMAP embeddings of the GSE146409 scRNA-seq dataset, comparing expert manual annotations with ClustoCell clustering at the full-dataset level and after ClustoCell-driven re-clustering of a selected first-level subset (cluster C4). **a**, original manual annotations of the entire dataset. **b**, ClustoCell clusters of the entire dataset. **c**, original manual annotations restricted to the ClustoCell C4 subset. **d**, ClustoCell re-clustering of the same C4 subset, illustrating the within-cluster substructure recovered at higher resolution.

#### Supplementary Figure 19

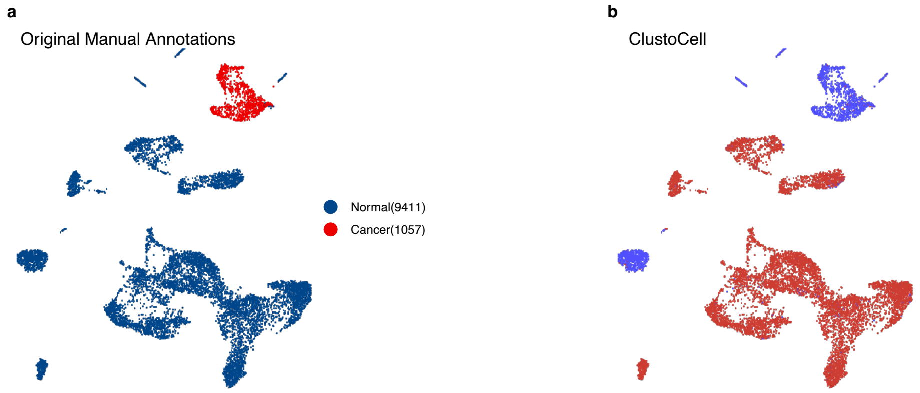

**Supplementary Fig. 19: UMAP Plots of GSE146771 Dataset Colored by Original Manual Annotations and ClustoCell.**
Side-by-side UMAP embeddings of the GSE146771 colorectal cancer scRNA-seq dataset showing concordance between expert manual annotations and ClustoCell-derived clusters. **a**, original manual annotations. **b**, ClustoCell clusters.

#### Supplementary Figure 20

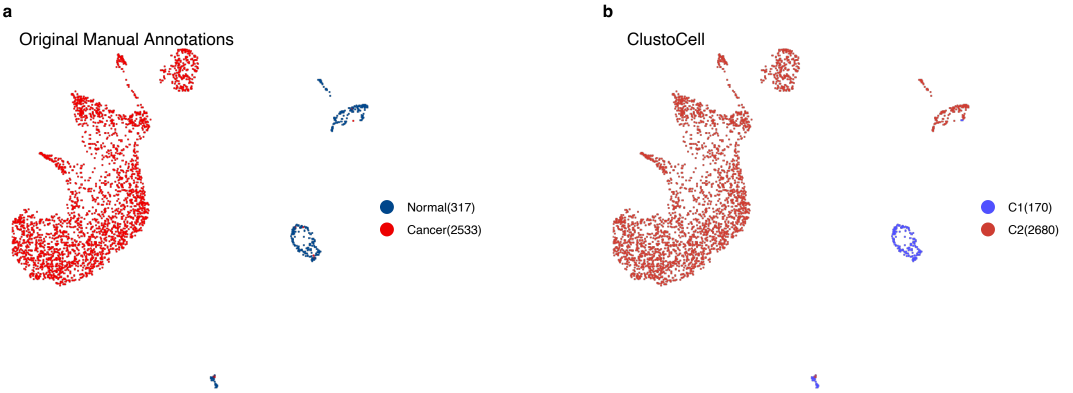

**Supplementary Fig. 20: UMAP Plots of GSE159115 Dataset Colored by Original Manual Annotations and ClustoCell.**
Side-by-side UMAP embeddings of the GSE159115 renal cell carcinoma scRNA-seq dataset showing concordance between expert manual annotations and ClustoCell-derived clusters. **a**, original manual annotations. **b**, ClustoCell clusters.

#### Supplementary Figure 21

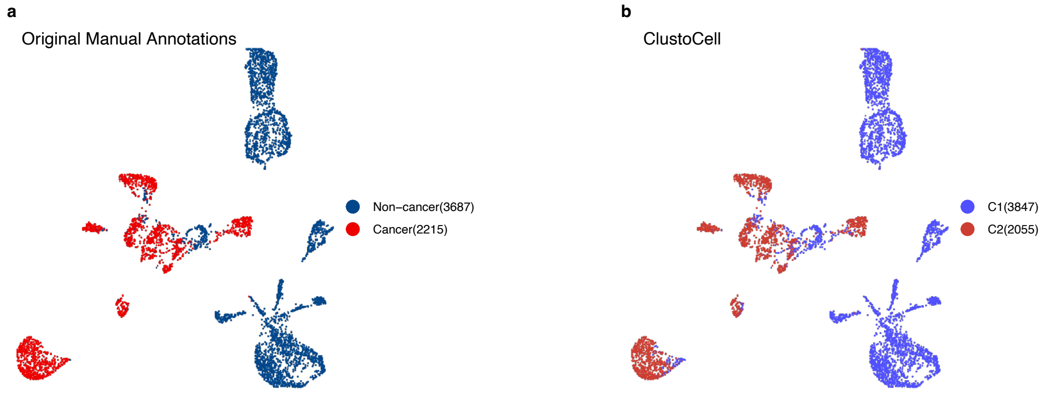

**Supplementary Fig. 21: UMAP Plots of GSE103322 Dataset Colored by Original Manual Annotations and ClustoCell.**
Side-by-side UMAP embeddings of the GSE103322 head and neck squamous cell carcinoma (HNSCC) scRNA-seq dataset showing concordance between expert manual annotations and ClustoCell-derived clusters. **a**, original manual annotations. **b**, ClustoCell clusters.

#### Supplementary Figure 22

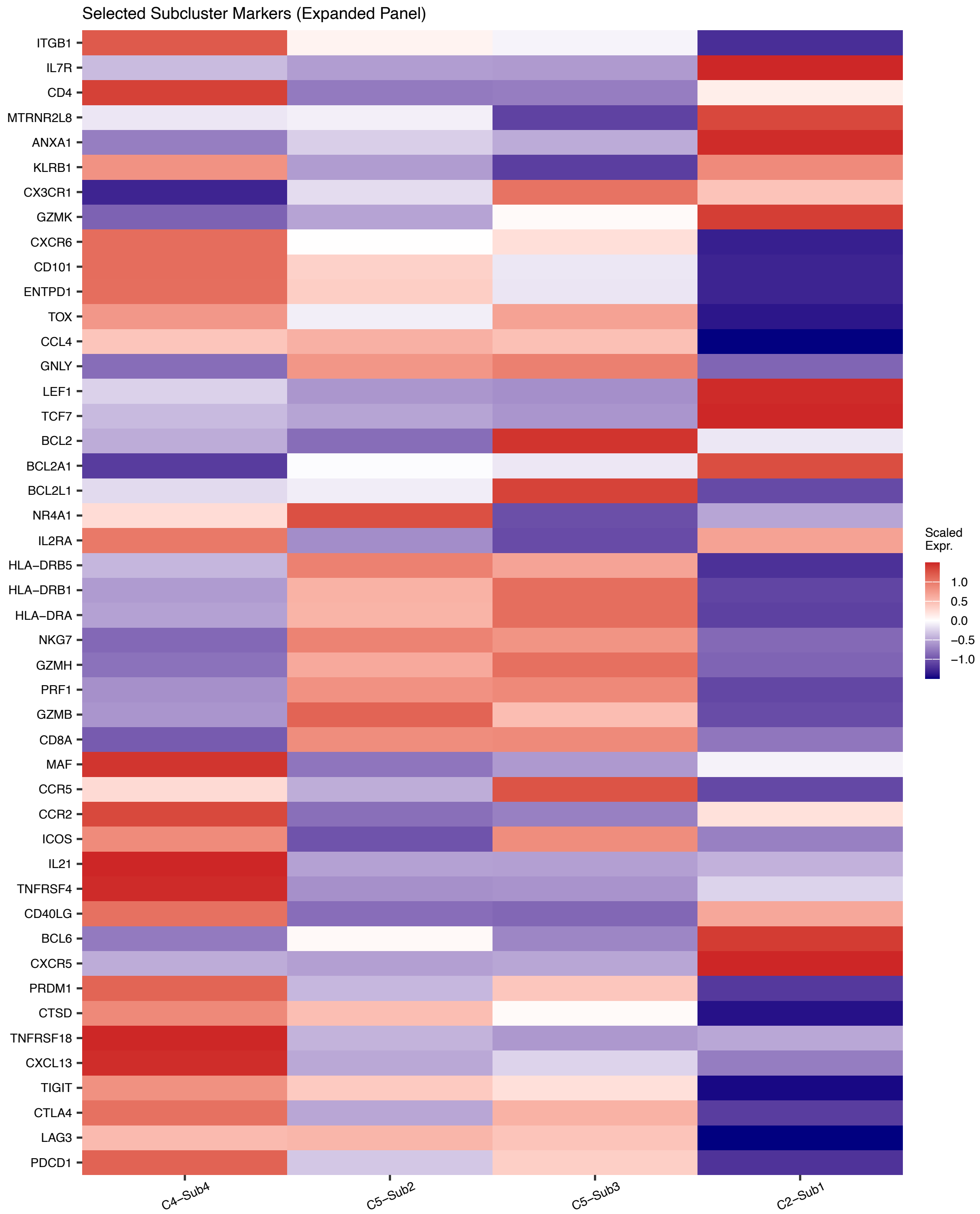

**Supplementary Fig. 22: Expanded canonical-marker panel resolves the cell-state identity of the four expansion-enriched predictive subclusters and supports their refined annotation.**

Scaled expression heatmap of an expanded panel of 46 canonical lineage, activation, and exhaustion-associated genes across the four expansion-enriched predictive ClustoCell subclusters identified in pre-treatment breast cancer biopsies of the Bassez *et al*. cohort: the HLA-DR$^{+}$ NR4A1$^{+}$ activated / early-effector-like exhausted CD8 state (C5-Sub2), the HLA-DR$^{+}$ CX3CR1$^{+}$ effector-like / intermediate exhausted CD8 state (C5-Sub3), the CXCL13$^{+}$ TPH-like CD4 exhausted helper state (C4-Sub4), and the IL7R$^{+}$ ANXA1$^{+}$ GZMK$^{+}$ KLRB1$^{+}$ CD4 effector-memory state (C2-Sub1). The gene panel extends the ClustoCell-derived markers shown in Fig. 6h with canonical references for: inhibitory checkpoints and CD4 exhaustion (*PDCD1*, *LAG3*, *CTLA4*, *TIGIT*, *TNFRSF18*, *PRDM1*, *CXCL13*); cytotoxic effector molecules (*GZMB*, *GZMH*, *PRF1*, *NKG7*, *GNLY*, *CCL4*); class II MHC / interferon licensing (*HLA-DRA*, *HLA-DRB1*, *HLA-DRB5*); stem-like / progenitor exhaustion transcription factors and survival regulators (*TCF7*, *LEF1*, *BCL2*); early TCR-driven activation and costimulation (*NR4A1*, *IL2RA*, *ICOS*, *TNFRSF4*/*OX40*, *BCL2L1*, *BCL2A1*); CD4 helper exhaustion programs distinguishing TPH from GC-TFH (*CXCR5*, *BCL6*, *IL21*, *CD40LG*, *ICOS*, *TNFRSF4*, *MAF*, *CCR2*, *CCR5*); terminal versus intermediate / effector-like exhaustion (*TOX*, *ENTPD1*/*CD39*, *CD101*, *CXCR6*, *GZMK*, *CX3CR1*); and CD4 effector-memory lineage anchors (*CD4*, *IL7R*, *ITGB1*, *ANXA1*, *KLRB1*, *CD40LG*). Mean expression per subcluster was computed on log-normalised counts and each gene was z-scored across only the four subclusters shown; relative enrichment within this panel therefore reflects ordering among these four populations rather than absolute expression across the full T-cell compartment of the dataset. Colour denotes scaled expression (z-score). Jointly, the panel shows that C5-Sub2 is GZMB$^{+}$ NR4A1$^{+}$ HLA-DR$^{+}$ but lacks the canonical TCF7$^{hi}$ LEF1$^{hi}$ BCL2$^{hi}$ stem-like progenitor exhausted (Tpex) signature, supporting its classification as an activated / early-effector-like exhausted CD8 state rather than a progenitor population; C5-Sub3 expresses *TOX* together with *CX3CR1*, *BCL2* and *BCL2L1* but is not selectively enriched for the terminal-exhaustion anchors *ENTPD1*, *CD101* or *CXCR6*, consistent with an intermediate / effector-like exhausted CD8 state rather than a terminally exhausted state; C4-Sub4 co-expresses a CD4 exhaustion program (*PDCD1*, *CTLA4*, *TIGIT*, *TNFRSF18*, *PRDM1*, *ENTPD1*, *TOX*, *CXCL13*) and a helper-output module (*IL21*, *CD40LG*, *TNFRSF4*/*OX40*, *ICOS*), and is further characterized by strong enrichment of the TPH-defining transcription factor *MAF* and chemokine receptor *CCR2* in the absence of the canonical germinal-centre TFH markers *CXCR5* and *BCL6*, supporting a CXCL13$^{+}$ T peripheral helper (TPH)-like CD4 exhausted helper identity rather than a bona fide GC-TFH cell and C2-Sub1 is unambiguously a CD4-compartment effector-memory state with selective enrichment of *IL7R*, *ANXA1*, *GZMK*, *KLRB1* and *CD40LG*, and with *CD8A*, *GZMB*, *PRF1*, *NKG7* and *GNLY* each markedly lower than in the three CD8 / effector-skewed predictive subclusters, indicating that the cluster itself is not confounded by CD8 or NK lineage contamination at single-cell resolution.

### Supplementary Figure 23

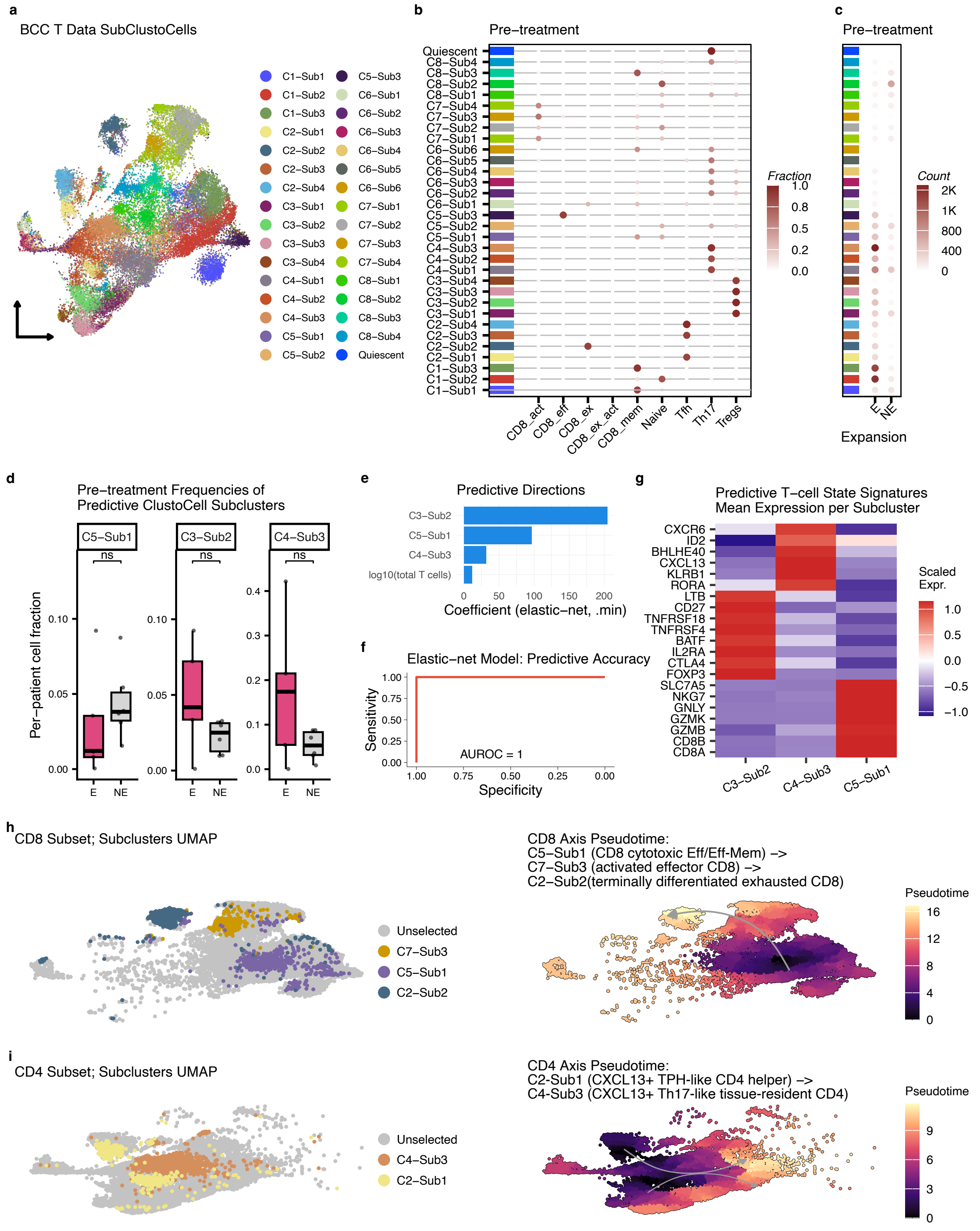

**Supplementary Fig. 23: Expansion-associated immune circuits and lineage trajectories in basal cell carcinoma.**
**a**, UMAP visualization of T cells from the basal cell carcinoma (BCC) dataset from *Yost et al.*, coloured by ClustoCell subclusters. **b**, Concordance between ClustoCell subclusters (rows) and original author-provided annotations (columns) at pre-treatment; each dot represents the fraction of cells from a given subcluster assigned to each annotated category. **c**, Mapping of ClustoCell subclusters to per-cell expansion status at pre-treatment (E = expanding, NE = non-expanding). **d**, Per-patient pre-treatment cell fractions of the three expansion-associated subclusters — C5-Sub1 (CD8 cytotoxic effector/effector-memory-like state), C3-Sub2 (activated effector regulatory T cell (eTreg) state), and C4-Sub3 (a Th17-like tissue-resident CD4 state) — stratified by patient-level expansion outcome (E vs NE). Individual points denote patients. **e**, Elastic-net logistic-regression coefficients for the three selected subclusters and for log10 total T-cell count in the pre-treatment patient-level expansion model. **f**, Receiver operating characteristic (ROC) curve for patient-level prediction of expansion status using the combined pre-treatment fractions of C5-Sub1, C3-Sub2, and C4-Sub3 together with $\log_{10}$ total T-cell count (AUROC = 1.00). **g**, ClustoCell-derived marker panel across the three expansion-associated BCC subclusters (C3-Sub2, C4-Sub3, C5-Sub1). Mean expression per subcluster was computed on log-normalised counts and each gene was z-scored across these three subclusters. Colour denotes scaled expression (z-score). **h**, CD8 compartment pseudotime analysis. Left, UMAP of the CD8 subset coloured by ClustoCell subclusters; right, the same embedding coloured by inferred pseudotime, showing a transcriptional continuum from C5-Sub1 (CD8 cytotoxic effector/effector-memory-like state) through an activated effector CD8 state (C7-Sub3) to a terminally differentiated exhausted CD8 state (C2-Sub2). **i**, CD4 compartment pseudotime analysis. Left, UMAP of the CD4 subset coloured by ClustoCell subclusters; right, the same embedding coloured by inferred pseudotime, showing a transcriptional continuum from a $CXCL13^{+}$ IL-$21^{+}$ $MAF^{+}$ $CD40LG^{+}$ TPH-like CD4 helper state (C2-Sub1) to the $CXCL13^{+}$ Th17-like tissue-resident CD4 state (C4-Sub3).

### Supplementary Figure 24

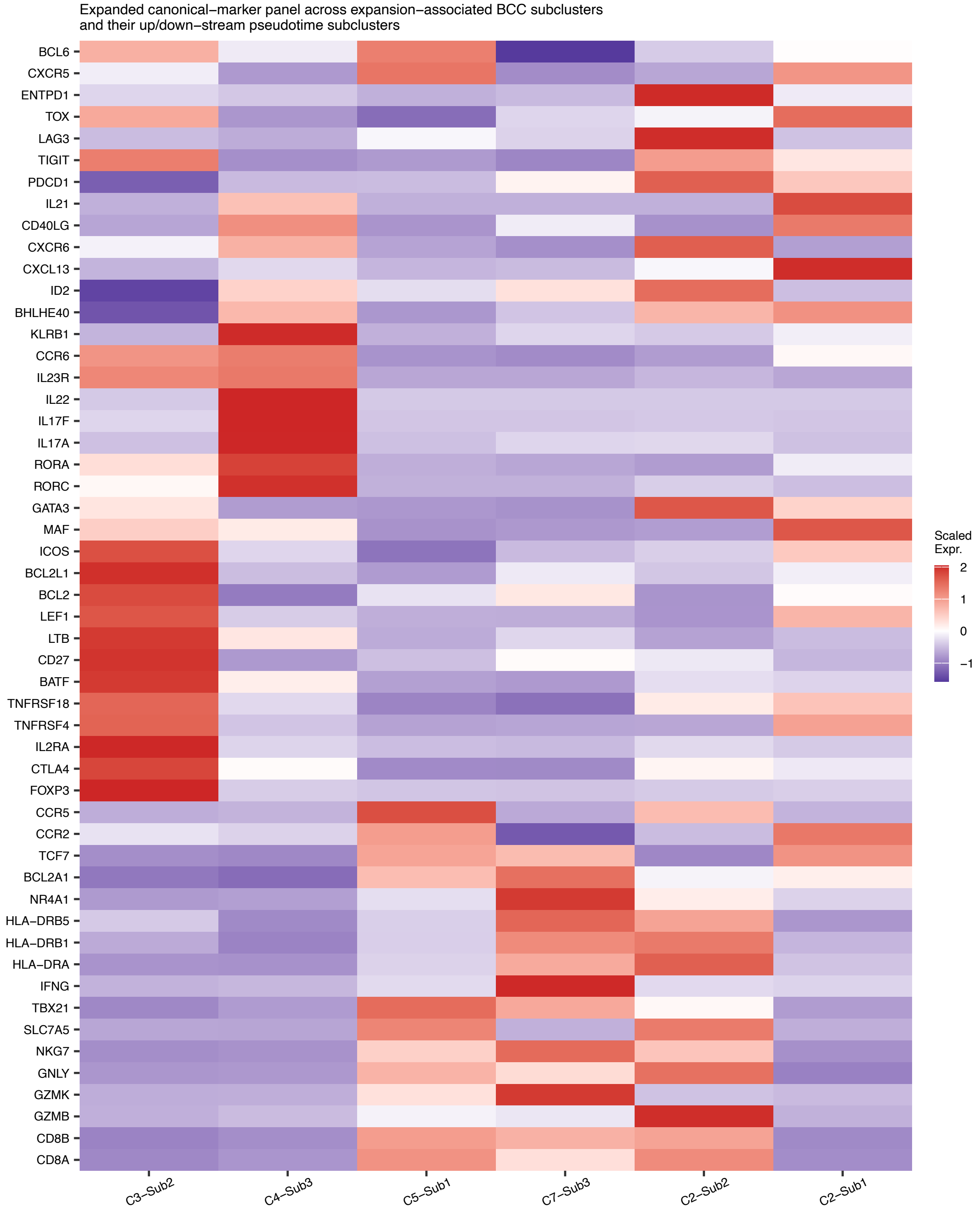

**Supplementary Fig. 24: Expanded canonical-marker panel resolves the cell-state identity of all six annotated BCC T-cell subclusters.**

Mean expression per subcluster was computed on log-normalised counts and each gene was z-scored across all six subclusters (C3-Sub2, C4-Sub3, C5-Sub1, C7-Sub3, C2-Sub2, C2-Sub1). The panel comprises 52 canonical genes covering: CD8 cytotoxic/effector program (*CD8A, CD8B, GZMB, GZMK, GNLY, NKG7, SLC7A5, TBX21, IFNG*); MHC class II and activation (*HLA-DRA, HLA-DRB1, HLA-DRB5, NR4A1*); CD8 effector-memory survival and chemotaxis (*BCL2A1, TCF7, CCR2, CCR5*); inhibitory receptors and exhaustion markers (*PDCD1, TIGIT, LAG3, TOX, ENTPD1*); regulatory T cell core with memory coat (*FOXP3, CTLA4, IL2RA, TNFRSF4/OX40, TNFRSF18/GITR, BATF, CD27, LTB, LEF1, BCL2, BCL2L1, ICOS, MAF, GATA3*); Th17 lineage core (*RORC, RORA, IL17A, IL17F, IL22, IL23R, CCR6, KLRB1, BHLHE40, ID2*); and Th17-like helper/TFH exclusion module (*CXCL13, CXCR6, CD40LG, IL21, CXCR5, BCL6*). The panel confirms the following cell-state identities. C3-Sub2 is an activated effector regulatory T cell (eTreg) state: *FOXP3, CTLA4, IL2RA, TNFRSF4, TNFRSF18, BATF*, and *ICOS* are all near-maximal, a memory/central-memory transcriptional coat (*LEF1, BCL2, BCL2L1, CD27, LTB, MAF*) is co-expressed, and all CD8-effector, Th17, and Th1 lineage markers are depleted. C4-Sub3 is a CXCL13$^+$ IL-17A/F$^+$ RORC$^+$ Th17-like tissue-resident CD4 state: the full Th17 lineage core (*RORC, RORA, IL17A, IL17F, IL22, IL23R, CCR6, KLRB1, BHLHE40, ID2*) and tissue-resident helper module (*CXCL13, CXCR6, CD40LG, IL21*) are near-maximal, while Th1 (*TBX21, IFNG*), Th2 (*GATA3*), Treg (*FOXP3*), GC-TFH (*CXCR5, BCL6*), and TPH (*MAF, CCR2, CCR5*) lineage-defining markers are all depleted. C5-Sub1 is a CD8 cytotoxic effector/effector-memory-like state: *CD8A, CD8B, GZMB, GZMK, GNLY, NKG7, SLC7A5, TBX21, IFNG, HLA-DRA, HLA-DRB1, TCF7, CCR2*, and *CCR5* are near-maximal, and all Treg, Th17, and *Th2* lineage markers are depleted. C7-Sub3 is a GZMK$^+$ HLA-DR$^+$ NR4A1$^+$ activated effector CD8 state: *IFNG, GZMK, NR4A1, HLA-DRB5*, and *NKG7* are near-maximal, *BCL2A1* is elevated, *TCF7* is intermediate, and inhibitory receptors (*PDCD1, LAG3, TIGIT*) are low. C2-Sub2 is a PDCD1$^+$ LAG3$^+$ ENTPD1$^+$ CXCR6$^+$ GZMB$^+$ HLA-DR$^+$ tissue-resident-like terminally differentiated exhausted CD8 state: *PDCD1, LAG3, ENTPD1, CXCR6, GZMB, GNLY, HLA-DRA*, and *HLA-DRB1* are all near-maximal, and *TCF7, LEF1*, and *BCL2* are depleted, consistent with a terminally differentiated phenotype; TOX is not elevated in this cluster (highest in C2-Sub1), distinguishing this state from the canonical TOX-driven terminal-exhaustion program and supporting the descriptor "terminally differentiated" over "terminally exhausted". C2-Sub1 is a CXCL13$^+$ IL-21$^+$ MAF$^+$ CD40LG$^+$ TPH-like CD4 helper state: *CXCL13, IL21, MAF, CCR2, CD40LG, TCF7*, and *CXCR5* are near-maximal, *BCL6* is near zero (excluding canonical GC-TFH commitment), and all CD8-effector, Th17, and Treg markers are depleted; the CXCR5-intermediate/BCL6-negative profile distinguishes this state from committed GC-TFH cells while the CXCL13$^+$ MAF$^+$ CCR2$^+$ signature supports a TPH-like identity. Colour denotes scaled expression (z-score).

#### Supplementary Figure 25

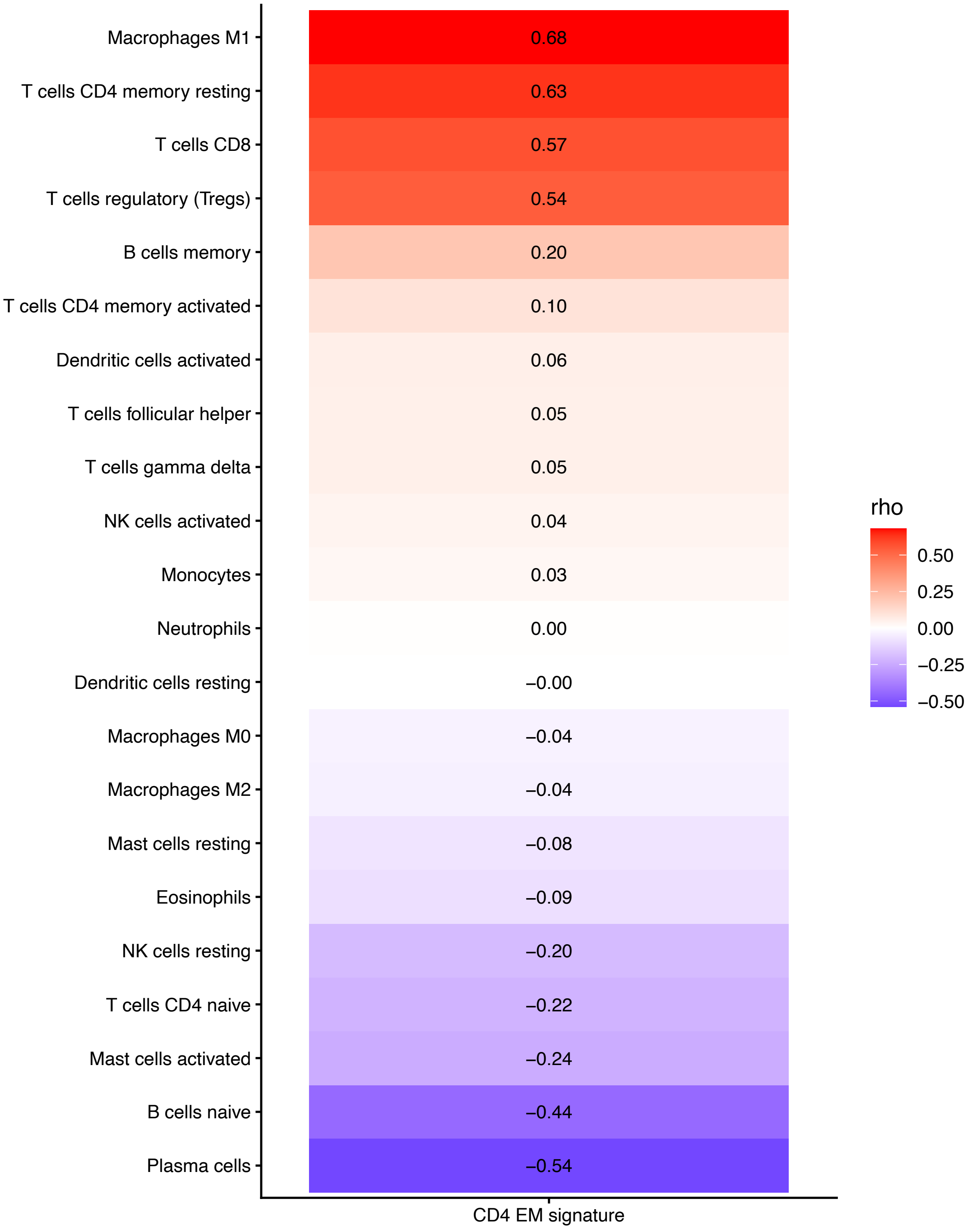

**Supplementary Fig. 25: Correlation between the C2-Sub1 CD4 effector-memory transcriptional program and CIBERSORT-inferred immune cell populations in TCGA breast cancer**

Spearman correlations were computed between the C2-Sub1 Singscore-derived gene-set score and relative abundances of immune cell populations estimated using the CIBERSORT LM22 signature matrix. Positive correlations with CD8 T cells, resting memory CD4 T cells and M1 macrophages indicate that the prognostic C2-Sub1 program is associated with a broader immune-inflamed tumour microenvironment in bulk RNA-seq samples.
